# Translating GWAS-identified loci for cardiac rhythm and rate using an *in vivo* image- and CRISPR/Cas9-based approach

**DOI:** 10.1101/385500

**Authors:** Benedikt von der Heyde, Anastasia Emmanouilidou, Eugenia Mazzaferro, Silvia Vicenzi, Ida Höijer, Tiffany Klingström, Sitaf Jumaa, Olga Dethlefsen, Harold Snieder, Eco de Geus, Adam Ameur, Erik Ingelsson, Amin Allalou, Hannah L. Brooke, Marcel den Hoed

## Abstract

A meta-analysis of genome-wide association studies (GWAS) identified eight loci that are associated with heart rate variability (HRV), but candidate genes in these loci remain uncharacterized. We developed an image- and CRISPR/Cas9-based pipeline to systematically characterize candidate genes for HRV in live zebrafish embryos. Nine zebrafish orthologues of six human candidate genes were targeted simultaneously in eggs from fish that transgenically express GFP on smooth muscle cells (Tg[*acta2:GFP*]), to visualize the beating heart. An automated analysis of repeated 30s recordings of beating atria in 381 live, intact zebrafish embryos at 2 and 5 days post-fertilization highlighted genes that influence HRV (*hcn4* and *si:dkey-65j6.2 [KIAA1755]*); heart rate (*rgs6 and hcn4*); and the risk of sinoatrial pauses and arrests (*hcn4*). Exposure to 10 or 25µM ivabradine – an open channel blocker of HCNs – for 24h resulted in a dose-dependent higher HRV and lower heart rate at 5 days post-fertilization. Hence, our screen confirmed the role of established genes for heart rate and rhythm (*RGS6* and *HCN4*); showed that ivabradine reduces heart rate and increases HRV in zebrafish embryos, as it does in humans; and highlighted a novel gene that plays a role in HRV (*KIAA1755*).

## Introduction

Heart rate variability (HRV) reflects the inter-beat variation of the RR interval. HRV is controlled by the sinoatrial node, which receives input from the autonomic nervous system. Autonomic imbalance has been associated with work stress and other modifiable or non-modifiable risk factors^1^, and is reflected in lower HRV. Lower HRV has been associated with higher cardiac morbidity and mortality^2^, as well as with a higher risk of all-cause mortality^3^. HRV can be quantified non-invasively using an ECG, making HRV a useful clinical marker for perturbations of the autonomic nervous system. However, the mechanisms influencing HRV remain poorly understood.

Recently, we and others identified the first loci that are robustly associated with HRV, using a meta-analysis of genome-wide association studies (GWAS) with data from 53,174 participants^3^. Five of the identified loci had previously been associated with resting heart rate^4^. The heart rate associated loci in turn are associated with altered cardiac conduction and risk of sick sinus syndrome^5^. *In silico* functional annotation of the five loci that are associated with both HRV^3^ and heart rate^4^ resulted in the prioritization of six candidate genes^3^. Functional follow-up experiments - ideally *in vivo* - are required to conclude if these genes are indeed causal, and examine if they influence HRV, heart rate, and/or cardiac conduction.

Mouse models are commonly used in cardiac research, but adult mice show substantial differences in cardiac rate and electrophysiology when compared with humans^5^. Inherently, these differences complicate extrapolation of results from mouse models to humans^5^. Additionally, rodents are not suitable for high-throughput, *in vivo* characterization of cardiac rhythm and rate. Such screens are essential to systematically characterize positional candidate genes in the large number of loci that have now been identified in GWAS for HRV^3^, heart rate^6^ and cardiac conduction^7,8^. Hence, novel model systems that facilitate systematic, *in vivo* characterization of a large number of candidate genes are needed.

In recent years, the zebrafish has become an important model system for genetic and drug screens for human diseases^9,10^. Despite morphological differences to the human heart, zebrafish have a functional, beating heart at ∼24 hours post-fertilization, and the ECG of the two-chambered adult zebrafish heart is similar to that of humans^11^. Conditions affecting cardiac electrophysiology have previously been successfully modeled in zebrafish. For example, *nkx2.5* is necessary for normal HRV^12^, mutations in *kcnh2* have been used to model long QT syndrome^13^, *trpm7* was shown to influence heart rate and risk for sinoatrial pauses^14^, and knockdown of *hcn4* has been used to model human sick sinus syndrome in zebrafish^15^. Fluorescently labeled transgenes facilitate visualization of cell types and tissues of interest, which can now be accomplished in high throughput thanks to advances in automated positioning of non-embedded, live zebrafish embryos^16–18^. In addition, the zebrafish has a well-annotated genome, with orthologues of at least 71.4% of human genes^19^.

These genes can be targeted efficiently and in a multiplexed manner using Clustered, Regulatory Interspaced, Short Palindromic Repeats (CRISPR) and CRISPR-associated systems (Cas)^20^. All characteristics combined make zebrafish embryos an attractive model system to systematically characterize candidate genes for cardiac rhythm, rate and conduction.

The aim of this study was to objectively characterize the most promising candidate genes in loci associated with both HRV and heart rate for a role in cardiac rhythm, rate and function, using a large-scale, image-based screen in zebrafish embryos.

## Results

### Experimental pipeline

Our experimental pipeline is outlined in **Fig 1 and Supp Fig 1**. CRISPR/Cas9 founders (F_0_) were generated in which all nine zebrafish orthologues of six human candidate genes were targeted simultaneously using a multiplexed approach (**Table 1**)^20^. We used a background with transgenically expressed, fluorescently labelled smooth muscle cells (Tg[*acta2:GFP*])^21^, to visualize the beating heart. Founders were raised to adulthood and in-crossed. On the morning of day 2 post fertilization (dpf), F_1_ embryos were individually anesthetized, to minimize exposure to tricaine and to standardize the imaging procedure. We then captured bright field images of whole embryos in 12 orientations to quantify body size, and recorded the beating atrium for 30s in 383 live, intact embryos, at 152 frames/s using an automated positioning system, fluorescence microscope and CCD camera (see Methods). After image acquisition, embryos were washed, dispensed in 96-well plates, and placed in an incubator at 28.5°C, until 5dpf. Of the 383 embryos imaged at 2dpf, 326 were imaged again on the morning of day 5 (**Supp Fig 2**). Importantly, only three embryos died between 2 and 5dpf. This repeated measures design allowed us to quantify genetic effects on cardiac outcomes and body size at two key stages of early cardiac development^22^. A custom-written MatLab script was used to automatically quantify HRV and heart rate from the recordings acquired at 2 and 5dpf (see Methods). After imaging at 5dpf, embryos were sacrificed and sequenced at the CRISPR/Cas9-targeted sites, followed by alignment; quality control; variant calling; variant annotation; and statistical analysis.

**Figure 1:**
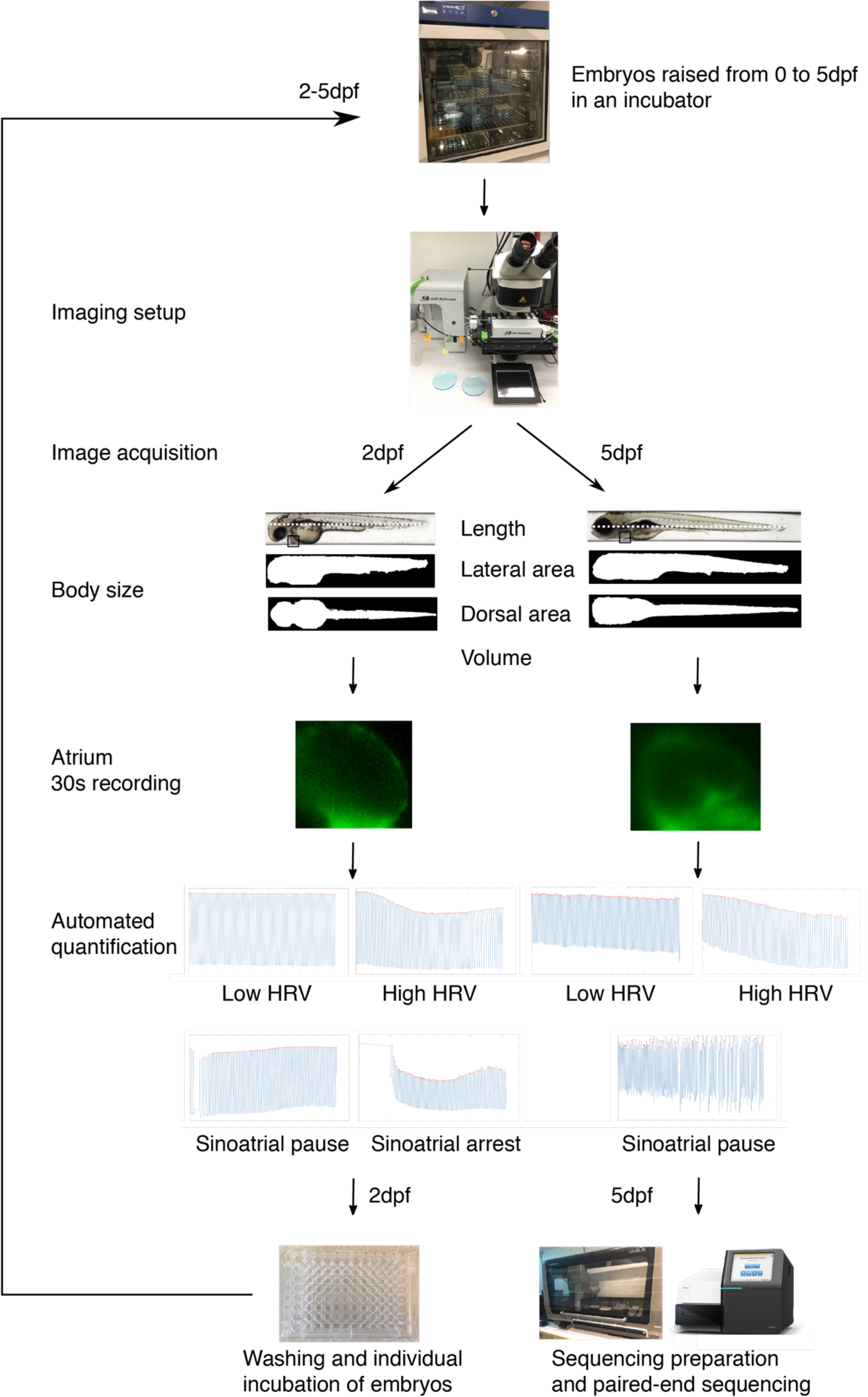
Overview of the experimental pipeline showing the image-based acquisition and post-processing of images and samples. dpf = days post-fertilization.

**Figure 2:**
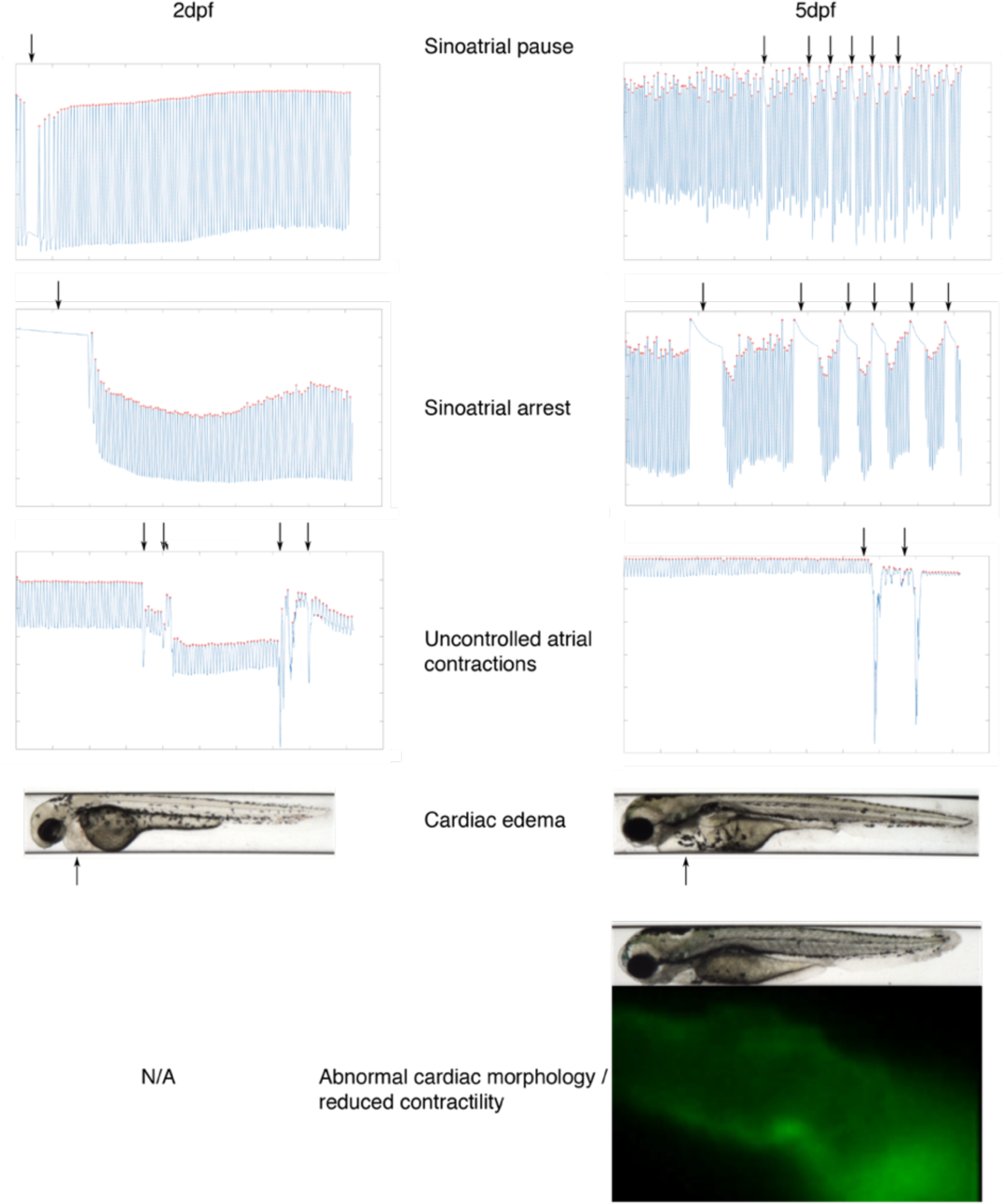
Visualization of cardiac rhythmic or other abnormalities, highlighted by arrows. Sinoatrial pauses were defined as the atrium ceasing to contract for >3x the median inter-beat interval of the embryo. A sinoatrial arrest was defined as an event where the atrium stopped contracting for >2s. Abnormal cardiac morphology was defined as the atrium appearing as a tube-like structure, and impaired cardiac contractility was defined as the atrium vibrating rather than contracting.

**Table 1:**
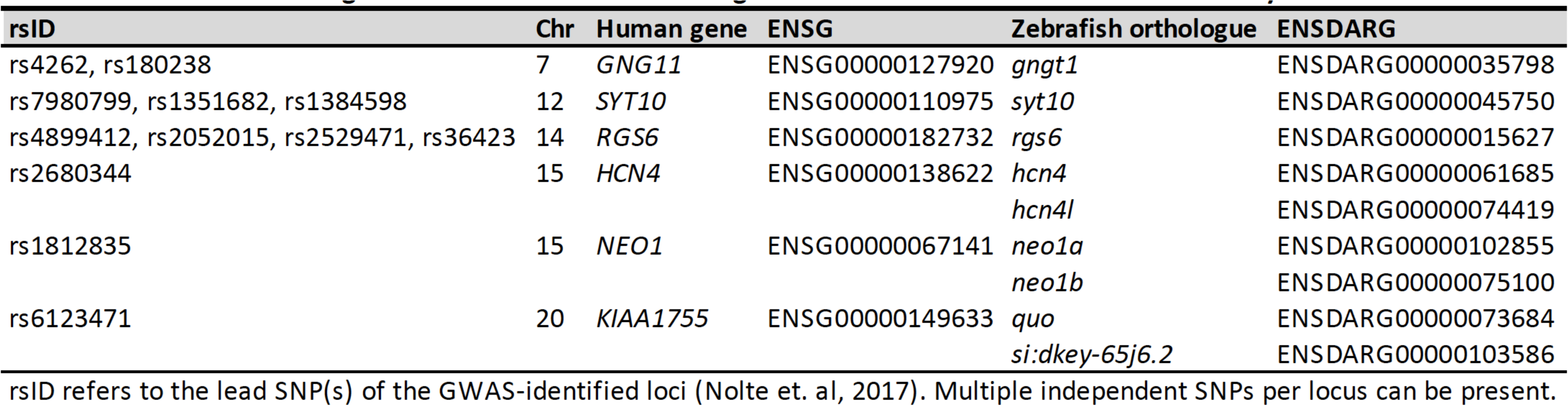
Selected candidate genes and their zebrafish orthologues in GWAS-identified heart rate variability associated loci

### Descriptive results

At 2dpf, some embryos showed sinoatrial pauses (n=39, 10.3%**, Supp recording 1, Fig 2**) and arrests (n=36, 9.5%, **Supp recording 2, Fig 2**); cardiac edema (n=16, 4.2%); or uncontrolled atrial contractions (n=1, 0.2%, **Supp recording 3, Fig 2**). At 5dpf, fewer embryos displayed sinoatrial pauses (n=9, 2.7%, only one of which also showed a sinoatrial pause at 2dpf); sinoatrial arrests (n=3, 0.9%, none of which showed a sinoatrial arrest at 2dpf); and cardiac edema (n=15, 4.5%, nine of which already had cardiac edema at 2dpf); while uncontrolled atrial contractions and an abnormal cardiac morphology (**Supp recording 4, Fig 2**) were first observed in some embryos at 5dpf (n=9, 2.7% and n=6, 1.8%, especially). For embryos free from cardiac abnormalities, distributions of HRV and heart rate at 2 and 5dpf are shown in **Supp Fig 3**. Distributions for body size are shown in **Supp Fig 4**.

**Figure 3:**
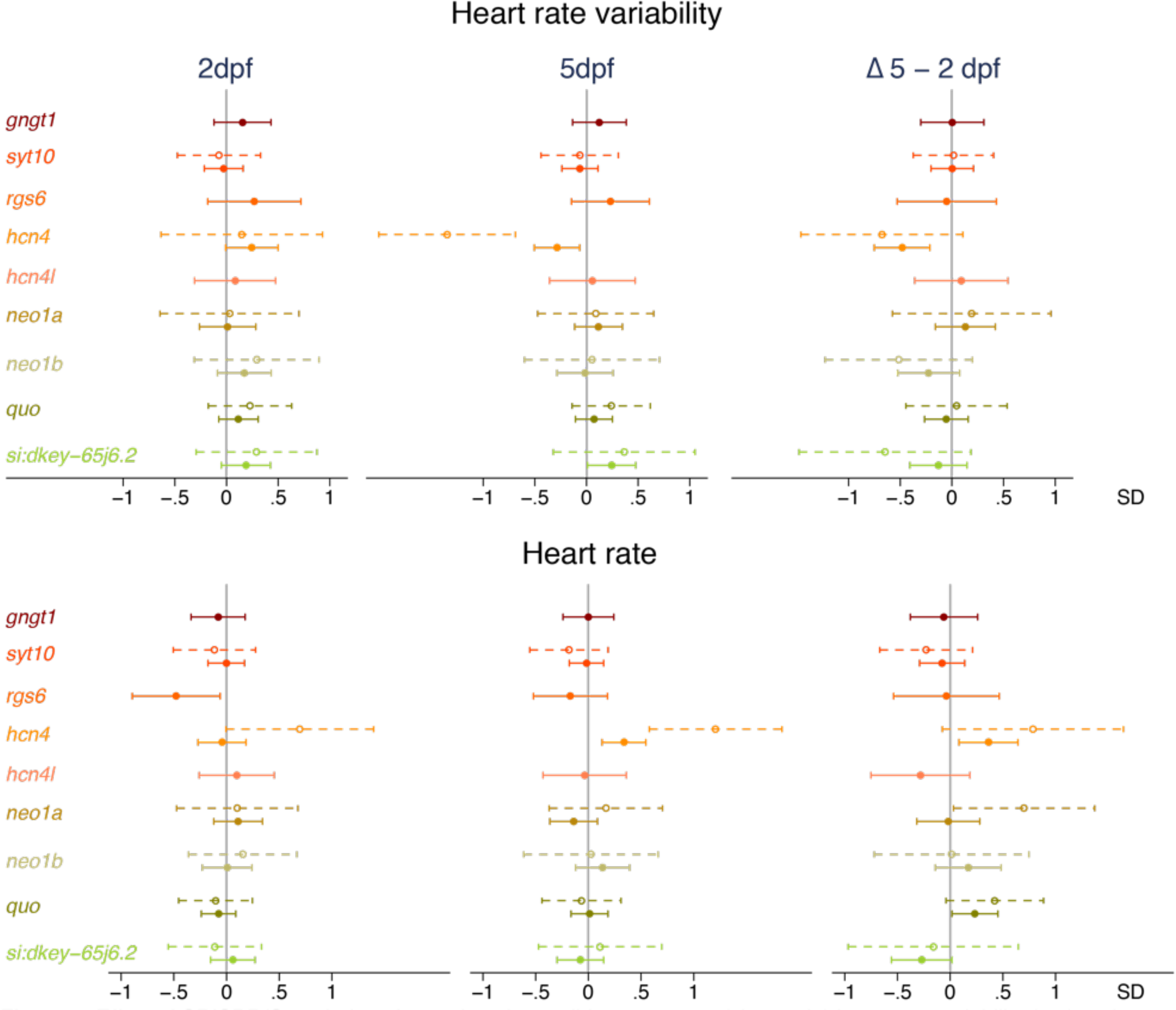
Effect of CRISPR/Cas9-induced mutations in candidate genes on (change in) heart rate variability (top) and heart rate (bottom) at 2 days post-fertilization (dpf, n=234) and 5dpf (n=285). Full dots and solid whiskers show the effect size and 95% confidence interval (CI) for each additional mutated allele, weighted by the mutation’s predicted effect on protein function. Open dots and dotted whiskers indicate the effect and 95% CI for nonsense mutations in both alleles vs. no CRISPR/Cas9-induced mutations. Effects were adjusted for the weighted number of mutated alleles in the other targeted genes, as well as for time of day (fixed factors), with embryos nested in batches (random factor). *quo* and *si:dkey-65j6.2* are orthologues of the human *KIAA1755*.

**Figure 4:**
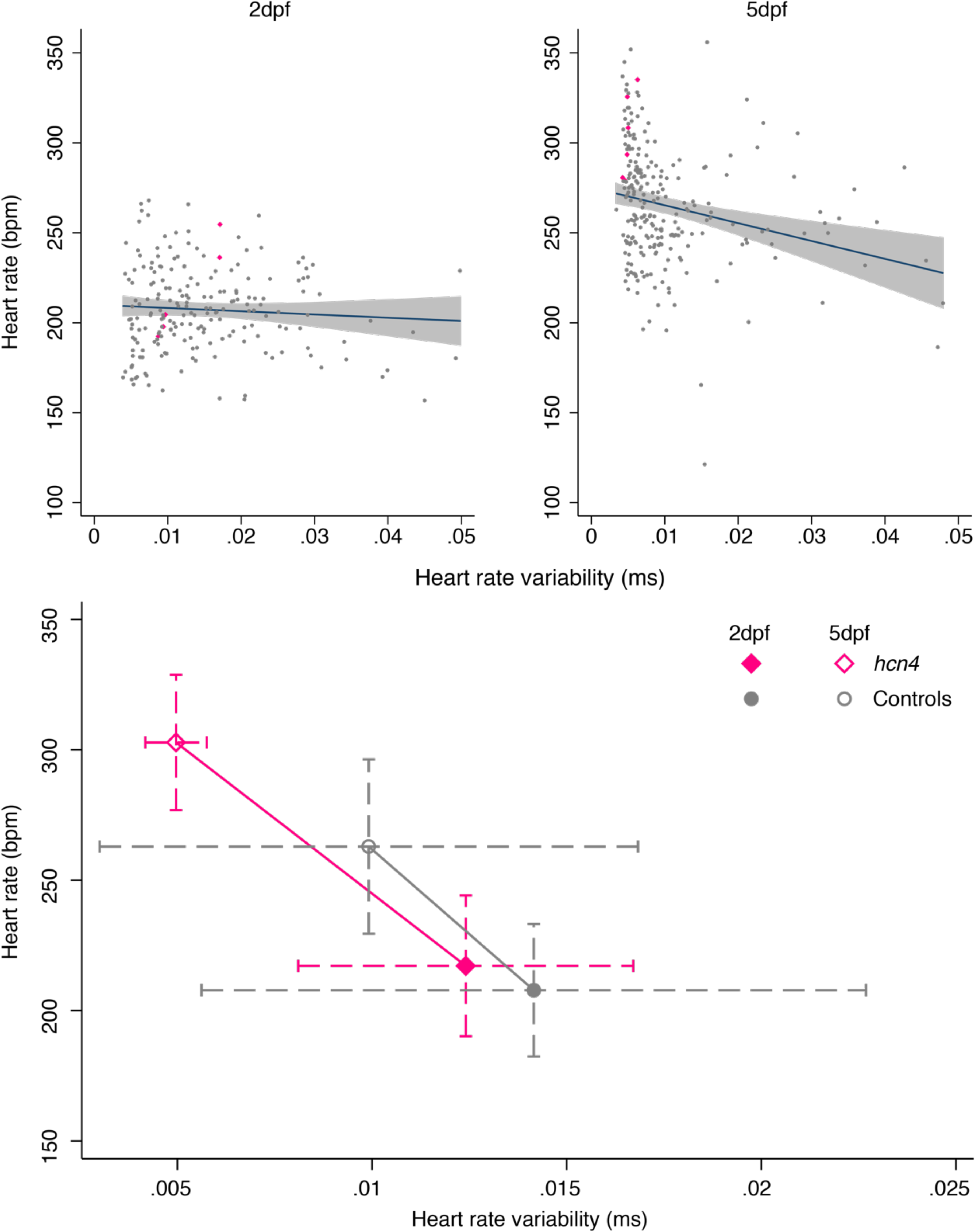
Association fo heart rate variability and heart rate at 2 and 5 days post-fertilization (dpf) and effect of nonsense mutations in both *hcn4* alleles. Top: the association of heart rate variability and heart rate at 2 and 5dpf, with embryos carrying nonsense mutations in both *hcn4* alleles shown as pink diamonds. Bottom: the mean + standard deviation of heart rate variability and heart rate at 2dpf and 5dpf for embryos with nonsense mutations in both *hcn4* alleles (n=5) vs. embryos without CRISPR/Cas9-induced mutations in *hcn4* (n=147).

After imaging at 5dpf, all 383 embryos were sequenced at the nine CRISPR/Cas9 targeted sites (**Table 1, Supp Fig 5**). Transcript-specific dosages were calculated by weighing the number of mutated alleles by the mutations’ predicted impact on protein function, based on Ensembl’s variant effect predictor (VEP). A total of 123 unique alleles were identified across the nine CRISPR/Cas9-targeted sites (**Supp Fig 5**), in which 169 unique mutations were called (**Supp Table 3**), ranging from three unique mutations in *hcn4l* to 34 in *si:dkey-65j6.2* (one of two *KIAA1755* orthologues). Frameshift inducing mutations were most common (47.9%), followed by missense variants (25.4%), in-frame deletions (14.2%), and synonymous variants (5.9%). Eighty-seven, seventy-two and ten mutations were predicted to have a high, moderate, and low impact on protein function, respectively (**Supp Table 3**). Two embryos with missed calls in more than two targeted sites were excluded from the analysis. In the remaining 381 embryos, sequencing call rate was >98% for all but one targeted site, and missed calls were imputed to the mean. Seventy-eight embryos had a missed call for *neo1b* (**Supp Table 4**). Since these 78 embryos showed a similar distribution for all outcomes compared with the remaining embryos (not shown), and since the mutant allele frequency of *neo1b* was 93.4% for the successfully sequenced embryos, missing *neo1b* calls were also imputed to the mean.

**Figure 5:**
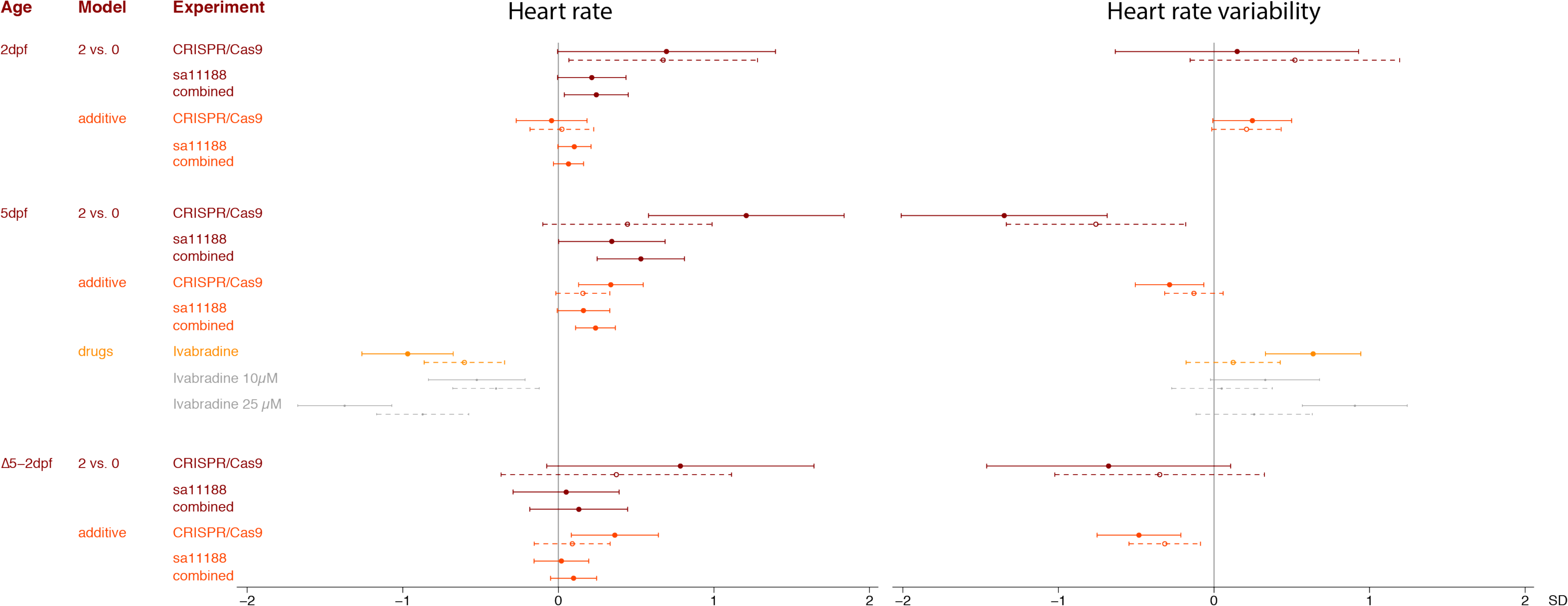
The effect of mutations in *hcn4* and treatment with ivabradine on (change in) heart rate variability and heart rate. Effects were examined by comparing embryos with nonsense mutations in both *hcn4* alleles and embryos free from CRISPR/Cas9 or sa11188 mutations; as well as using an additive model, weighing the number of mutated alleles by their predicted effect on protein function based on Ensembl’s Variant Effect Predictor (VEP). Effects were adjusted for time of day and the weighted number of mutated alleles in other targeted genes as fixed factors and with larvae nested in batches and experiment (for the combined analysis, random factor). Dots and whiskers represent effect sizes and 95% confidence intervals.

We observed a normal distribution of the number of mutated alleles across the nine targeted orthologues (range 2-14 mutated alleles, **Supp Fig 6**). Mutant allele frequencies – based on previously unreported variants located within ±30bp of the CRISPR/Cas9 cut sites - ranged from 4.1% for *hcn4l* to 93.4% for *neo1b* (**Supp Table 4**). We did not identify any mutations at three predicted off-target sites in the 381 successfully sequenced embryos (**Supp Table 2**).

**Figure 6:**
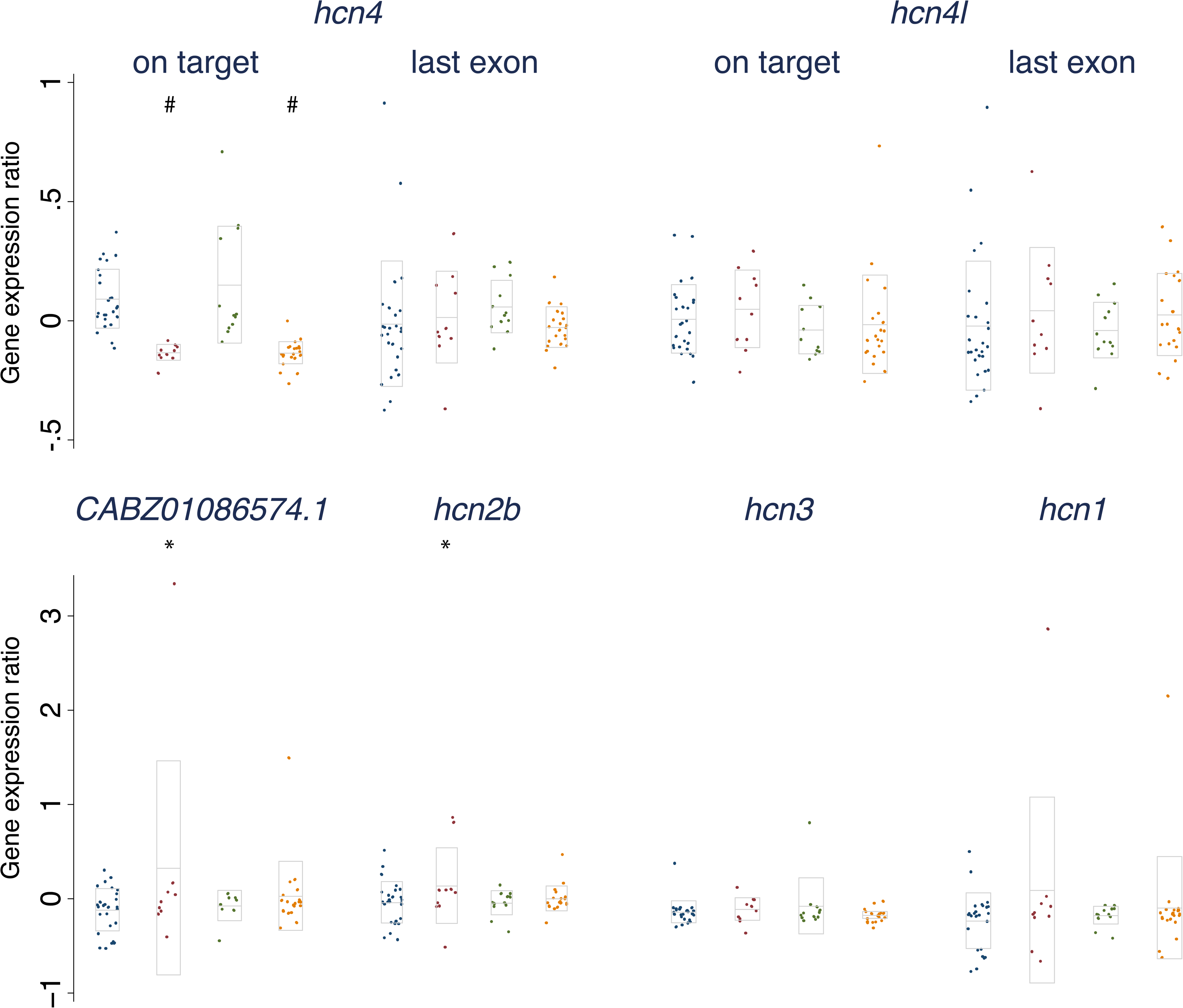
qRT-PCR results for the expression of transcripts with high (>75%) sequence similarity to the main zebrafish *hcn4* transcript, with and without CRISPR/Cas9 targeting of *hcn4*, *hcn4l,* or *hcn4* & *hcn4l*. Each sample consists of five pooled, 5 day-old embryos, which at the single cell stage had been injected with: 1) *hcn4* and *hcn4l* gRNAs, or Cas9 mRNA (controls, in blue, n=26); or with Cas9 mRNA together with 2) *hcn4* gRNA (red, n=10); 3) *hcn4l* gRNA (green, n=12); or 4) *hcn4* and *hcn4l* gRNA (orange, n=21). In all samples, technological triplicates of quantification cycles (Cq) were averaged, and normalized using expression in uninjected controls as calibrator, and expression of *mob4* as a reference gene, using the Pfaffl method. For *hcn4* and *hcn4l*, expression was quantified at the CRISPR/Cas9 target site and at the last exon. Differences between the three experimental conditions and controls were examined in a multiple linear regression analysis, adjusting for batch (n=2). *CABZ01086574.1* is likely an orthologue of the human *HCN2*. Significant differences with controls are highlighted by * (P<0.05) or # (P<1×10^-4^). Boxes show means ± 1 SD. Genes were ordered by sequence similarity to the main *hcn4* transcript.

**Table 2:**
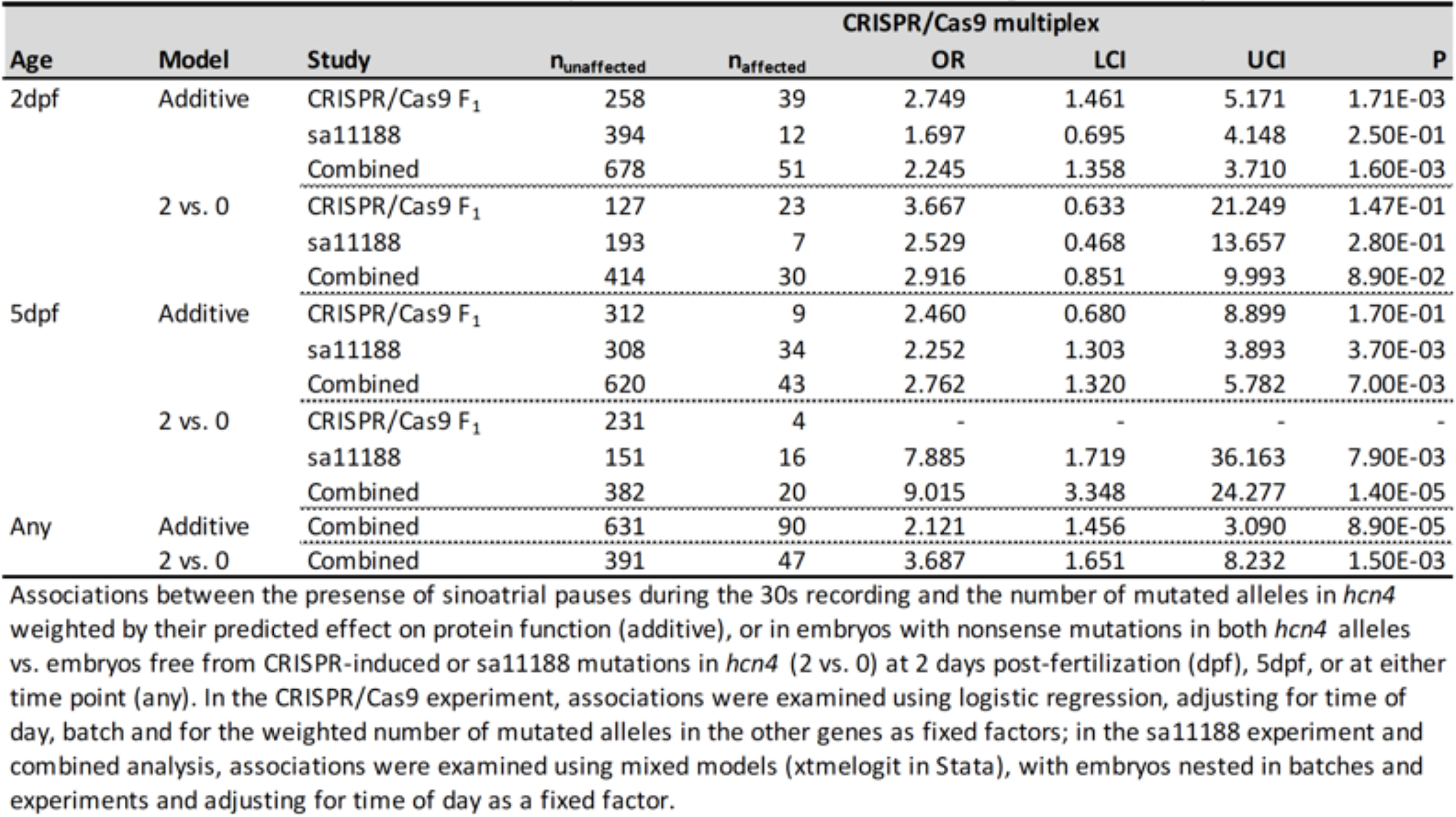
Effect of mutations in *hcn4* on sinoatrial pauses in data from the CRISPR/Cas9 F_1_ and sa11188 experiments combined

### Effects of mutations in candidate genes on sinoatrial pauses and arrests

At 2dpf, each additional CRISPR/Cas9 mutated allele in the first exon of *hcn4* resulted in a >2.5-fold higher odds of sinoatrial pauses or arrests (**Table 2, Supp Tables 5-6**). Embryos showing pauses or arrests during positioning and orienting in the microscope’s field of view, but not during image acquisition were not included in the statistical analysis. Still, it is worth noting that pauses were observed during positioning in four of the nine embryos that later turned out to be compound heterozygous for CRISPR/Cas9-induced nonsense mutations in *hcn4*; and in four of the eight embryos carrying a mutated allele in *hcn4* as well as in *hcn4l*. Importantly, zebrafish embryos can survive without a functionally beating heart at this stage of development, thanks to adequate tissue oxygenation by diffusion^23^. This allowed us to observe genetically driven sinoatrial arrests that would have been lethal in embryos of most other species.

Of the 39 embryos with a sinoatrial pause during the acquisition at 2dpf; two embryos died before imaging at 5dpf, and only one also showed a pause or arrest at 5dpf. Since only six new pauses were observed at 5dpf, the statistical power to detect genetic effects on sinoatrial pauses was low at 5dpf. Earlier we performed a pilot experiment in 406 2dpf offspring of an in-cross of heterozygous carriers of the sa11188 mutation^24^, a nonsense mutation in exon 2 of *hcn4*. Of these 406 embryos, 95, 206 and 105 carried nonsense mutations in 0, 1 and 2 *hcn4* alleles at 2dpf, respectively. Of the 406 embryos, 349 passed quality control at 5dpf, or which 81, 177 and 91 embryos carried 0, 1 and 2 mutated alleles, respectively. The statistical power to detect effects of nonsense mutations in *hcn4* on sinoatrial pauses was thus substantially higher in the pilot study. Combining data from the CRISPR/Cas9 and sa11188 experiments provided us with 51 and 43 sinoatrial pauses at 2 and 5dpf, and confirmed an >2-fold higher odds of sinoatrial pauses for each additional mutated *hcn4* allele at 2 and at 5dpf (**Table 2**). Embryos carrying nonsense mutations in both *hcn4* alleles even had 9-fold higher odds of sinoatrial pauses at 5dpf than embryos free from CRISPR/Cas9-induced and sa11188 mutations in *hcn4* (P=1.4×10^-5^, **Table 2**). Genotype distributions did not deviate from Hardy Weinberg equilibrium in either the CRISPR/Cas9 or sa11188 experiment, confirming that nonsense mutations in *hcn4* were not lethal between 2 and 5dpf.

Of the remaining candidate genes examined in the multiplexed CRISPR/Cas9 experiment, only mutations in *syt10* showed a trend for higher odds of sinoatrial pauses at 2dpf (**Supp Table 6**). This is likely a transient effect, since the six embryos with nonsense mutations in both *syt10* alleles that showed a sinoatrial pause at 2dpf did not die between 2 and 5dpf and did not show a sinoatrial pause at 5dpf. In fact, each additional mutated allele in *syt10* tended to *protect* from sinoatrial pauses at 5dpf; none of the 41 embryos with nonsense mutations in both *syt10* alleles showed a sinoatrial pause at 5dpf (**Supp Table 5**).

### Effects of mutations in candidate genes on heart rate variability and heart rate

We next examined the effect of CRISPR/Cas9-induced mutations on HRV and heart rate in embryos free from sinoatrial pauses and arrests. Embryos with CRISPR/Cas9-induced mutations in *hcn4* tended to have a higher HRV and a higher heart rate at 2dpf (**Fig 3, Supp Fig 3, Supp Tables 7-8)**. From 2 to 5dpf, embryos with CRISPR/Cas9-induced mutations in *hcn4* on average had a larger decrease in HRV and a larger increase in heart rate, resulting in a lower HRV and a higher heart rate at 5dpf (**Figs 3-5, Supp Fig 3, Supp Tables 7-8**). At 5dpf, standardized effect sizes of mutations in *hcn4* on HRV and heart rate were of similar magnitude but opposite direction (**Fig 5, Supp Tables 7-8**). For heart rate, effect estimates were directionally consistent across the CRISPR/Cas9 and sa11188 experiments, but were more conservative in the latter, both at 2 and at 5dpf (**Fig 5**). A lower frame rate in the sa11188 experiment (i.e. 20 frames/s) meant we could not examine the effect of sa11188 mutations in *hcn4* on HRV in the pilot study.

Across the remaining candidate genes, the 17 embryos with one CRISPR/Cas9 mutated *rgs6* allele on average had a nearly 0.5 SD units lower heart rate. The effect size for HRV was ∼2-fold lower and did not reach significance (**Fig 3, Supp Fig 3, Supp Tables 4, 7**). Furthermore, embryos with mutations in *si:dkey-65j6.2* (i.e. *KIAA1755*) tended to have: 1) a higher HRV at 2dpf; 2) a larger decrease in HRV from 2 to 5dpf; and 3) a higher HRV at 5dpf (**Fig 3, Supp Fig 3, Supp Table 7**). While we did not observe an effect of mutations in *si:dkey-65j6.2* or *quo* on heart rate at 2 or 5dpf, embryos with mutations in *si:dkey-65j6.2* tended to have a smaller increase in heart rate from 2 to 5dpf. An opposite trend was observed in embryos with mutations in *quo* (**Fig 3, Supp Table 7)**. Similarly, embryos with nonsense mutations in both *neo1a* alleles showed a larger increase in heart rate from 2 to 5dpf than embryos free from CRISPR/Cas9-induced mutations, without showing a difference in heart rate at 2 or 5dpf (**Fig 3, Supp Table 8**). This suggests that 5dpf may have been too early to detect true genetic effects of mutations in *KIAA1755* and/or *NEO1* orthologues on heart rate.

### Effect of mutations in candidate genes on body size

Besides having a lower heart rate at 2dpf, embryos with CRISPR/Cas9-induced mutations in *rgs6* were shorter at 2 and 5 dpf, and had a smaller dorsal body surface area normalized for body length at 5dpf (**Supp Figs 4, 7; Supp Table 9**). Embryos with mutations in *si:dkey-65j6.2* tended to be leaner at both 2 and 5dpf, while embryos with mutations in *quo* were larger at both time points (**Supp Figs 4, 7: Supp Tables 9-10**). Embryos with mutations in *neo1a* tended to be larger at 2dpf but not at 5dpf (**Supp Figs 4, 7; Supp Tables 9-10**).

### Transcriptomic analyses

Mutagenesis by targeting a proximal site in the protein-coding sequence using CRISPR/Cas9 has been shown to trigger a compensatory upregulation of the expression of transcripts with sequence similarity^25^. To explore if such compensation may have influenced our results, we next targeted *hcn4* and/or *hcn4l* using CRISPR/Cas9 at the single cell stage, pooled five embryos per sample at 5dpf, and performed a qRT-PCR analysis to examine compensatory effects on transcripts with at least 75% sequence similarity to the main zebrafish hcn4 transcript (**Supp Tables 11-12**). Compared with control-injected embryos, targeting *hcn4* or *hcn4* and *hcn4l* simultaneously – using the same gRNAs and protocols used to generate CRISPR/Cas9 founders – resulted in a lower expression of *hcn4* at the CRISPR/Cas9-targeted site at 5dpf, confirming on-target activity for the *hcn4* gRNA (**Fig 6**). We observed no effect of targeting *hcn4* on expression of the last exon of *hcn4*, which likely reflects the low proportion of embryos with a nonsense mutation in both *hcn4* alleles in the phenotypic screen (i.e. 2.4%, **Supp Table 4**). However, we did observe what may be a compensatory increase in the expression of zebrafish orthologues of *HCN1* and *HCN2* in samples of *hcn4*-targeted embryos (**Fig 6**). While the effect of targeting *hcn4* on the expression of *CABZ01086574.1* (likely an orthologue of *HCN2*) and the trend for an effect on *hcn1* expression were driven by one sample of *hcn4*-targeted embryos, the compensatory effect on *hcn2b* expression was more robust (**Supp Table 13**). Possible compensatory effects were observed when targeting *hcn4*, but not when targeting *hcn4* and *hcn4l* simultaneously.

### Nanopore off-target sequencing

We next used *in vitro* Nanopore off-target sequencing (Nano-OTS)^26^ in DNA from an adult Tg(*acta2*:*GFP*) positive fish used to generate CRISPR/Cas9 founders to explore if off-target mutagenic activity may have influenced the results for *hcn4*; and if it may have played a role in the relatively large number of missed sequencing calls for *neo1b* (**Supp Table 4**). No off-target mutations were identified for the *hcn4* and *neo1b* gRNAs. In spite of five mismatches, we did observe one off-target mutation for the *hcn4l* gRNA, inside the coding region of the glutamine transporter *slc38a3a* (**Supp Fig 8**). This off-target is unlikely to have influenced our results, since we did not observe effects of CRISPR/Cas9-induced mutations in *hcn4l*, and since the off-target mutagenic activity was even lower than the on-target activity for this gRNA (**Supp Fig 8**). Furthermore, *in vitro* off-target activity does not necessarily imply *in vivo* mutagenic activity.

One off-target - in spite of four mismatches - was observed for the *neo1a* gRNA. Since this off-target was also characterized by low mutagenic activity and was located >30kb away from the nearest gene (**Supp Fig 8**), large indel mutations at the *neo1b* target site due to *neo1a* or *neo1b* gRNA off-target activity are highly unlikely. Hence, it seems safe to conclude that either the 78 missing sequencing calls for *neo1b* were missing at random - e.g. due to inherently present variants interfering with primer binding – or that they were caused by large indel mutations in *neo1b* resulting from on-target *neo1b* gRNA activity that did not subsequently influence outcomes of interest.

### Potential druggability

Three of the six human candidate genes taken forward for experimental follow-up showed evidence for a role in cardiac rhythm and/or rate in zebrafish embryos (i.e. *HCN4*, *RGS6*, *KIAA1755*). Only a trend for an effect on sinoatrial pauses was observed for the *SYT10* orthologue, albeit with an opposite direction of effect at 2 and 5dpf. We next examined whether these four genes are already targeted by existing, FDA-approved medication using the drug-gene interaction (DGI) database (www.dgidb.org). Only *HCN4* is currently targeted^27^, i.e. using ivabradine, an open channel blocker of I_f_ channels^28^ (see below). We next identified predicted interaction partners of the proteins encoded by the four genes, using GeneMania^29^ and STRING^30^. Some partners are targeted by FDA-approved medication (**Supp Table 14**), amongst which are several anti-hypertensive agents (e.g. Hydrocholorothiazide and Diazoxide), as well as a neuromuscular blocking agent (i.e. Botulinum toxin type A) and statins.

### Ivabradine

Given the role of mutations in *hcn4* on HRV and heart rate, we next examined the effect of 24h of exposure to 0, 10 or 25 µM ivabradine in DMSO on these outcomes at 5dpf, in embryos free from CRISPR/Cas9-induced mutations. One embryo with a sinoatrial pause was excluded from the analysis, as were 13 embryos with suboptimal image or image quantification quality. In the remaining 118 embryos, ivabradine treatment resulted in a dose dependent higher HRV and a lower heart rate, i.e. directionally opposite to the effects of mutations in *hcn4* (**Fig 5, Supp Fig 9, Supp Table 15**). In fact, the effect on heart rate was of similar size – but opposite direction - for 10µM ivabradine when compared with nonsense mutations in both *hcn4* alleles, in data from the CRISPR/Cas9 and sa11188 experiments combined (**Fig 5**). Interestingly, the effect of ivabradine on heart rate was 1.5 to 1.6-fold higher than its effect on HRV (**Fig 5, Supp Table 15**).

## Discussion

Large-scale, *in vivo* follow-up studies of candidate genes in GWAS-identified loci remain sparse. Here we present an objective, image-based pipeline to systematically characterize candidate genes for cardiac rhythm, rate, and conduction-related disorders. Using a zebrafish model system, we confirmed a role for genes previously implicated in heart rate and rhythm (*rgs6* and *hcn4*); show that effects of ivabradine on heart rate and rhythm are opposite when compared with the early-stage effects of mutations in one of the drug’s molecular targets (*hcn4*); identified a previously unanticipated gene influencing heart rate variability (*si:dkey-65j6.2*, i.e. *KIAA1755*); and observed effects of previously unanticipated genes on early growth and development (*rgs6*, *quo*, *si:dkey-65j6.2*, *neo1a*). In addition, we confirmed that mutations in *hcn4* increase the odds of sinoatrial pauses and arrests^15^. The latter adds weight to the notion that GWAS-identified common variants for complex traits can flag genes for which rare, detrimental mutations cause severe, early-onset disorders^7,31,32^. We show here that an image- and CRISPR/Cas9-based zebrafish model system can be used for systematic characterization of candidate genes in GWAS-identified loci for complex traits. In the near future, integration of evidence across a range of species and approaches should close the existing gap between association and functional understanding. This will no doubt yield new targets that can be translated into efficient new medication for prevention and treatment of complex diseases.

The locus harboring *HCN4* has been identified in GWAS for HRV^3^, heart rate^4^ and atrial fibrillation^33^. *HCN4* belongs to the family of I_f_ or “funny” channels, aptly named for being activated upon hyperpolarization and a non-selective permeability for Na^+^ and K^+^. It is expressed in the sinoatrial node^34,35^ and plays an important role in cardiac pace making^36^. The heart rate lowering agent ivabradine^37^ is an open channel blocker of I_f_ channels, and has been shown to reduce heart rate and increase HRV in humans^38,39^. It also reduces cardiovascular and all-cause mortality in heart failure patients^40^. In line with findings in humans, one day of treatment with ivabradine resulted in a dose dependent lower heart rate and higher HRV in 5dpf zebrafish embryos. Since the drug had a 1.5 to 1.6 fold larger effect on heart rate than on HRV, its protective effect on cardiovascular mortality in heart failure patients may indeed be driven by its heart rate lowering effect^28^. The larger effect size for heart rate than for HRV also suggests that ivabradine may influence HRV at least in part through its non-vagal effects on heart rate, supporting an intrinsic effect of heart rate on HRV, as has been postulated earlier^41^. Exercise training-induced bradycardia has previously been shown to result from HCN4 downregulation^42^. Our findings suggest that downregulation of HCN4 may be at least partly responsible for the higher HRV in exercisers, in contrast or addition to a higher vagal tone in exercisers^43^. With only one affected embryo in the ivabradine experiment, we could not examine the role of the drug in prevention of sinoatrial pauses.

Across two separate experiments, we observed that embryos with nonsense mutations in both *hcn4* alleles have 9-fold higher odds of sinoatrial pauses at 5dpf, as well as a *higher* heart rate and *lower* HRV in embryos free from sinoatrial pauses.

Hence, nonselective open I_f_ channel blocking using ivabradine and nonsense mutations in *hcn4* have directionally opposite effects on heart rate and HRV in 5dpf zebrafish embryos. Others previously showed that *HCN4^+/-^* humans are typically characterized by bradycardia^44^; *Hcn4^-/-^* mice die prenatally, without arrhythmias and with lower heart rates when compared with *Hcn4^+/-^ and Hcn4^+/+^* mice^36^; and 2dpf zebrafish embryos with experimentally downregulated *hcn4* expression vs. un-injected controls have a higher odds of sinoatrial arrests and lower heart rate^15^. In this light, a higher heart rate in 5dpf zebrafish embryos with nonsense mutations in both *hcn4* alleles could be considered unexpected. However, results from our qRT-PCR experiment suggest that this higher heart rate is likely driven by a compensatory upregulation of the expression of *HCN1* and *HCN2* orthologues. The potential compensatory increase in *hcn1* and *CABZ01086574.1* expression is driven by only one sample, but since technical triplicates show coherent results, this may reflect the low mutagenic efficiency of the *hcn4* gRNA. The likely compensatory increase in *hcn2b* expression upon CRISPR/cas9 targeting of *hcn4* was more robust. Results from a Nanopore-OTS screen showed that off-target activity of the *hcn4* gRNA is highly unlikely to explain our results. Thus, it seems likely that a higher heart rate in embryos with nonsense mutations in *hcn4* - likely facilitated by a higher expression of *HCN1* and *HCN2* orthologues – serves as an attempt to prevent sinoatrial arrests and sudden cardiac death. This physiological attempt to compensate a genetic defect will no doubt be followed by the bradycardia observed in other species once the heart can no longer cope with the persistently elevated workload. Besides from an absence of the compensatory response when downregulating gene expression using morpholino oligonucleotides^45^, the lower heart rate previously reported in 2dpf zebrafish embryos with downregulated *hcn4*^15^ could also have resulted from RNA toxicity, off-target effects, or a developmental delay caused by the microinjection itself^15,46^.

Regulator of G protein signaling 6 (RGS6) plays a role in the parasympathetic regulation of heart rate^47^ and is a negative regulator of muscarinic signaling, thus decreasing HRV to prevent bradycardia. Common variants near *RGS6* have been identified in GWAS for resting heart rate^6,48,49^, heart rate recovery after exercise^49,50^ and HRV^3^. An eQTL analysis using GTEx data showed that the minor T-allele in rs4899412 - associated with lower HRV - is also associated with a higher expression of *RGS6* in whole blood and tibial nerve. This is directionally consistent with humans with loss-of-function variants in *RGS6* showing higher HRV^51^. Also in line with this, *Rgs6^-/-^* mice were previously characterized by lower heart rate, higher HRV and higher susceptibility to bradycardia and atrioventricular block^52^. In our study, zebrafish embryos that were heterozygous for CRISPR/Cas9-induced mutations in *rgs6* on average had a nearly 0.5 SD units lower heart rate at 2dpf. Hence, our results for heart rate at 2dpf are in line with studies in mice. The effect on heart rate was no longer apparent at 5dpf, in spite of a slightly better statistical power. We did not detect an effect of mutations in *rgs6* on HRV or risk of sinoatrial pauses or arrests at 2 or 5dpf.

Non-synonymous SNPs in *KIAA1755* have been identified in GWAS for HRV^3^ and heart rate^5^. In our analysis, mutations in *si:dkey-65j6.2* tend to result in a higher HRV at 2 and 5dpf, but do not affect heart rate. In humans, the minor C-allele of the HRV-associated rs6123471 in the non-coding 3’ UTR of *KIAA1755* tends to be associated with a lower expression of *KIAA1755* in the left ventricle and atrial appendage, amongst other tissues (GTEx^53^); as well as with a higher HRV^3^ and a lower heart rate^4^. Hence, a higher HRV at 2 and 5dpf for each additional mutated allele in *si:dkey-65j6.2* is directionally consistent with results in humans. Our results suggest that the GWAS-identified association in the *KIAA1755* locus - and possibly other loci - with heart rate may have been driven by HRV. This would explain why the eleven HRV-associated loci that showed evidence of an association with heart rate all did so in the expected (i.e. opposite) direction from a phenotypic point of view^3^.

*KIAA1755* is a previously uncharacterized gene that shows a broad expression pattern, including different brain regions and the left atrial appendage (GTEx^53^). Future mechanistic studies are required to distill how *KIAA1755* influences heart rhythm.

In addition to a comparison of specimens with nonsense mutations in both alleles vs. zero mutated alleles, we examined the role of mutations in each gene using an additive model. The much larger sample size of this alternative approach helped put results from the more traditional approach into perspective. In our pipeline, we decided to raise founders to adulthood, and to phenotypically characterize and sequence F_1_ embryos, with stable genotypes. Screening the F_0_ or F_2_ generation would have each had their own advantages and disadvantages. Wu and colleagues described a powerful approach to efficiently disrupt and analyze F_0_ embryos^54^. However, this approach requires drawing conclusions from results in injected, mosaic founders, and is not compatible with multiplexing of multiple gRNAs and target sites. Screening the F_2_ generation on the other hand requires hand picking of F_1_ fish with suitable mutations, which limits the throughput.

The large differences in mutant allele frequency across the targeted genes can be attributed to several factors. First, mosaic founders were in-crossed six times, using random mating. Hence, different founder fish may have produced the screened F_1_ offspring in each crossing. Secondly, while all gRNAs were pre-tested for mutagenic activity, mutated alleles detected in the test injections that didn’t affect the germline were not included in our screen. Thirdly, mutations that are embryonic lethal prior to 2dpf did not make it into the screen.

Of the three genes for which we show effects of mutations on cardiac rhythm and/or rate, only *HCN4* is currently targeted by FDA-approved drugs^55^. Although the other two genes are not highlighted as being druggable by small molecules, targeting via antisense oligonucleotides, antibodies, or other approaches can be explored.

Furthermore, expanding our search revealed several druggable interaction partners of putative causal genes that are already targeted by FDA-approved anti-hypertensive and neuromuscular blocking agents, amongst others. *SNAP25*, a proposed interaction partner of *SYT10*, is the target of botulinum toxin A. Injection of botulinum toxin A into the neural ganglia of CAD/atrial fibrillation patients was recently shown to reduce the occurrence of post-operational atrial fibrillation^56^. For existing drugs that target interaction partners of putative causal genes, it is worthwhile examining the effect on cardiac rhythm, rate and development, since repurposing FDA-approved drugs would imply the quickest and safest route to the clinic. Quantifying possibly unknown beneficial or adverse side effects of these drugs related to cardiac rhythm would also be informative.

Four potential limitations of our approach should be discussed. Firstly, acquiring 30s recordings implies that false negatives for sinoatrial pauses or arrests are inevitable. We decided to exclude the embryos with a sinoatrial pause or arrest during positioning under the microscope from the analysis, because the case status of these embryos cannot be confirmed objectively, but they are not appropriate controls either. This limitation will at most have resulted in conservative effect estimates.

Secondly, CRISPR/Cas9 gRNAs with predicted off-target effects free from mismatches were avoided. However, two of the selected targets - i.e. for *hcn4* and *si:dkey-65j6.2* - had predicted off-target activity with three mismatches in *galnt10* and *dclk1*, respectively, at the time we designed the gRNAs (**Supp Table 2**). Human orthologues of these two genes have previously been associated with heart rate variability-related traits^57^ (*dclk1*), as well as with carotid intima-media thickness^58^ and body mass index^59–61^ (*galnt10*). However, none of the 381 embryos carried CRISPR/Cas9-induced mutations at the three predicted off-target sites (**Supp Table 2**). Furthermore, *in vitro* genome-wide Nano-OTS only revealed two off-target sites across the four gRNAs of *HCN4* and *NEO1* orthologues, both with low mutagenic activity. Taken together, this implies that gRNA off-target mutagenic activity is extremely unlikely to have influenced our results. Thirdly, we in-crossed mosaic founders (F_0_) and phenotypically screened and sequenced the F_1_ generation. For some genes, this yielded a very small number of embryos with 0 or 2 mutated alleles (i.e. for *rgs6*, *hcn4*, and *hcn4l*), resulting in a low statistical power to detect a true role for mutations in these genes. In spite of this limitation, we still detected significant effects of mutations in *rgs6* and *hcn4*, and confirmed our results for mutations in *hcn4* on heart rate and risk of sinoatrial pauses in an independent experiment with more statistical power. Unfortunately, the frame rate of image acquisition in the pilot experiment was too low to also replicate effects of mutations in *hcn4* on HRV. Finally, we recorded the atrium only, to enable a higher frame rate, a higher resolution in time for HRV quantification, and a higher statistical power to detect small genetic effects on HRV. As a result, any ventricular abnormalities that may have occurred were not registered, and uncontrolled atrial contractions may thus reflect atrial fibrillation, premature atrial contractions, high atrial rate, or atrial tachycardia.

Strengths of our study include its repeated measures design, which enabled us to capture genetic effects at different stages of early development in zebrafish, as well as genetic effects on phenotypic changes over time. Furthermore, the throughput of the setup allowed us to examine the effect of mutations in multiple genes simultaneously, in a larger than usual sample for *in vivo* genetic screens. Our results demonstrate that a large sample size is paramount to robustly detect genetic effects on complex traits in zebrafish embryos when screening the F_1_ generation, even for nonsense mutations.

Identifying CRISPR/Cas9-induced mutations allele-specifically using a custom-written algorithm helped us distinguish between heterozygous and compound heterozygous embryos, which in turn helped pinpoint the effect of mutations in these genes. Also, our study is based on a high-throughput imaging approach with objective, automated quantification, as compared with manual counting and annotation of heart rate in most^4^ but not all^12,62^ earlier studies. Finally, for all genes that showed an effect on HRV, observed effects were directionally consistent with eQTL associations in humans. This further emphasizes the strength of our model system and the robustness of our findings.

In conclusion, our large-scale imaging approach shows that zebrafish embryos can be used for rapid and comprehensive follow-up of GWAS-prioritized candidate genes for HRV and heart rate. This will likely increase our understanding of the underlying biology of cardiac rhythm and rate, and may yield novel drug targets to prevent cardiac death.

## Methods

### Candidate gene selection

Candidate genes in GWAS-identified loci for HRV were identified as described in detail in Nolte et al.^3^. Of the 18 identified candidate genes, six were selected for experimental follow-up. This selection was based on overlap with findings from GWAS for heart rate (*KIAA1755*, *SYT10*, *HCN4*, *GNG11*)^4^, as well as with results from eQTL analyses in sinoatrial node and brain (*RGS6*). Additional candidate genes from the same or nearby loci were also selected for experimental follow-up, i.e. *NEO1*, which resides next to *HCN4*. Zebrafish orthologues of the human genes were identified using Ensembl, as well as using a comprehensive synteny search using Genomicus^63^ (**Supp Table 1**). Of the selected genes, *GNG11*, *SYT10* and *RGS6* have one orthologue in zebrafish, and *HCN4*, *NEO1* and *KIAA1755* each have two orthologues, resulting in a total of nine zebrafish orthologues for six human candidate genes (**Table 1**).

### CRISPR/Cas9-based mutagenesis

All nine zebrafish genes were targeted together using a multiplexed CRISPR/Cas9 approach^20^. Briefly, guide-RNAs (gRNAs) were selected using ChopChop^64^ and CRISPRscan^65^ (**Supp Table 2**), based on their predicted efficiency, a moderate to high GC-content, proximal location in the protein-coding sequence, and absence of predicted off-target effects without mismatches. Oligonucleotides were designed as described^21^, consisting of a T7 or SP6 promoter sequence (for gRNAs starting with ‘GG’ or ‘GA’, respectively), a gene-specific gRNA-target sequence, and an overlap sequence to a generic gRNA. The gene-specific oligonucleotides were annealed to a generic 80bp long oligonucleotide at 98°C for 2 mins, 50°C for 10 mins, and 72°C for 10 mins. The products were checked for correct length on a 2% agarose gel. The oligonucleotides were subsequently transcribed *in vitro* using the manufacturer’s instructions (TranscriptAid T7 high yield transcription kit / MEGAscript SP6 transcription kit, both ThermoFisher Scientific, Waltham, USA). The gRNAs were purified, after which the integrity of the purified gRNAs was examined on a 2% agarose gel. The zebrafish codon-optimized plasmid pT3TS-nls-zCas9-nls was used as a template to produce Cas9 mRNA^66^. The plasmid was linearized with Xba1, and then purified using the Qiaprep Spin Miniprep kit (Qiagen, Hilden, Germany). The DNA was transcribed using the mMESSAGE mMACHINE T3 Transcription Kit (ThermoFisher Scientific, Waltham, USA), followed by LiCl precipitation. The quality of the RNA was confirmed on a 1% agarose gel.

### Husbandry & microinjections

A zebrafish line with GFP-labelled α-smooth muscle cells Tg(*acta2*:*GFP*)^21^ was used to visualize the beating heart. To this end, eggs from an in-cross of Tg(*acta2*:*GFP*) fish were co-injected with a mix of Cas9 mRNA (final concentration 150 ng/µl) and all nine gRNAs (final concentration 25 ng/µl each) in a total volume of 2nL, at the single-cell stage. CRISPR/Cas9 injected embryos were optically screened for the presence of Tg(*acta2*:*GFP*) at 2 days post fertilization (dpf), using an automated fluorescence microscope (EVOS FL Cell imaging system, ThermoFisher Scientific, Waltham, USA). Tg(*acta2*:*GFP*) carriers were retained and raised to adulthood in systems with circulating, filtered and temperature controlled water (Aquaneering, Inc, San Diego, CA). All procedures and husbandry were conducted in accordance with Swedish and European regulations, and have been approved by the Uppsala University Ethical Committee for Animal Research (C142/13 and C14/16).

### Experimental procedure imaging

The mosaic founders (F_0_ generation) were only used to yield offspring (F_1_ embryos) for experiments. To reach the experimental sample size, founders were in-crossed six times using random mating. Eggs were collected after founders were allowed to reproduce for 45 mins, to minimize variation in developmental stage. Fertilized eggs were placed in an incubator at 28.5°C. At 1dpf, embryos were dechorionated using pronase (Roche Diagnostics, Mannheim, Germany).

At 2dpf, embryos were removed from the incubator and allowed to adapt to controlled room temperature (21.5 °C) for 20 mins. Individual embryos were exposed to 100 µg/ml Tricaine (MS-222, Sigma-Aldrich, Darmstadt, Germany) for 1 min before being aspirated, positioned in the field of view of a fluorescence microscope, and oriented dorsally using a Vertebrate Automated Screening Technology (VAST) BioImager (Union Biometrica Inc., Geel, Belgium). We subsequently acquired twelve whole-body images, one image every 30 degrees of rotation, using the camera of the VAST BioImager, to quantify body length, dorsal and lateral surface area and volume, as well as the presence or absence of cardiac edema. The VAST BioImager then positioned and oriented the embryo to visualize the beating atrium and triggered the upright Leica DM6000B fluorescence microscope to start imaging using an HCX APO L 40X/0.80 W objective and L5 ET, k filter system (Micromedic AB, Stockholm, Sweden). Images of the beating atrium were acquired for 30s at a frame rate of 152 frames/s using a DFC365 FX high-speed CCD camera (Micromedic AB, Stockholm, Sweden). After acquisition, the embryos were dispensed into a 96-well plate, rinsed from tricaine, and placed back into the incubator. The procedure was repeated at 5dpf, to allow capturing of genetic effects that influence HRV and heart rate differently at different stages of development^22^. After imaging at 5dpf, the embryos were once again dispensed into 96-well plates, sacrificed, and stored at −80°C for further processing.

### Quantification of cardiac traits and body size

A custom-written MATLAB script was used to convert the images acquired by the CCD camera into quantitative traits. To acquire the heart rate, each frame of the sequence was correlated with a template frame. The repeating pattern yields a periodic graph from the correlation values and by detecting the peaks in the graph we can assess the heart rate. The template frame should represent one of the extreme states in the cardiac cycle, i.e. end-systole or end-diastole. To detect these frames, we examined the correlation between the first 100 frames. The combination of frames that showed the lowest correlation corresponded to the heart being in opposite states. One of these frames was chosen as the template^67^. This numeric information was subsequently used to quantify: 1) heart rate as the inverse of RR-interval; 2) the standard deviation of the normal-to-normal RR interval (SDNN); and 3) the root mean square of successive heart beat interval differences (RMSSD). Finally, a graph of pixel changes over time was generated across the 30s recording to help annotate the script’s performance. Inter-beat-intervals were used to objectively quantify sinoatrial pauses (i.e. the atrium stops contracting for longer than 3x the median inter-beat-interval of the embryo, **Figure 2, Supp recording 1**) and sinoatrial arrests (i.e. the atrium stops contracting for longer than 2s, **Figure 2, Supp recording 2**). The graphs of pixel changes over time were also used to identify embryos with other abnormalities in cardiac rhythm. Such abnormalities were annotated as: uncontrolled atrial contractions (**Figure 2, Supp Recording 3**); abnormal cardiac morphology (i.e. a tube-like atrium, **Figure 2, Supp Recording 4**); or impaired cardiac contractility (i.e. a vibrating rather than a contracting atrium, **Supp Recording 4**). These phenotypes were annotated independently by two investigators (BvdH and MdH), resulting in an initial concordance rate >90%. Discrepancies in annotation were discussed and re-evaluated to reach consensus.

Bright-field images of the embryos were used to assess body length, dorsal and lateral surface area, and body volume. Images were automatically segmented and quantified using a custom-written CellProfiler^68^ pipeline, followed by manual annotation for segmentation quality. Embryos with suboptimal segmentation quality due to the presence of an air-bubble on the capillary, a bent body, an incomplete rotation during imaging, partial capturing of the embryo, or an over-estimation of size were replaced by images with a 180° difference in rotation, or excluded from the analysis for that outcome if the second image was also sub-optimally segmented. The embryo was excluded from the analysis for body volume if more than four of the 12 images had a bad segmentation. Imaging, image quantification and image quality control were all performed blinded to the sequencing results.

### Quality control of phenotype data

A series of quality control steps was performed to ensure only high-quality data was included in the genetic association analysis (**Supp Fig. 2**). First, graphs indicating that one or more true beats were missed by the script were removed from the analysis *a priori* (**Supp Fig. 2**). Second, embryos showing uncontrolled atrial contractions, sinoatrial pauses or arrests, edema, abnormal morphology and/or reduced contractility, and embryos showing a sinoatrial pause or arrest during positioning under the microscope were excluded from the analyses for HRV and heart rate (**Suppl Fig. 2**). The latter were also excluded from the analysis for sinoatrial pauses and arrests, since we cannot ascertain case status for sinoatrial pauses and arrests in the same rigorous manner for such embryos, but they are not appropriate controls either. Third, genetic effects were only examined for cardiac outcomes with at least ten cases, i.e. for sinoatrial pauses and arrests only.

### Sample preparation for sequencing

After imaging at 5dpf, embryos were sacrificed and DNA was extracted by exposure to lysis buffer (10mM Tris-HCl pH8, 50mM KCl, 1mM EDTA, 0.3% Tween 20, 0.3% Igepal) and proteinase K (Roche Diagnostics, Mannheim, Germany) for 2 h at 55°C, followed by 10 min at 95°C to deactivate the proteinase K. Gene-specific primers (150bp-300bp) amplifying gRNA-targeted and putative off-target regions in *dclk1b* and both *galnt10* orthologues (**Supp Table 2**) were distilled from ChopChop^64^ and Primer3^69^, and Illumina adaptor-sequences were added. Additionally, we included 96 un-injected Tg(*acta2*:*GFP*) embryos for sequencing across all gene-specific regions to not mistake naturally occurring variants for CRISPR/Cas9-induced mutations in our main exposures. The first PCR was conducted by denaturation at 98°C for 30s; amplification for 35 cycles at 98°C for 10s, 62°C for 30s and 72°C for 30s; followed by a final extension at 72°C for 2 mins. Amplified PCR products were cleaned using magnetic beads (Mag-Bind PCR Clean-up Kit, Omega Bio-tek Inc.

Norcross, GA). The purified products were used as a template for the second PCR, in which Illumina Nextera DNA library sequences were attached to allow multiplexed sequencing of all CRISPR/Cas9-targeted sites across 383 embryos in a single lane.

The second PCR amplification was performed by denaturation at 98°C for 30s; amplification for 25 cycles at 98°C for 10s, 66°C for 30s and 72°C for 30s; followed by a final extension at 72°C for 2 mins. Products were then purified using magnetic beads. All liquid handling was performed using a Hamilton Nimbus robot equipped with a 96-head (Hamilton robotics, Bonaduz, Switzerland). Samples were pooled and sequenced in a single lane on a MiSeq (2×250 bp paired-end, Illumina Inc., San Diego, CA) at the National Genomics Infrastructure, Sweden.

### Processing of sequencing data

A custom-written bioinformatics pipeline was developed in collaboration with the National Bioinformatics Infrastructure Sweden, to prepare .fastq files for analysis. First, a custom-written script was used to de-multiplex the .fastq files by gene and well. PEAR^70^ was then used to merge paired-end reads, followed by removal of low-quality reads using FastX^71^. The reads were then mapped to the wildtype zebrafish genome (Zv11) using STAR^72^. Next, we converted files from .sam to .bam format using samtools^73^, after which variants - mostly indels and SNVs - were called allele specifically using a custom-written variant calling algorithm in R (Danio rerio Identification of Variants by Haplotype - DIVaH). A summary of all unique sequences identified at each targeted site is shown in **Supp Fig 5**. All unique variants (**Supp Table 3**) located within ±30bps of the CRISPR/Cas9-targeted sites that were identified across the two alleles were subsequently pooled, and used for functional annotation using Ensembl’s VEP^74^. Naturally occurring variants based on sequencing of un-injected Tg(*acta2:GFP*) embryos or Ensembl were excluded. In absence of a continuous score provided by variant effect prediction algorithms, we attributed weights of 0.33, 0.66 and 1 for variants with a predicted low, moderate and high impact on protein function. Pilot experiments suggested that this was a more powerful approach than not weighting. Transcript-specific dosage scores were then calculated by retaining the variant with the highest predicted impact on protein function for each allele, target site, and embryo, followed by summing the scores across the two alleles at each target site and embryo. Since all transcripts within a target site were affected virtually identically, we only used the main transcript of each target site for the genetic association analysis.

Most embryos had successfully called sequences in all nine CRISPR/Cas9-targeted sites. Embryos with missing calls at more than two of the nine targeted sites were excluded from the genetic association analysis. For the remaining 381 embryos, missing calls were imputed to the mean dosage of the transcript. For *neo1b*, calling failed in 78 embryos (**Supp Table 4**). The imputed mean dosage for the main transcript of *neo1b* was still included in the genetic association analysis, since: 1) the mutant allele frequency in successfully called embryos was very high (i.e. 0.934) and an imputed call thus likely closely resembles the truth; and 2) the distribution of embryos with a missed call was similar for all outcomes when compared with the remaining embryos. Hence, using an imputed dosage was concluded to influence the results less than to either exclude the gene from the analysis while it had been targeted; or to discard the 78 embryos since they had been successfully phenotyped and sequenced for the other targeted genes. However, the results for *neo1b* should still be interpreted in light of its call rate. The mutant allele frequency was low for *hcn4*, *hcn4l* and *rgs6*. No embryos carried CRISPR/Cas9-induced mutations within ±30bp of any of the three putative off-targets regions (i.e. *dclk1b* and both *galnt10* orthologues, **Supp Table 2**).

### qPCR to evaluate targeting hcn4 and hcn4l

We performed a qRT-PCR experiment to examine if targeting hcn4 and/or hcn4l resulted in a compensatory upregulation of the expression of transcripts with >75% sequence similarity to the main transcript of *hcn4*. To this end, fertilized eggs of Tg(acta2*:GFP*) positive fish were either: 1) left un-injected; or injected at the single cell stage with: 2) Cas9 mRNA only; 3) *hcn4* and *hcn4l* gRNA only; 4) *hcn4* gRNA and Cas9 mRNA; 5) *hcn4l* gRNA and Cas9 mRNA; or 6) *hcn4* and *hcn4l* gRNAs and Cas9 mRNA. The same protocols, quantities and gRNAs were used as described above for generating founders (see: *Husbandry & microinjections*). Embryos were raised to 5dpf in an incubator at 28.5°C. On the morning of day 5, within each of the six conditions, multiple pools of five embryos per sample were flash frozen in liquid nitrogen. The experiment was performed twice, with all conditions generated on both occasions, to generate a total of 69 samples for conditions 1 to 6 described above.

Samples were homogenized in Trizol using a 20-gauge needle and a 1ml syringe. RNA was extracted using the TRIzol™ Plus RNA Purification Kit and Phasemaker™ Tubes Complete System (Cat.No: A33254, Invitrogen, Waltham, MA). The quality and concentration of the extracted RNA was measured with the Agilent RNA 6000 nano kit (Cat-No: 5067-1511, Agilent, Santa Clara, CA) on an Agilent 2100 Bioanalyzer. *In vitro* transcription of 100ng of RNA per sample was performed using the SuperScript IN VILO Master Mix (Cat.No: 11755500, Invitrogen), and quantitative PCR was performed in triplicate using the PowerUp SYBR Green Master Mix on either an AriaMx (Agilent) or a StepOnePlus (Applied Biosystems, Waltham, MA) Real-Time PCR System. Pooled cDNA samples were used to optimize the primer concentrations and generate standard curves for the calculation of primer efficiencies. Primers with an efficiency close to 100% and a single peak in melt curve analysis were selected (**Supp Table 11**). We used 1ng of cDNA as input for the rest of the reactions with the following cycle conditions: 50°C for 2 min (1 cycle), 95°C for 2 min (1 cycle), 95°C for 3s and 60°C for 30s (40 cycles). Differences between the three experimental conditions (conditions 4 to 6) and the two injected controls combined (conditions 2 and 3) on gene expression at the *hcn4* and *hcn4l* CRISPR/Cas9-targeted site and at the last exon, as well as for *CABZ01086574.1* (*HCN2*), *hcn2b*, *hcn3* and *hcn1* (**Supp Table 11**) were examined using Pfaffl’s method^75^, taking into account primer efficiencies (**Supp Table 12**); expression levels in non-injected controls as calibrator; and expression of *mob4* as the reference gene^76^.

### Nanopore off-target sequencing (OTS)

We next used Nanopore off-target sequencing (Nano-OTS) to explore whether CRISPR/Cas9 off-target activity may have influenced the results for *hcn4*, and whether off-targets may have caused the relatively large number of missed calls for *neo1b*. To this end, DNA was extracted using the MagAttract HMW kit (Qiagen, Hilden, Germany) according to the manufacturer’s instructions for extraction from tissue. Genomic DNA was fragmented to 20 kb using Megaruptor 2 (Diagenode, Liege, Belgium) and size selected with a 10 kb cut-off using the Blue Pippin system (Sage Science, Beverly, MA). Libraries for Nano-OTS were prepared as described recently by Höijer et al^26^. In short, ribonucleoproteins (RNPs) were prepared as a pool using the *hcn4*, *hcn4l*, *neo1a* and *neo1b* gRNAs. Next, 3µg of fragmented and size-selected DNA was dephosphorylated and digested by Cas9 using the RNPs, and DNA ends were dA-tailed. Finally, sequencing adapters from the SQK-LSK109 kit (Oxford Nanopore Technologies, Oxford, UK) were ligated to the Cas9-cleaved ends.

Sequencing was performed on one R9.4.1 flow cell using the MinION system (Oxford Nanopore Technologies). Guppy v3.4 was used for base calling. Analysis and calling of OTS sites were performed as described by Höijer et al.^26^, using the GRCz11 reference genome.

### Examining druggability of putative causal genes

All human orthologues of the zebrafish genes for which we observed (a trend for) an effect were explored in the drug-gene interaction database (DGIdb) v3.0.2^27^. Possible interaction partners of the human proteins and genes (**Supp Table 14**) were distilled from STRING v10.5^30^ and GeneMania^29^, respectively. The GeneMania search was limited to physical interactions and pathway data.

### Characterizing the effect of treatment with ivabradine

To examine the effect of exposure to ivabradine, the procedures described in “*Experimental procedure imaging*” were repeated in embryos free from CRISPR/Cas9-induced mutations. On the morning of day 4, embryos were placed in 0, 10 or 25µM ivabradine in DMSO, for 24h (Sigma-Aldrich, St Louis, MO). On the morning of day 5, conditions were blinded and embryos were imaged as described in “*Experimental procedure imaging*”. Embryos from the three blinded conditions were imaged in an alternating manner, to prevent differences in developmental stage from influencing the results. The experiment was performed four times to generate a total of 44 embryos per condition. One embryo with a sinoatrial pause and 13 embryos with suboptimal image or image quantification quality were excluded from the analysis, leaving 118 embryos for the analysis of effects on HRV and heart rate.

### Statistical analysis

The standard deviation of NN-intervals (SDNN) and the root mean square of successive differences (RMSSD) were strongly correlated at 2dpf (r^2^=0.78) and at 5dpf (r^2^=0.83), so a composite endpoint ‘HRV’ was calculated as the average of SDNN and RMSSD. In embryos free from sinoatrial pauses and arrests; abnormal cardiac morphology; impaired cardiac contractility; and edema, we inverse-normally transformed HRV and heart rate at 2dpf (n=279) and 5dpf (n=293) to a mean of 0 and a standard deviation of 1, so effect sizes and their 95% confidence intervals can be interpreted as z-scores. This approach also ensured a normal distribution of all quantitative outcomes, and allowed a comparison of effect sizes across traits. For body size analyses, we first normalized dorsal surface area, lateral surface area and volume for body length, and subsequently inverse normalized these outcomes as well as body length for the statistical analysis.

Effects of CRISPR/Cas9-induced mutations in candidate genes on sinoatrial pauses and arrests at 2dpf, and on sinoatrial pauses at 5dpf were examined using a logistic regression analysis. In embryos free from sinoatrial pauses or arrests during the image acquisition, effects of mutations in candidate genes and of ivabradine on HRV and heart rate were examined using hierarchical linear models at 2 and 5dpf separately (Stata’s xtmixed), adjusted for the time of imaging (fixed factor), and with embryos nested in batches (random factor with fixed slope). If a sinoatrial pause or arrest was observed before but not during imaging, embryos were not included in the statistical analysis. In a sensitivity analysis, we mutually adjusted associations for HRV and heart rate for the other outcome (i.e. for heart rate and HRV) by adding the outcome as an independent variable to the model. Effects of CRISPR/Cas9-induced mutations in candidate genes on body size were also examined using hierarchical linear models, as described above.

For dichotomous outcomes and continuous outcomes alike, genetic effects were examined using an additive model, with dosage scores for all nine CRISPR/Cas9 targeted sites as independent exposures in the same model, i.e. mutually adjusted for effects of mutations in the other targeted sites. Additionally, for the six genes where at least five embryos carried CRISPR/Cas9-induced nonsense mutations in both alleles, these embryos were compared with embryos free from CRISPR/Cas9-induced mutations at that site, adjusting for dosage scores in the eight remaining genes. For each outcome, associations were examined for the main transcript of the orthologue.

In the qRT-PCR experiment, the effect of targeting *hcn4*, *hcn4l*, or *hcn4* & *hcn4l* on gene expression was examined using multiple linear regression analysis, adjusting for batch. P-values <0.05 were considered to reflect statistically significant effects. All statistical analyses were performed using Stata MP version 14.2 (StataCorp, College Station, TX).

## Acknowledgments

The computations were performed on resources provided by SNIC through Uppsala Multidisciplinary Center for Advanced Computational Science (UPPMAX) under Project SNIC b2015283. The authors would like to acknowledge support from Science for Life Laboratory, the National Genomics Infrastructure (NGI) and UPPMAX for aiding in massive parallel sequencing and computational infrastructure. Support from the National Bioinformatics Infrastructure Sweden (NBIS) is also gratefully acknowledged. Constructive discussions with Drs Shawn Burgess and Gaurav Varshney, as well as with the Genome Engineering Zebrafish (GEZ) facility are also acknowledged. Support from João Campos Costa when setting up the lab is also acknowledged. MdH is a Beijer Researcher and a fellow of the Swedish Heart-Lung Foundation (20170872). He is supported by project grants from the Swedish Heart-Lung Foundation (20140543, 20170678, 20180706), the Swedish Research Council (2015-03657, 2019-01417), and NIH/NIDDK (R01DK106236, R01DK107786, U01DK105554).

## Author contributions

BvdH and MdH conceived the study; BvdH, SV and SJ developed the experimental protocols; TK and AE implemented the CRISPR/Cas9 pipeline in the lab; BvdH performed the CRISPR/Cas9 experiment; SV performed the sa11188 experiment; AE performed the ivabradine experiment; AAl and HLB developed the image quantification pipeline; OD generated the sequencing quality control pipeline; EM conceived the variant calling algorithm; AE performed the qRT-PCR experiment; IH and AAm developed and performed the Nanopore off-target sequencing; BvdH, HLB and MdH performed the statistical analysis; BvdH, EM, IH, HLB and MdH generated the figures; BvdH and MdH wrote the manuscript; All authors provided critical feedback to the manuscript.

## Competing interests

The authors declare no competing interests.

## Data availability

Raw data and scripts will be made publicly available after publication if desirable.

## SUPPLEMENTARY FIGURES

**Supplementary Figure 1:**
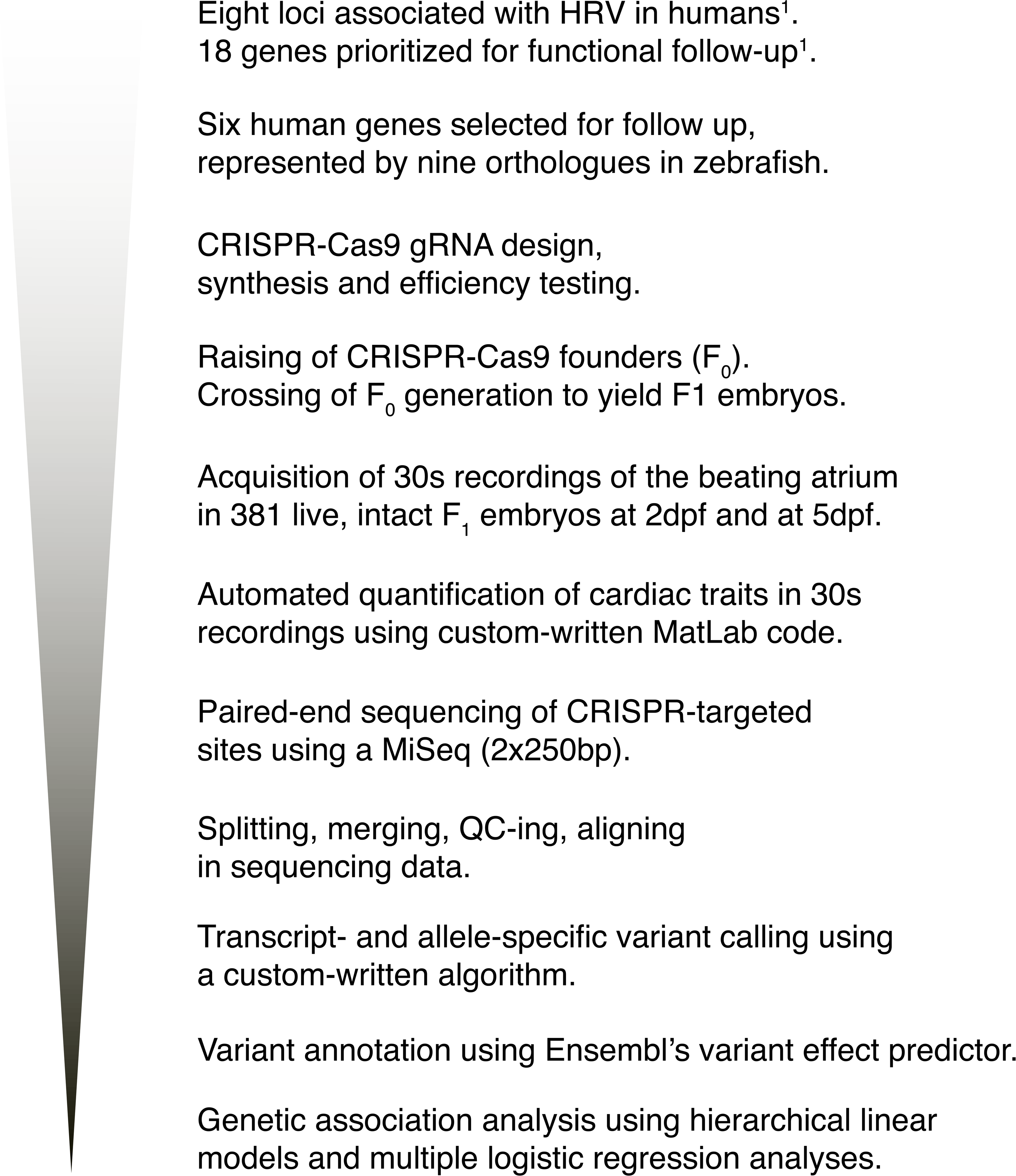
Experimental pipeline describing the workflow from GWAS in humans to stastist^P^ic^ag^a^e^l^48^a°n^f^a^91^lysis in zebrafish embryos.

**Supplementary Figure 2:**
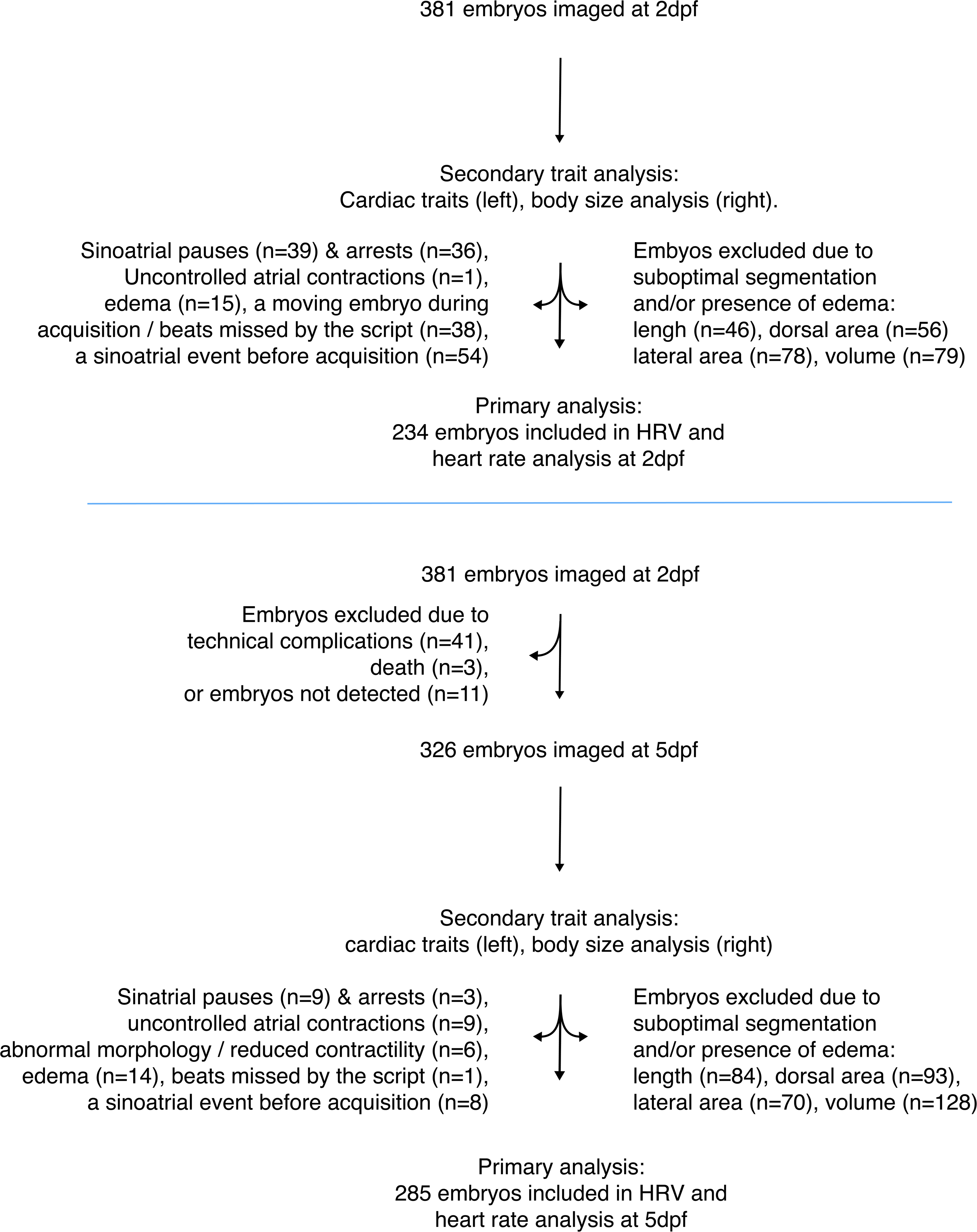
Flow chart showing the of embryos included in the analysis and reasons for exclusion. Embryos can be excluded for more than one reason.

**Supplementary Figure 3:**
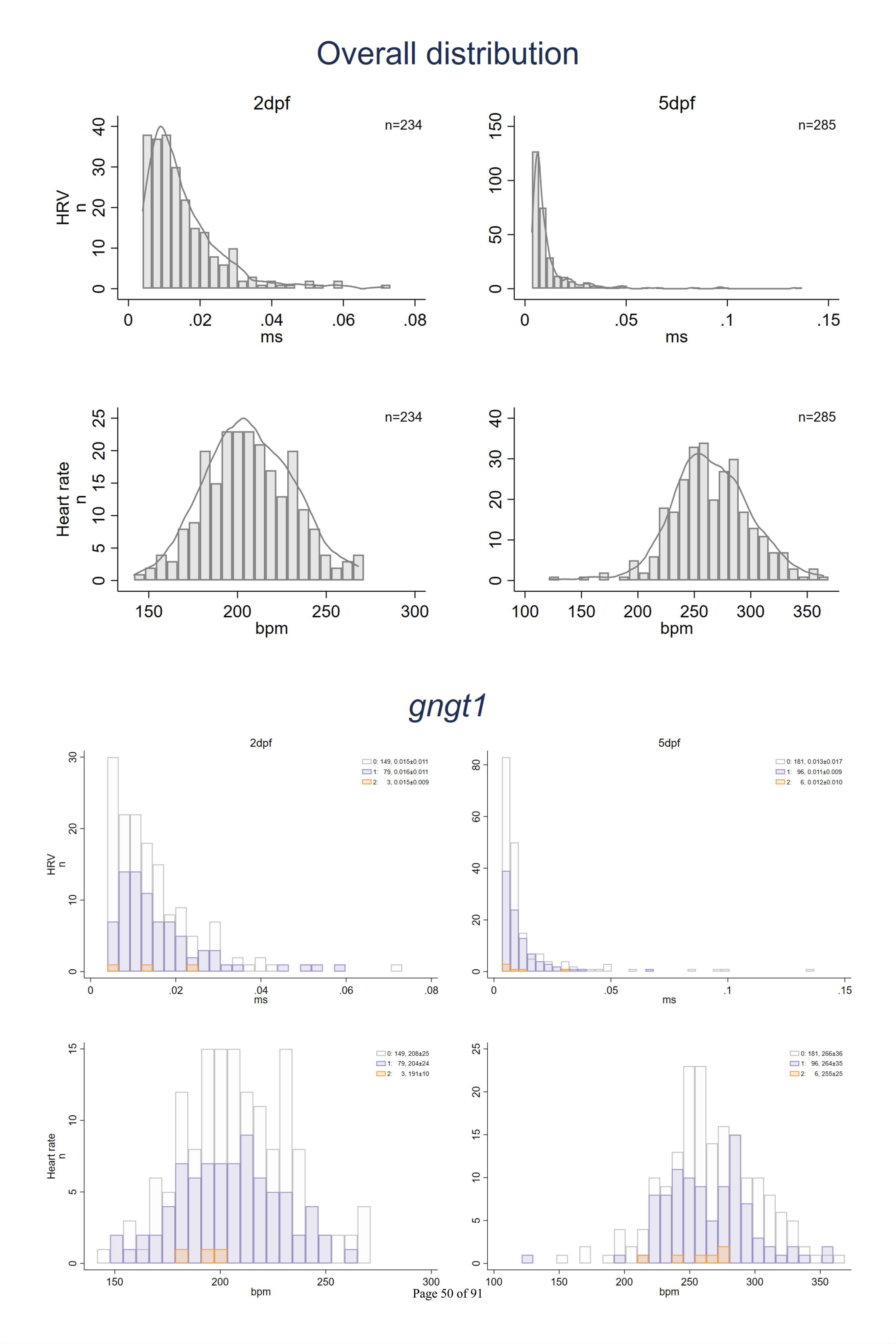
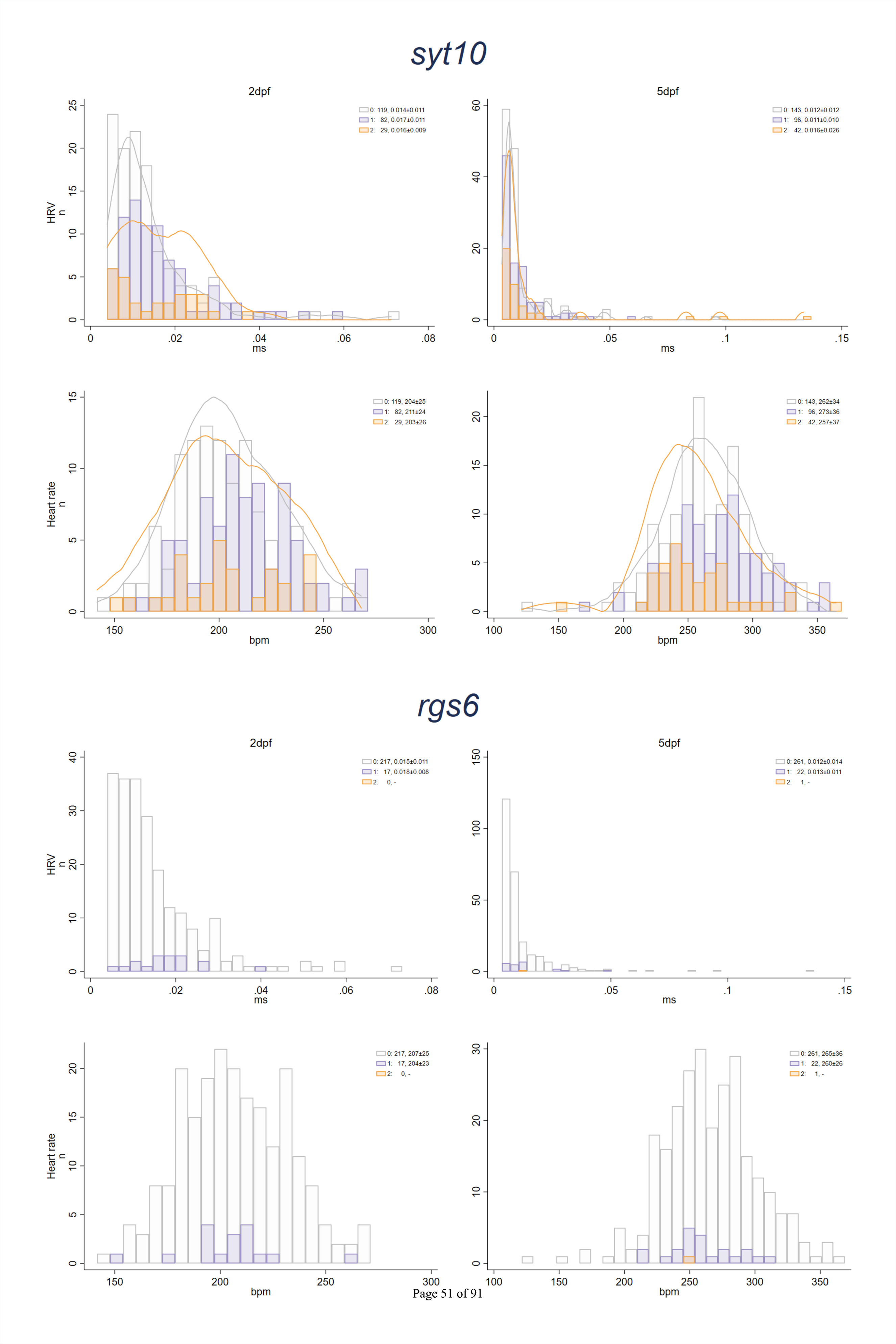
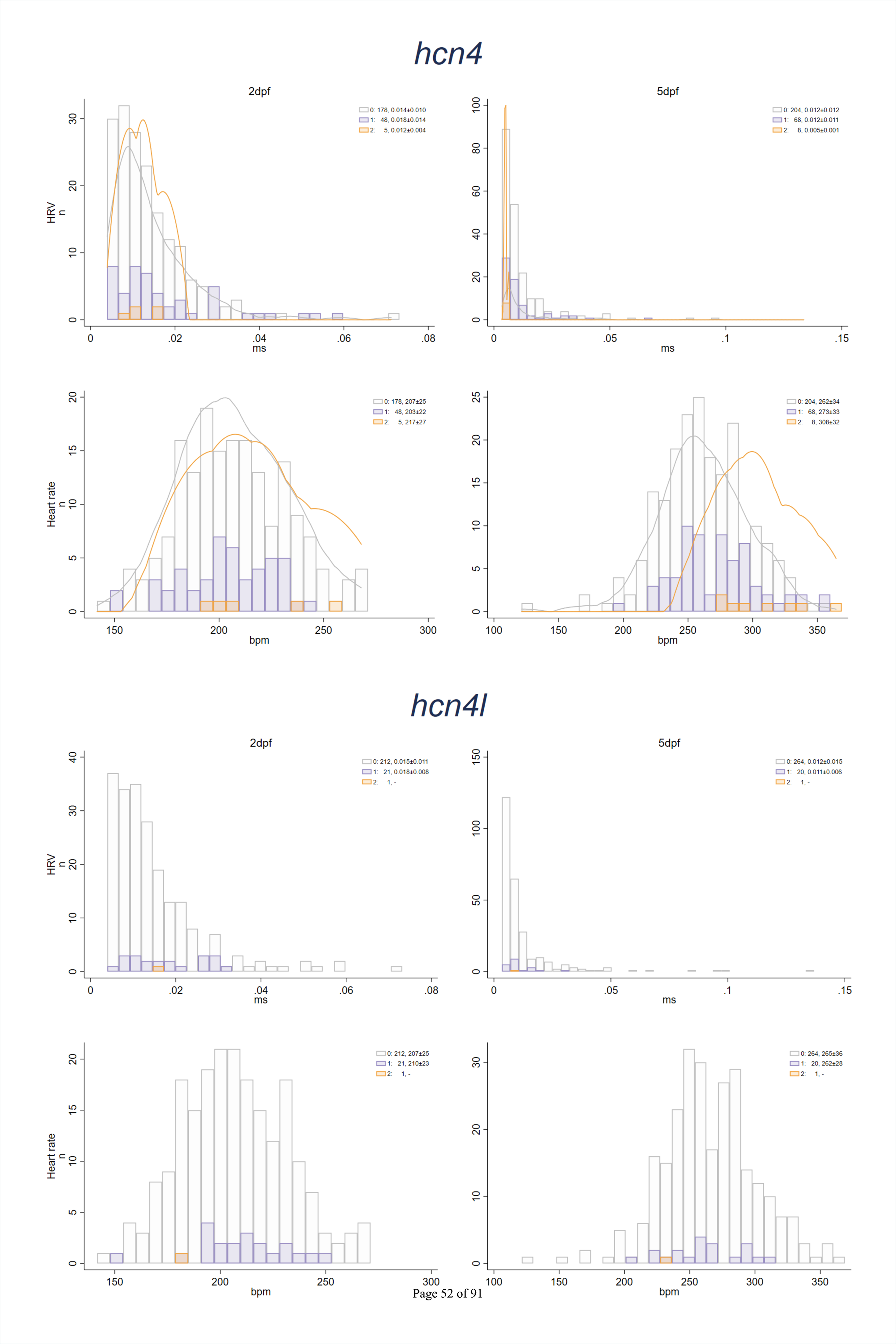
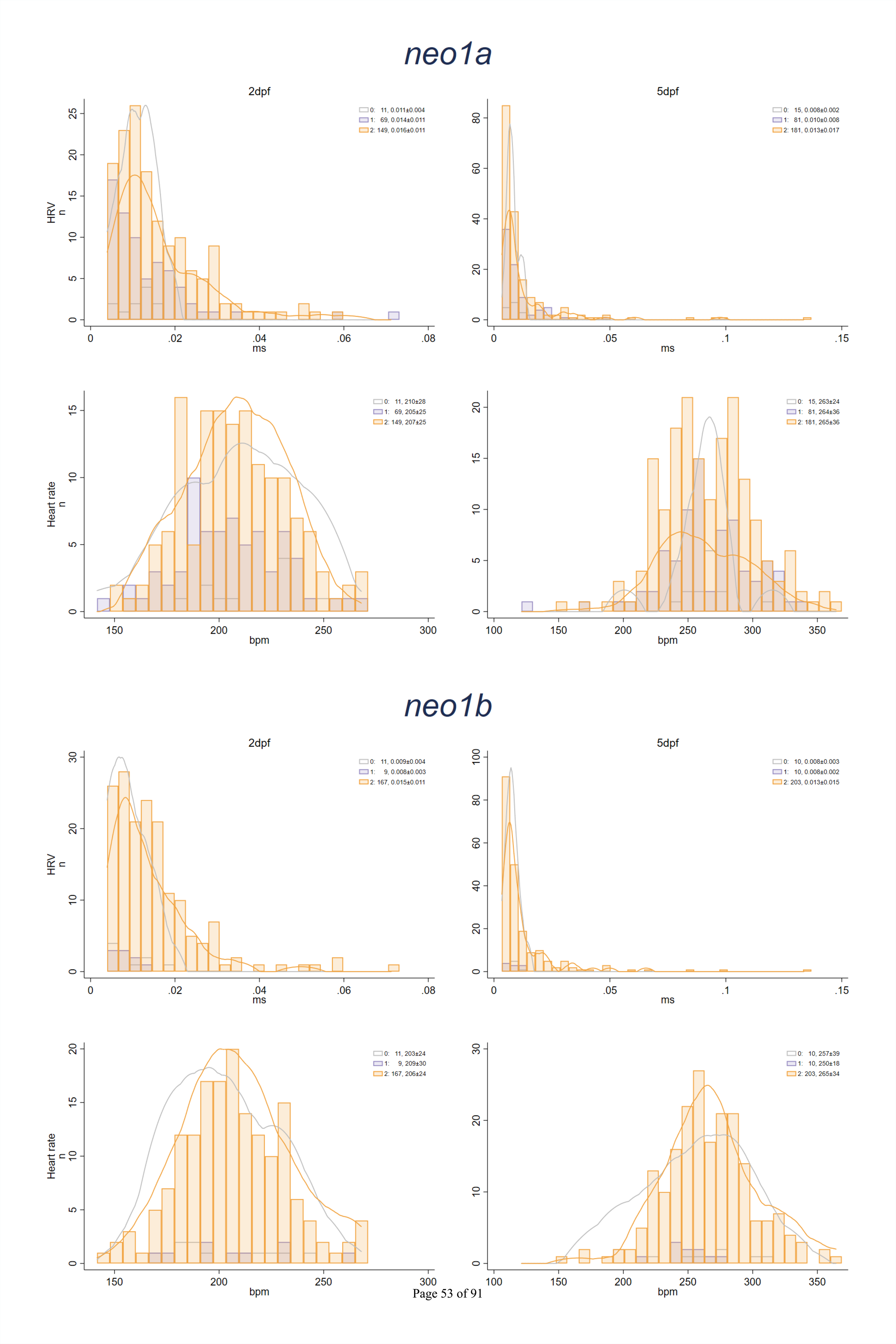
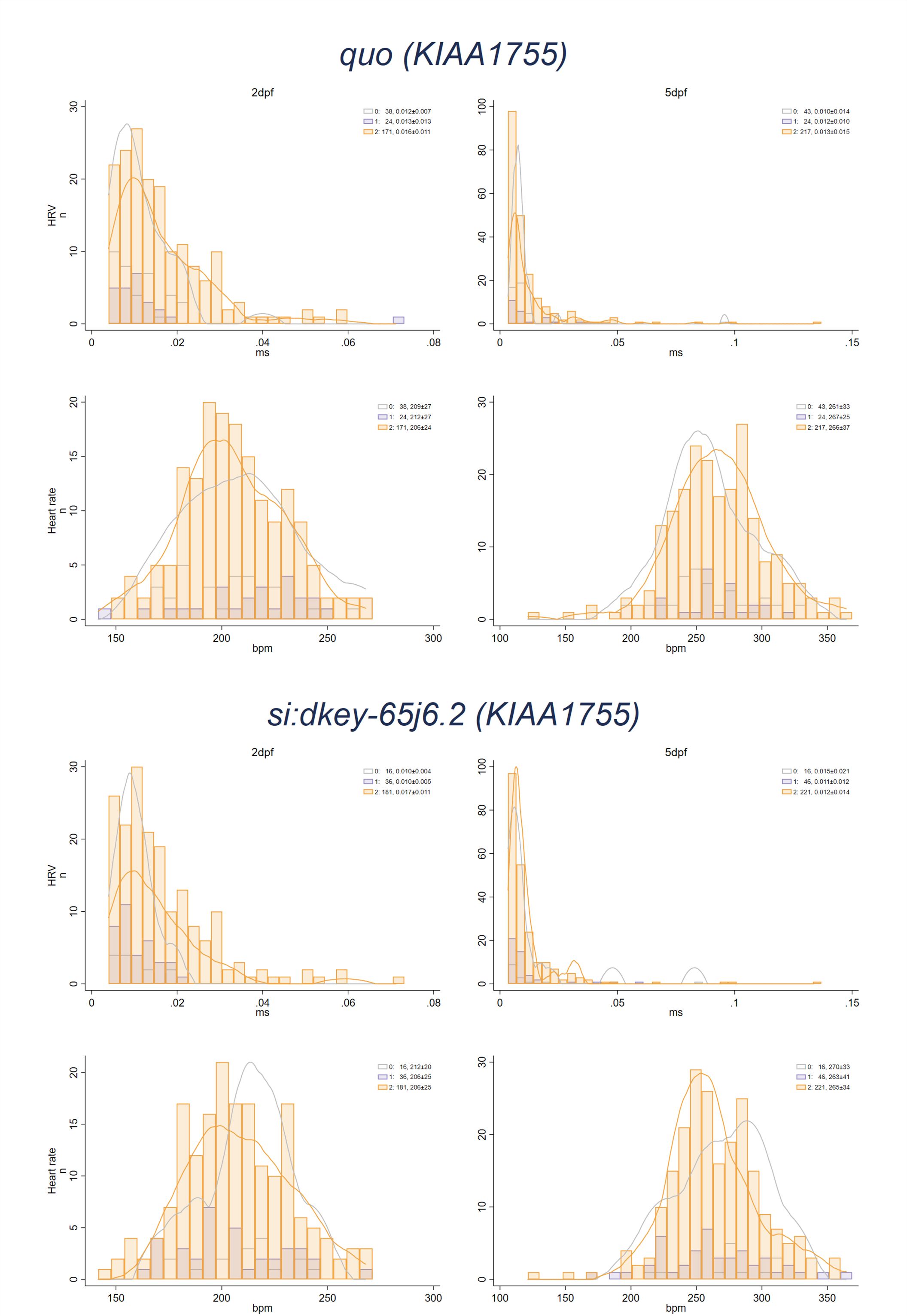
Distributions of heart rate variability (HRV, in ms) and heart rate (in beats per min, bpm) shown in all embryos combined, as well as by number of mutated alleles for each of the nine CRISPR/cas9 targeted candidate genes. In each histogram, the mean±SD in embryos with 0, 1 and 2 mutated alleles is shown in the top right corner. Orange and gray lines show Kernell density plots for embryos with CRISPR/Cas9-induced nonsense mutations in both alleles, and for embryos free from CRISPR/Cas9-in^Pa^d^ge 54^u^of 91^ced mutations, respectively, if n>5 for both.

**Supplementary Figure 4:**
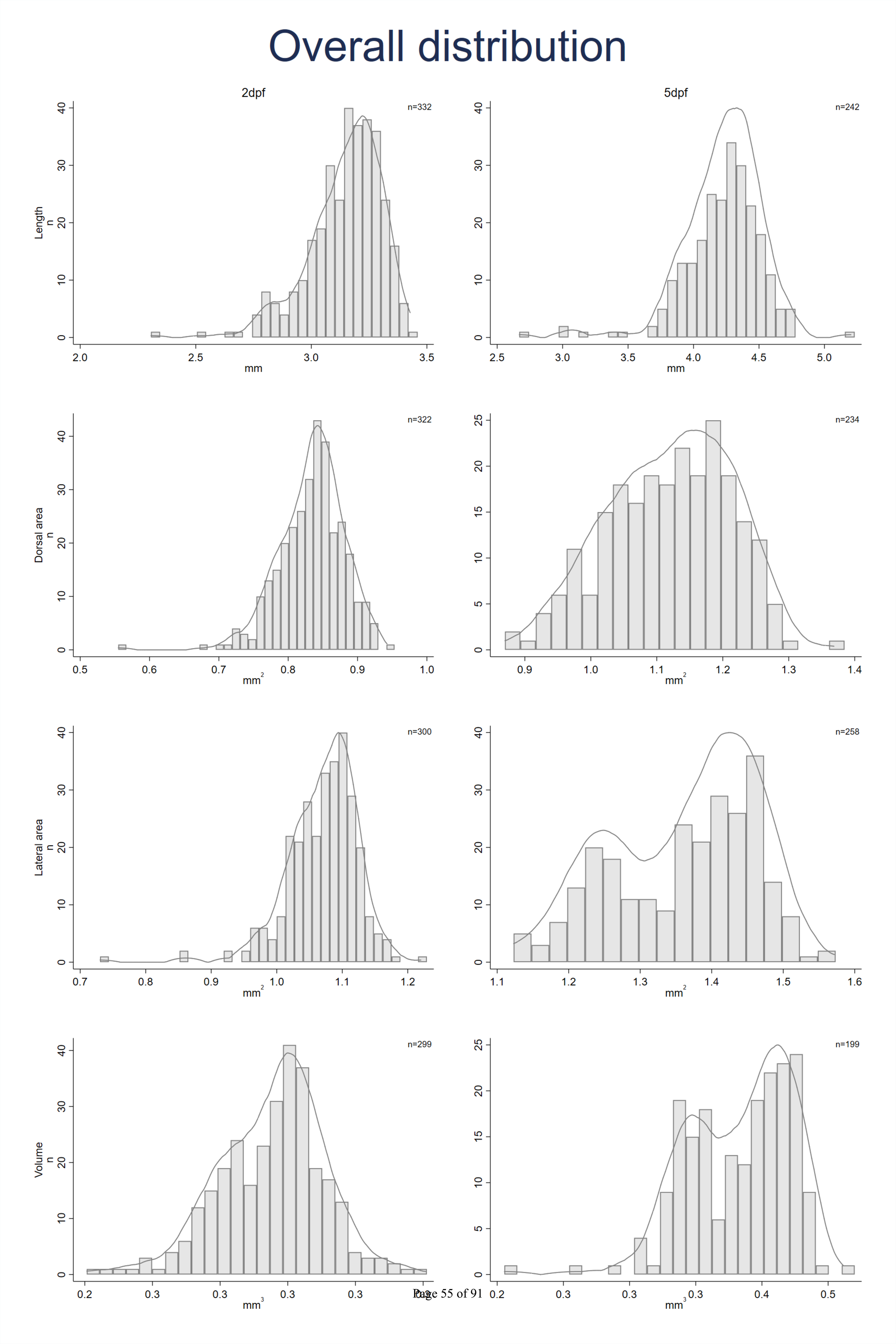
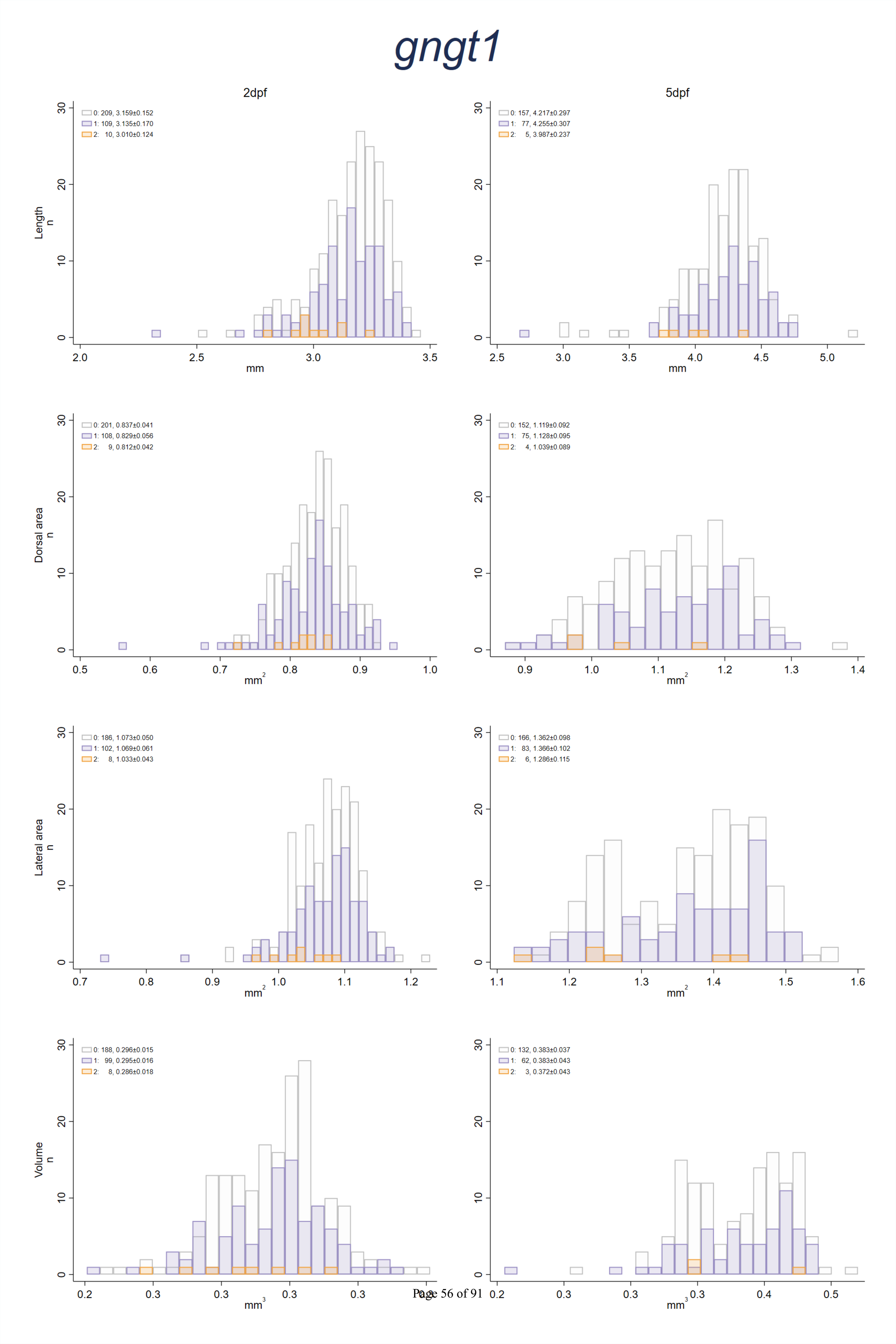
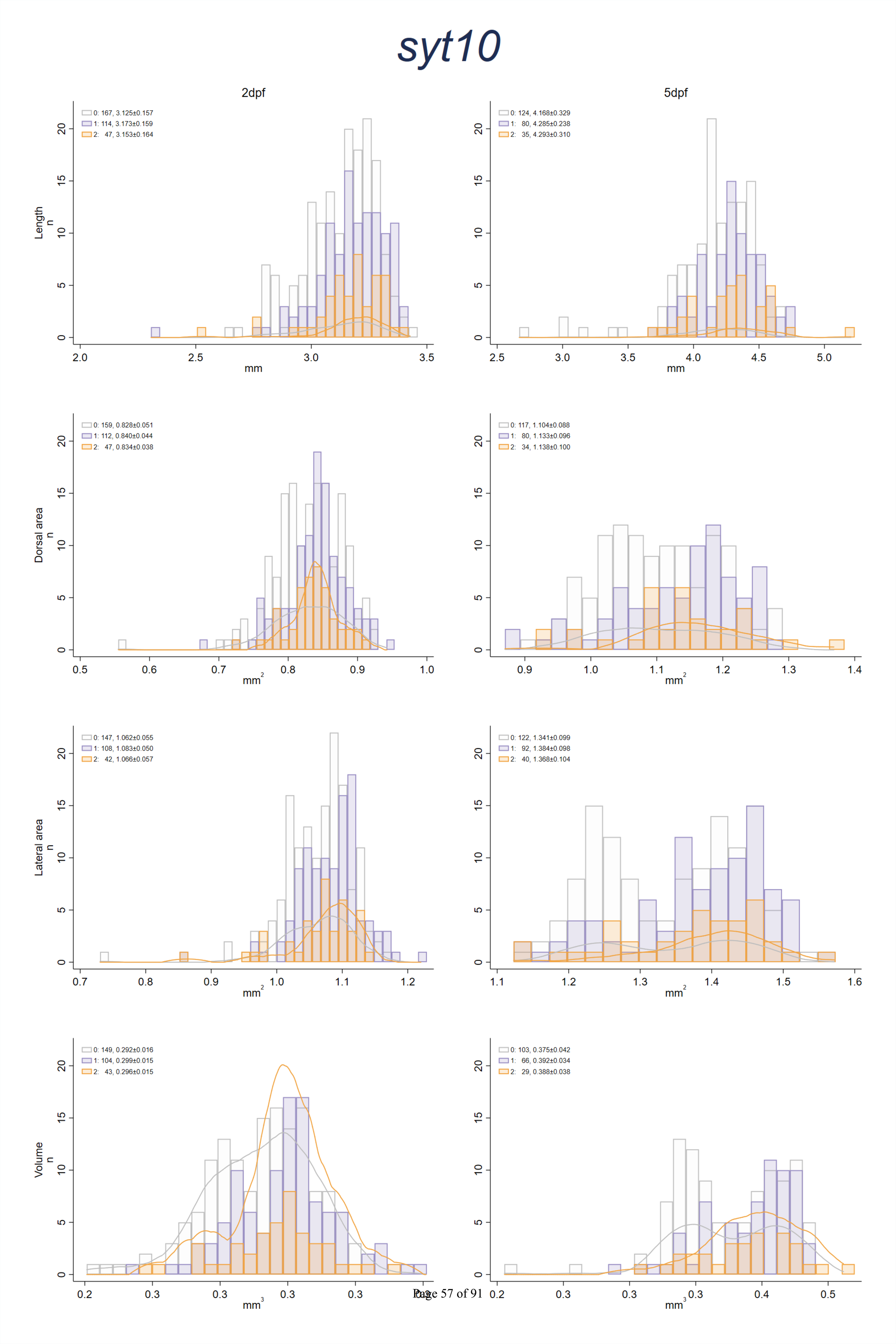
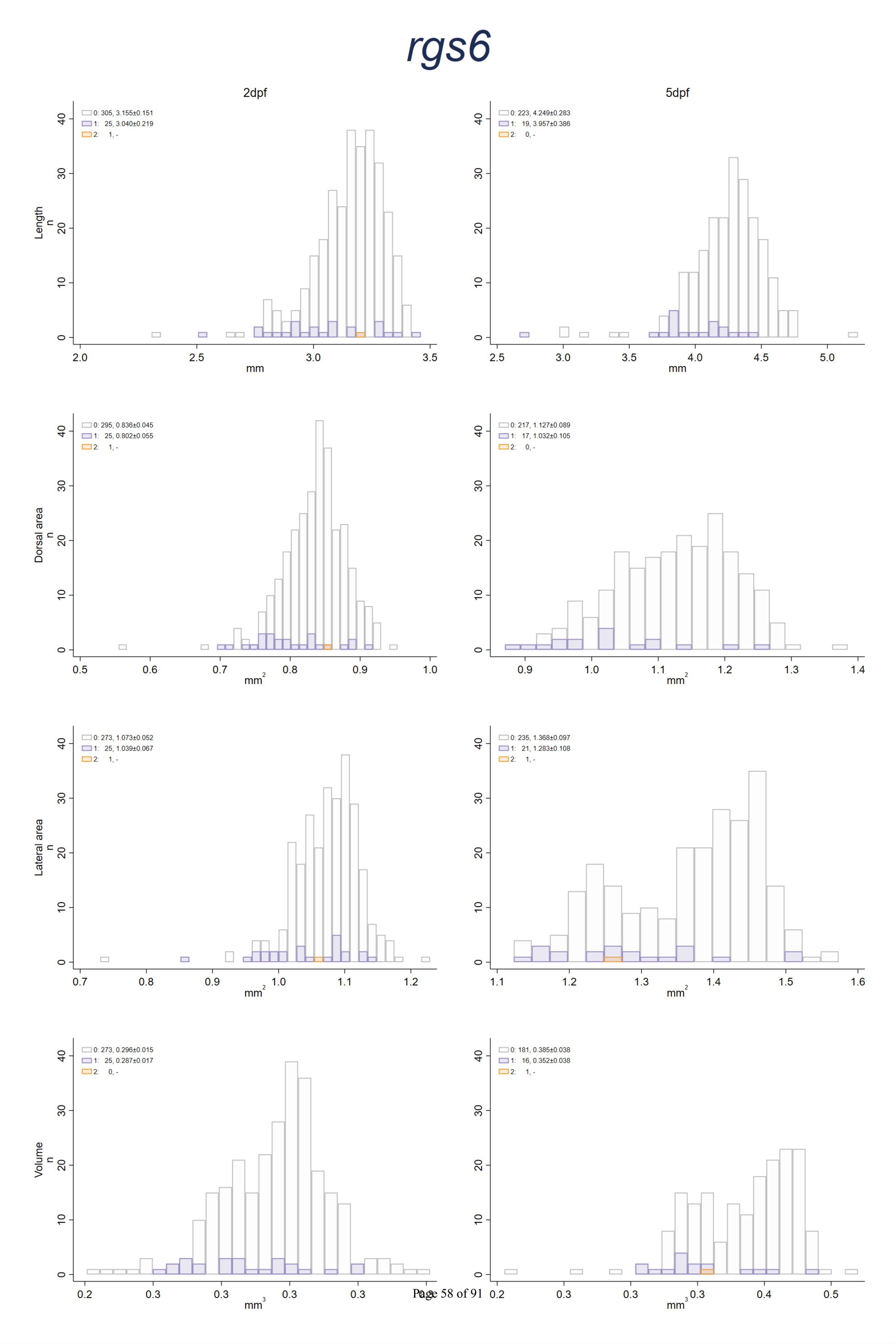
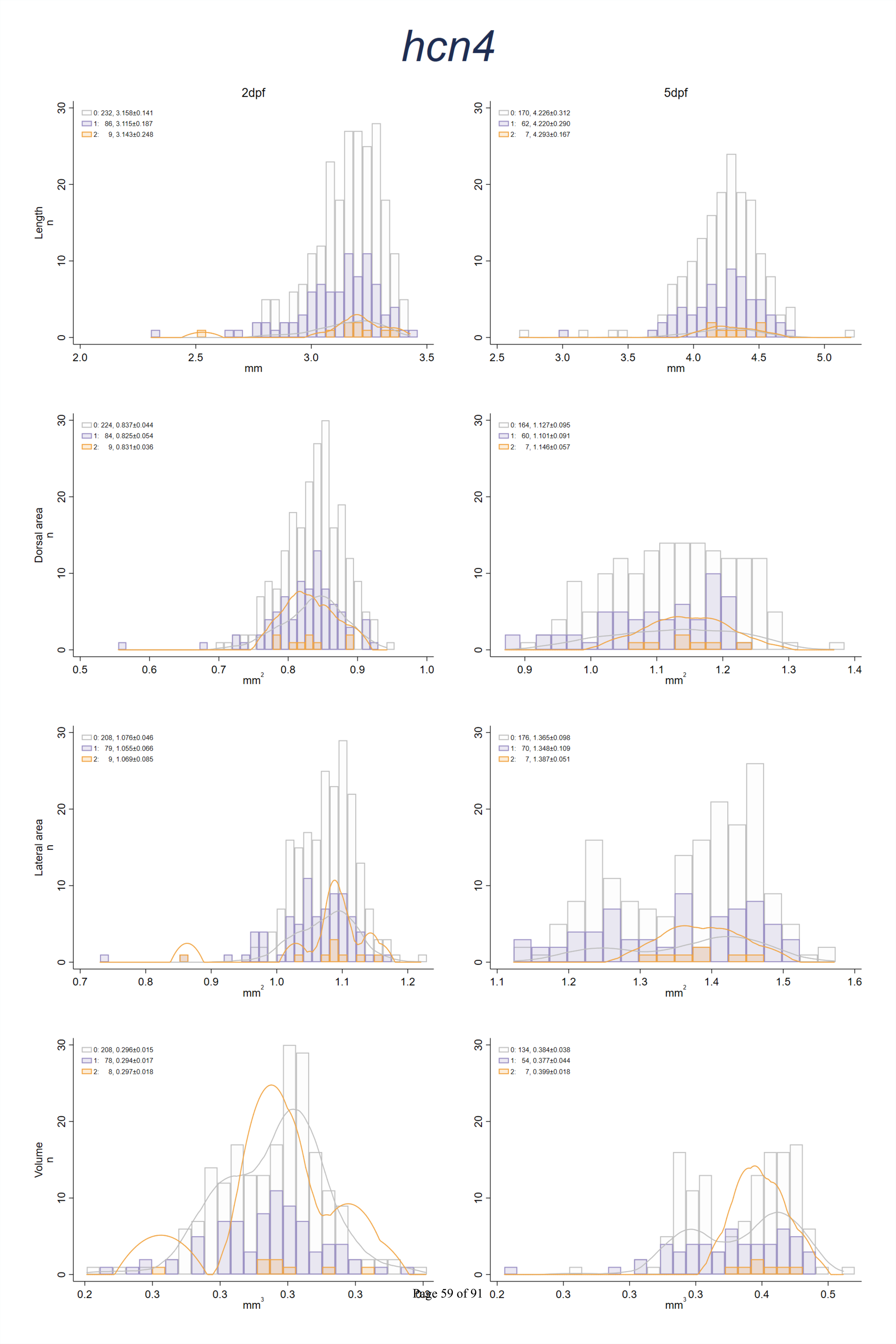
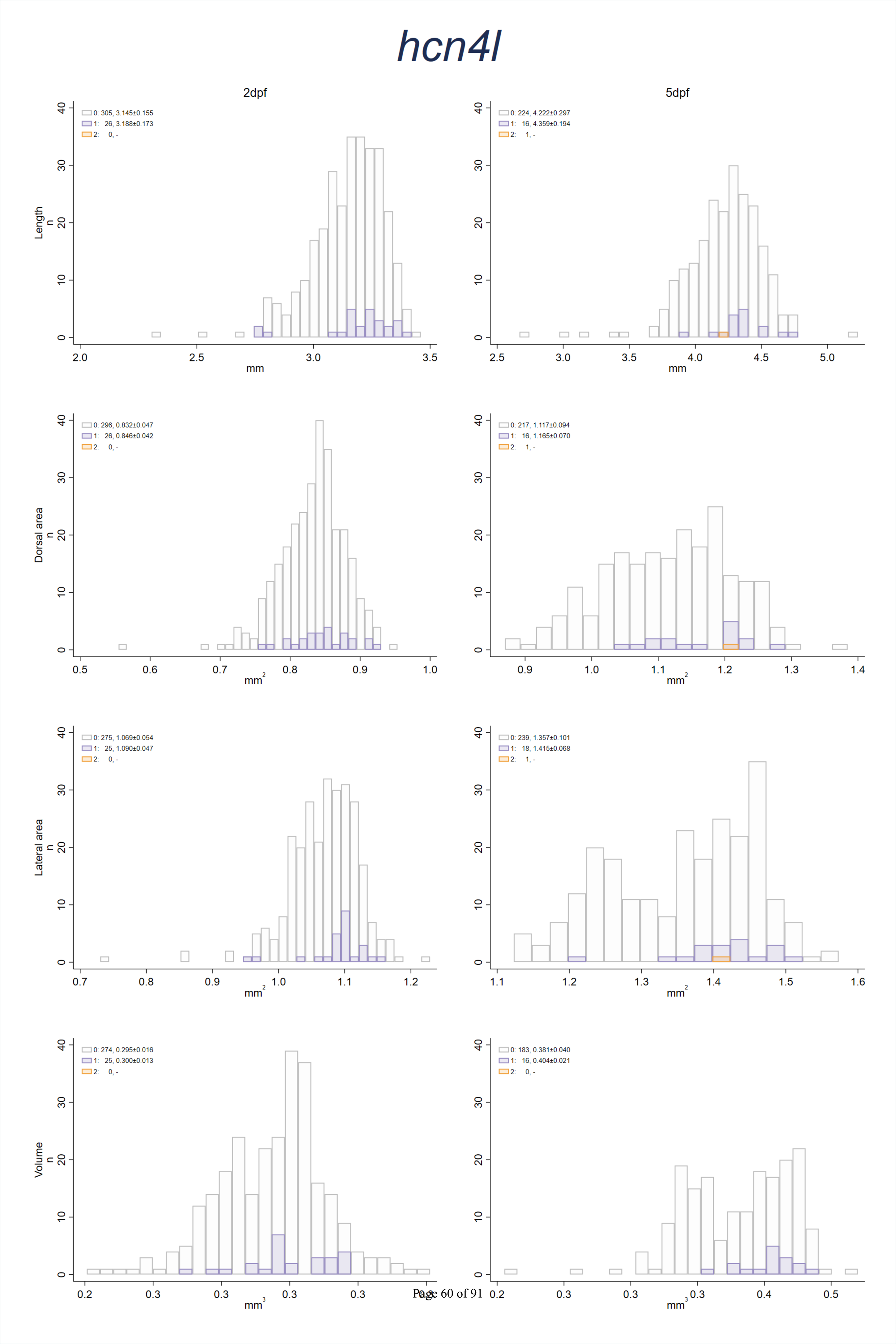
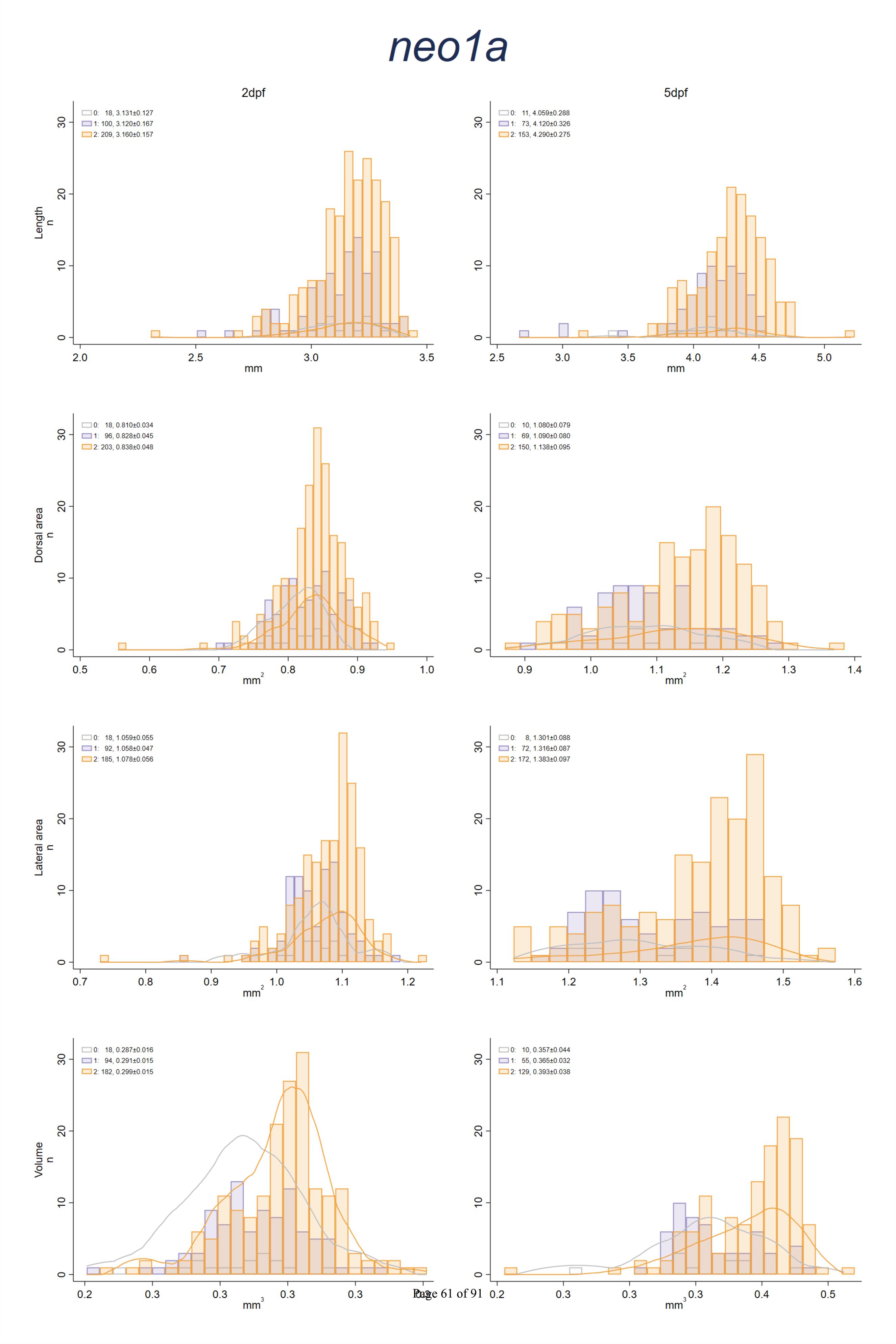
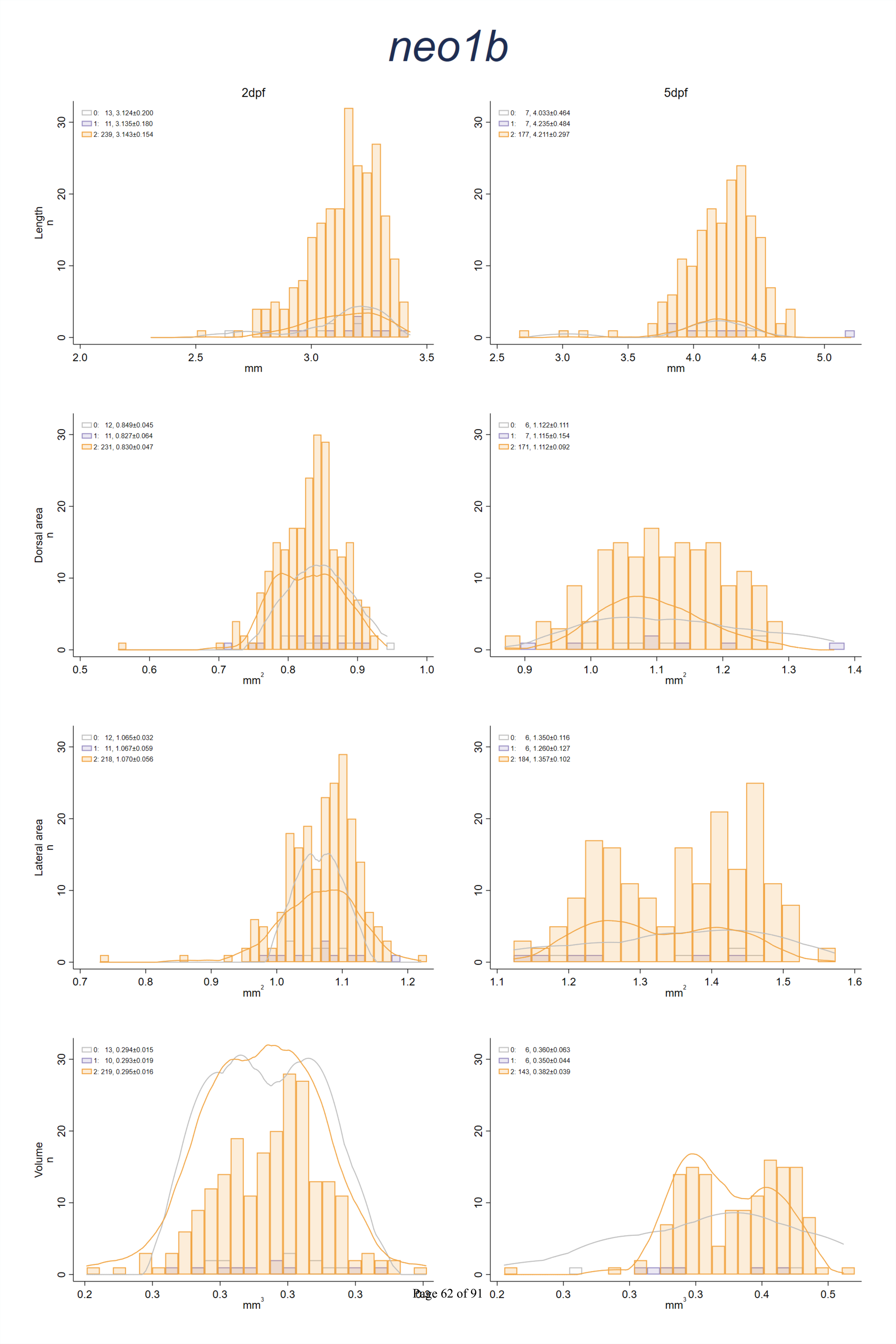
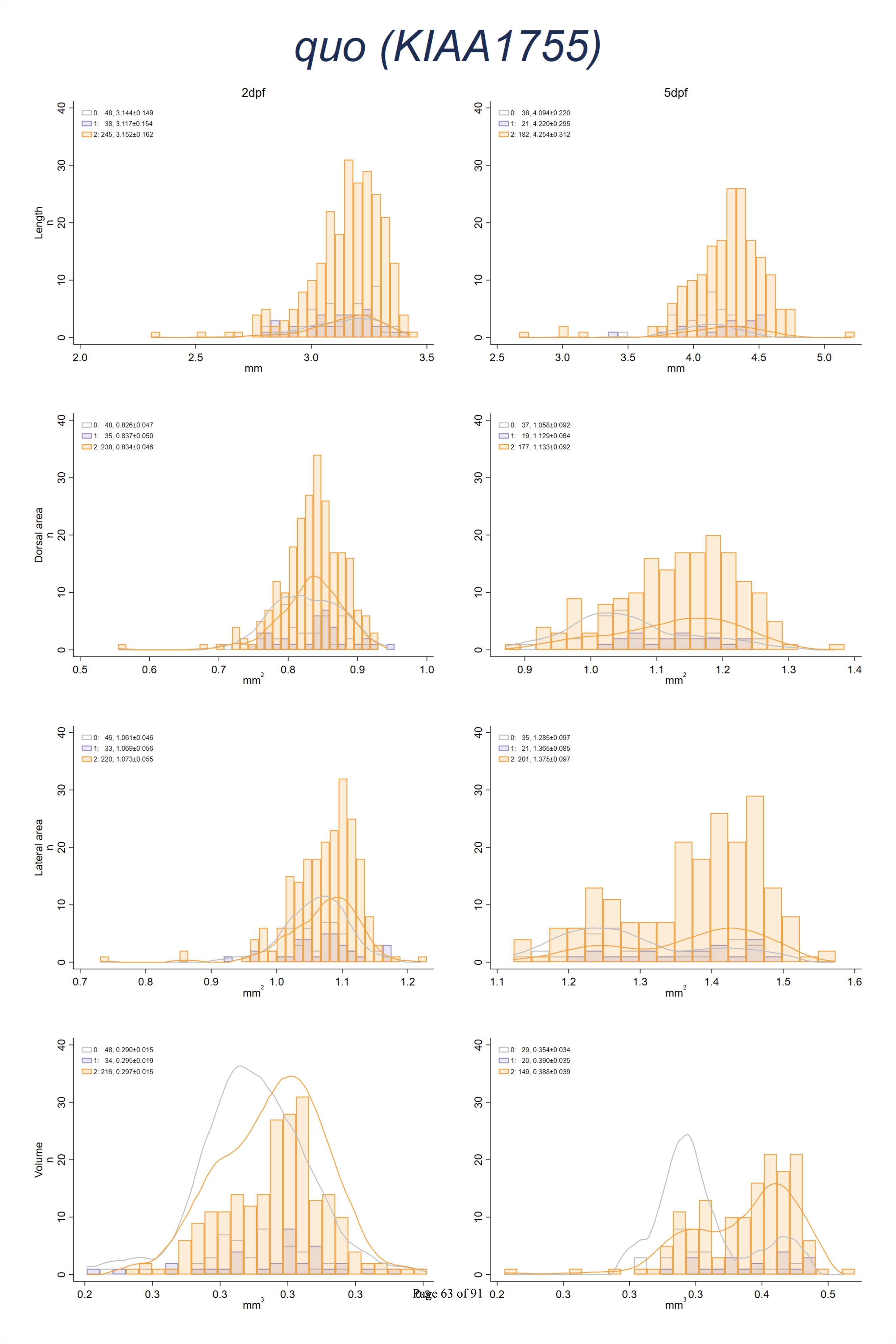
Distributions of body length, dorsal and lateral body surface area, and body volume shown in all embryos combined, as well as by number of mutated alleles for each of the nine CRISPR/cas9 targeted candidate genes. In each histogram, the mean±SD in embryos with 0, 1 and 2 mutated alleles is shown in the top left corner. Orange and gray lines show Kernell density plots for embryos with CRISPR/Cas9-induced nonsense mutations in both alleles, and for embryos free from CRISPR/Cas9-^P^i^ag^n^e64 o^d^f91^uced mutations, respectively, if n>5 for both.

**Supplementary Figure 5:**
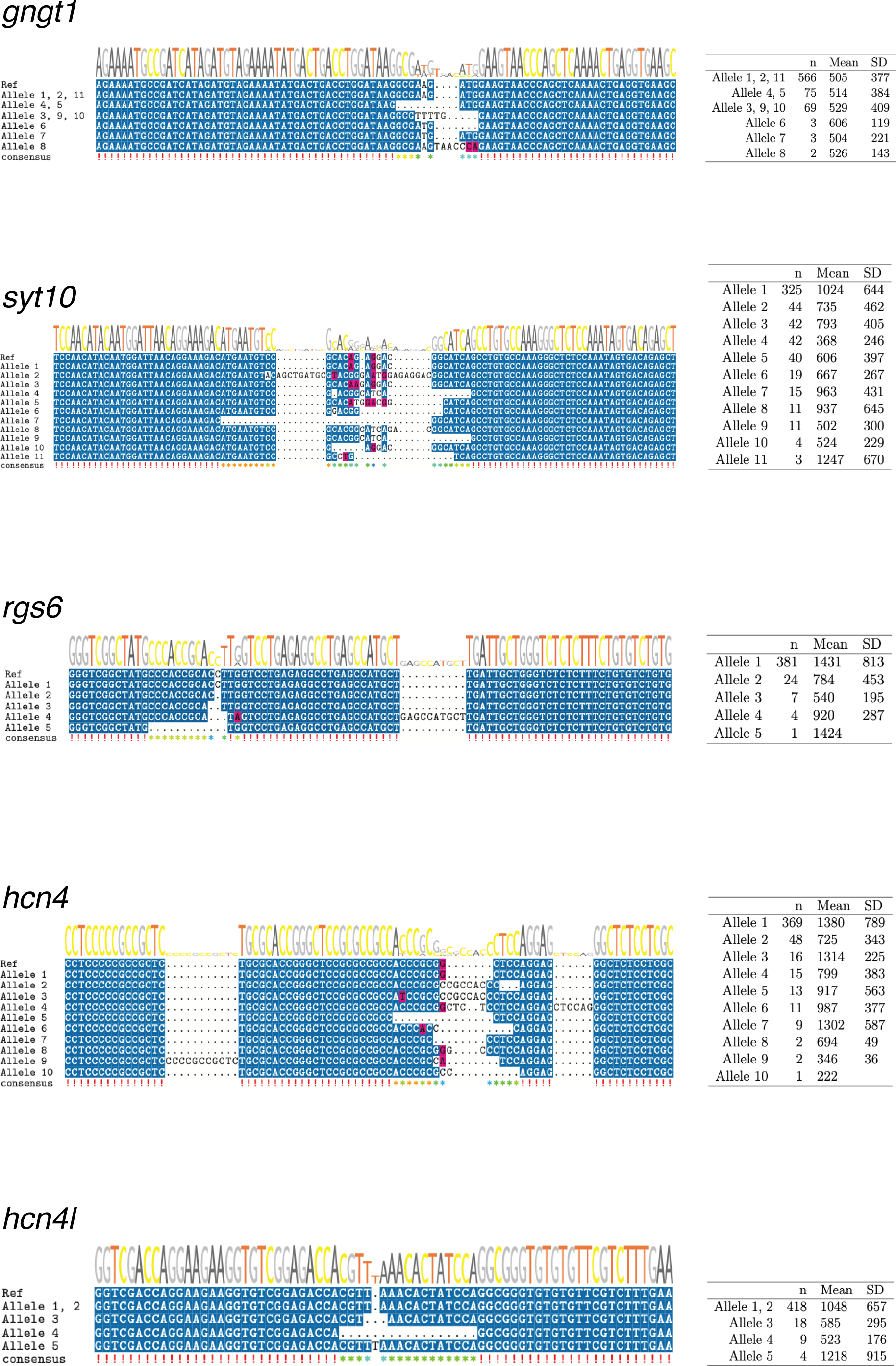
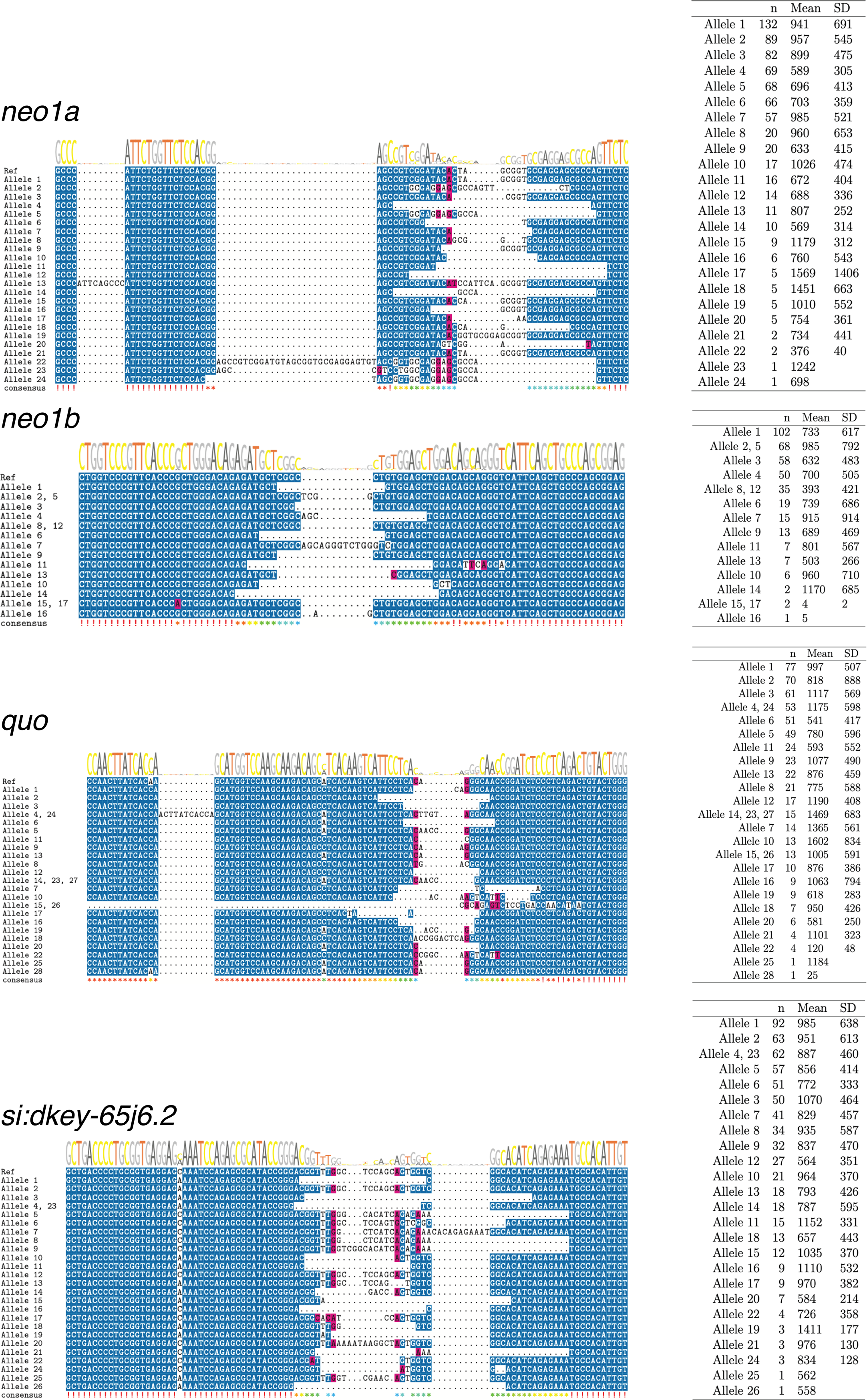
Unique alleles in nine candidate genes showing CRISPR/Cas9-induced mutations compared with the zebrafish reference genome, GRCz11. Colour coding illustrates if base pairs are ≥50% conserved (blue), similar (pink) or non-conserved (white). Tables on the right show the number times each allele appears across the 381 successfully sequenced embryos (max 2×381), and the mean and standard deviation for the number of reads calls are based on. Alleles that differ by inherently present variants not attributed to CRISPR/Cas9 are grouped together as indicated in the figure.

**Supplementary Figure 6:**
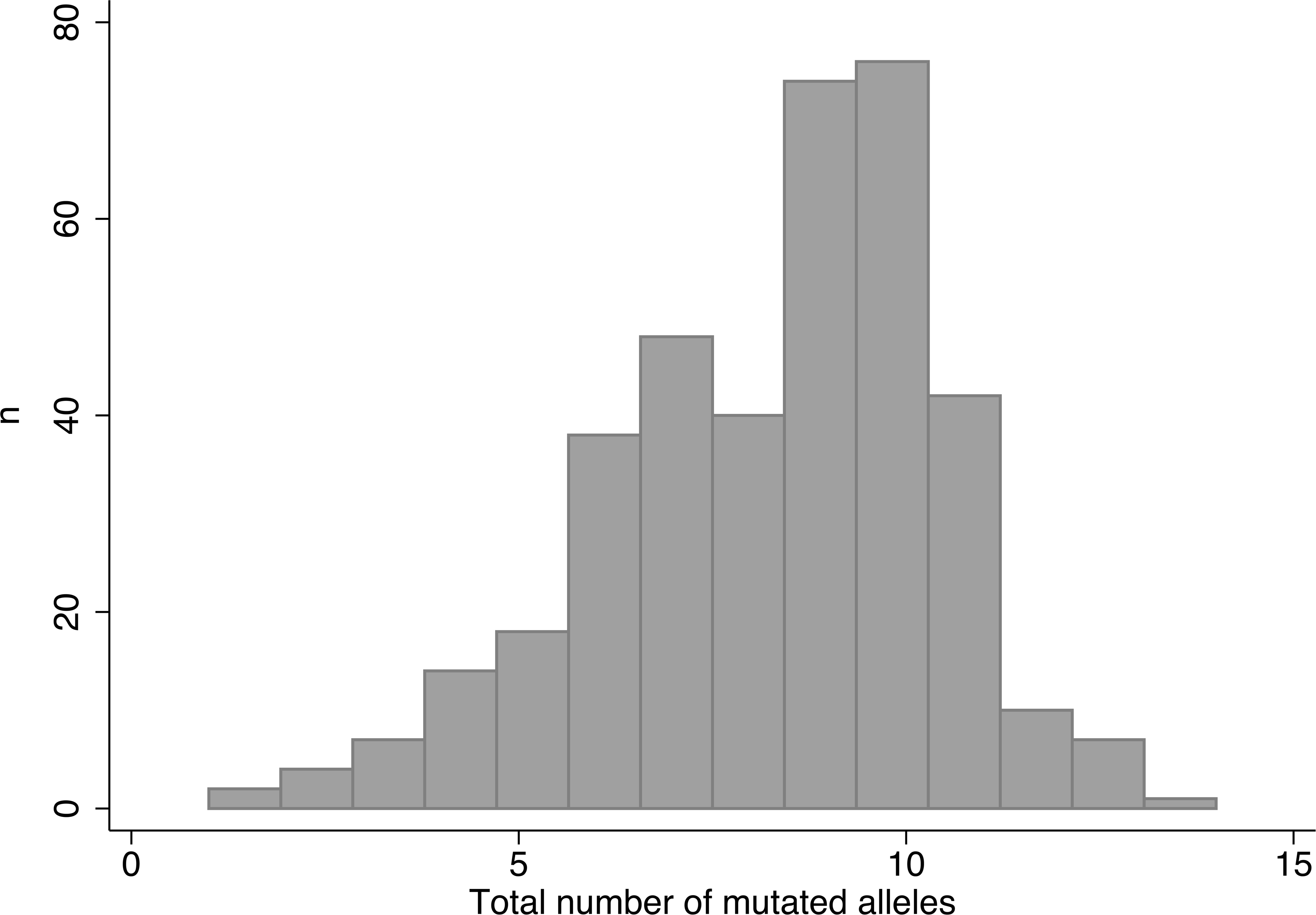
Distribution of the number of mutated alleles across the nine CRISPR-targeted sites. A mutation is defined as any previously undescribed variant lo^P^c^ag^a^e^t^6^e^7^d^of^w^91^ithin ±30bp of the CRISPR/Cas9 cut site.

**Supplementary Figure 7:**
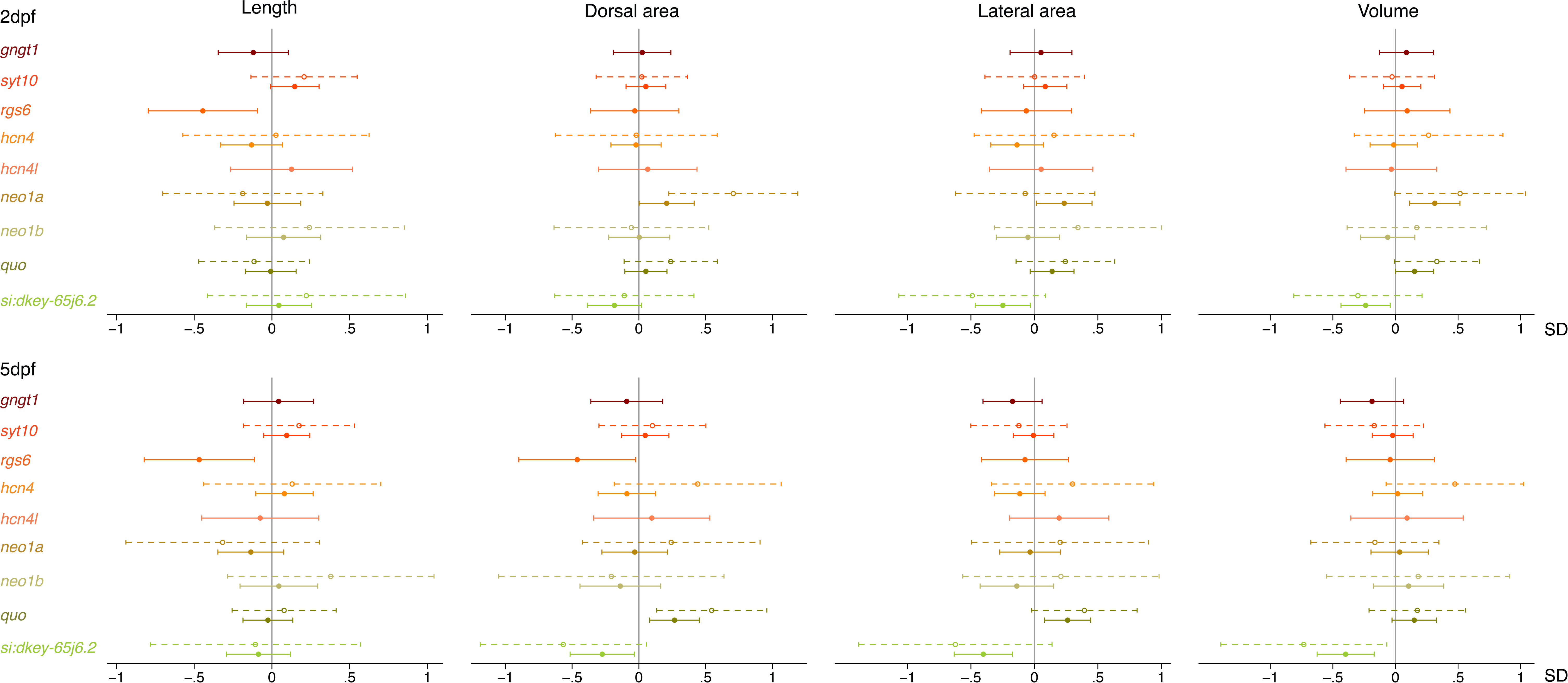
Effect of mutations in candidate genes on body length; and on dorsal body surface area, lateral body surface area and body volume normalised for length at 2 days post-fertilization (dpf, top) and 5dpf (bottom). Fulldots and solid whiskers show the effect size and 95% confidence interval for each additional mutated allele, weighted by the predicted effect on protein function. Open dots and dotted whiskers indicate the effect size and 95% confidence interval for frameshift or premature stop codon inducing mutations in two alleles vs. no CRISPR-induced mutations. Effects were adjusted for the weighted number of mutated alleles in the other genes, a^P^s^ag^w^e 6^e^8^l°l^f^a^91^s for time of day (fixed factors), with embryos nested in batches (random factor). *quo* and *si:dkey-65j6.2* are orthologues of the human *KIAA1755*.

**Supplementary Figure 8:**
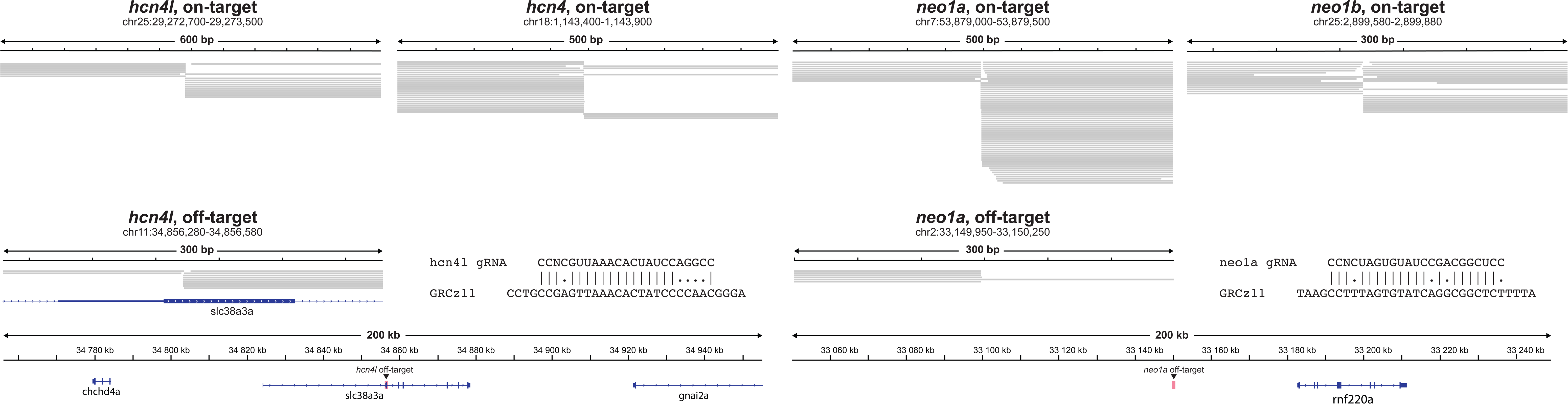
*In vitro* Nano off-target sequencing results for the *hcn4*, *hcn*^P^*4*^a^*l*^g^,^e^*n*^6^*e*^9^*o*°*1*^f^*a*^91^and *neo1b* gRNAs in DNA extracted from a Tg(*acta2*:GFP) positive fish used to generate CRISPR/Cas9 founders. One off-target site was observed for the *hcn4l* and *neo1a* gRNAs, in spite of five and four mismatches, respectively. For *hcn4l,* this off-target site was in the coding region of *slc38a3a*. Off-target mutagenic activity was lower than on-target activity for both gRNAs.

**Supplementary Figure 9:**
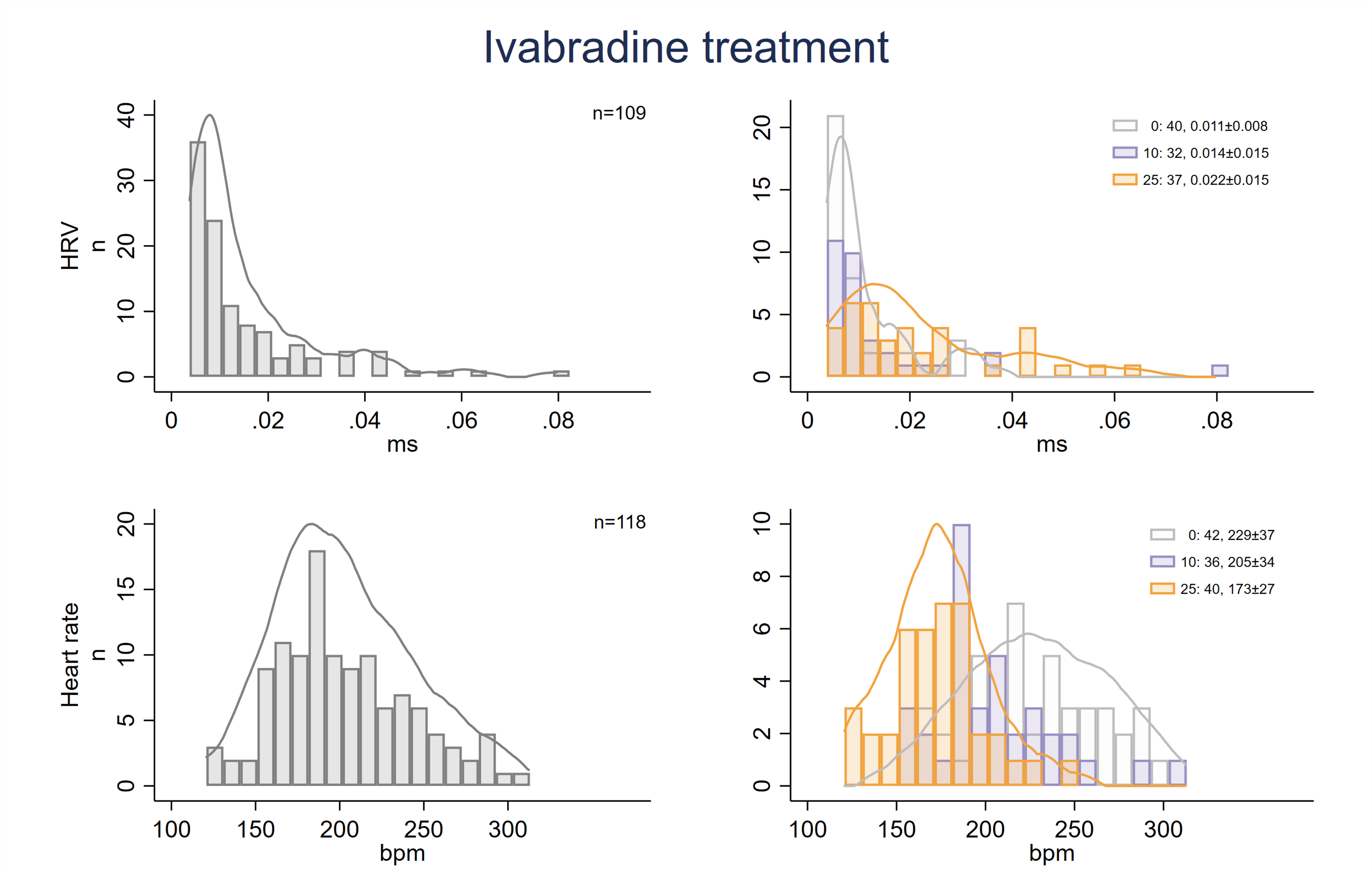
Distributions of heart rate variability (HRV, in ms) and heart rate (in beats per min, bpm) shown in all embryos combined (left), as well as by ivabradine treatment condition (right). In each histogram, the mean±SD in embryos treated with 0, 10 and 25µM ivabradine is shown in the top right corner. Orange and gray lines show Kernel density plots for embryos treated with 25 and 0 µM ivabrad_Page 7_i_0_n_of 91_e, respectively.

## SUPPLEMENTARY TABLES

**Supplementary Table 1.** Overview of zebrafish orthologues of human candidate genes, CRISPR/Cas9 guide RNAs, and predicted off-targets

| Human gene | ENSG Stable ID | Zebrafish orthologue | ENSDARG stable ID | Target %id | Query %id | Conservation<br>Zebrafish - Human | Conservation Zebrafish -<br>Spotted gar | Main zebrafish<br>transcript |
| --- | --- | --- | --- | --- | --- | --- | --- | --- |
| *GNGT1* /<br>*GNG11* | ENSG00000127928 /<br>ENSG00000127920 | *gngt1* | ENSDARG00000035798 | 67.12 / 61.64 | 66.22 / 61.64 | 17 | 21 | ENSDART00000051950 |
| *SYT10* | ENSG00000110975 | *syt10* | ENSDARG00000045750 | 64.55 | 75.53 | 0 | 1 | ENSDART00000136049 |
| *RGS6* |  | *rgs6* | ENSDARG00000015627 | 82.21 | 82.04 | 9 | 20 | ENSDART00000135513 |
| *HCN4* | ENSG00000138622 | *hcn4* | ENSDARG00000061685 | 63.1 | 58.27 | 6 | 3 | ENSDART00000136140 |
| *HCN4* | ENSG00000138622 | *hcn4l* | ENSDARG00000074419 | 59.76 | 49.88 | 3 | 3 | ENSDART00000088249 |
| *NEO1* | ENSG00000067141 | *neo1a* | ENSDARG00000102855 | 70.45 | 68.86 | 1 | 5 | ENSDART00000158160 |
| *NEO1* | ENSG00000067141 | *neo1b* | ENSDARG00000075100 | 64.26 | 62.15 | 2 | 6 | ENSDART00000115280 |
| *KIAA1755* | ENSG00000149633 | *quo* | ENSDARG00000073684 |  |  | 2 | 5 | ENSDART00000109679 |
| *KIAA1755* | ENSG00000149633 | *si:dkey-65j6.2* | ENSDARG00000103586 |  |  | 3 | 5 | ENSDART00000158832 |

Table showing zebrafish orthologues of human candidate genes and their main transcript. Target %id is the percentage of the orthologous sequence that matches the human sequence (Ensembl); Query %id is the percentage of the human sequence matching the sequence of the orthologue; Conservation shows the number of genes in the locus that are preserved in the same locus across species according to Genomicus.

**Supplementary Table 2.**
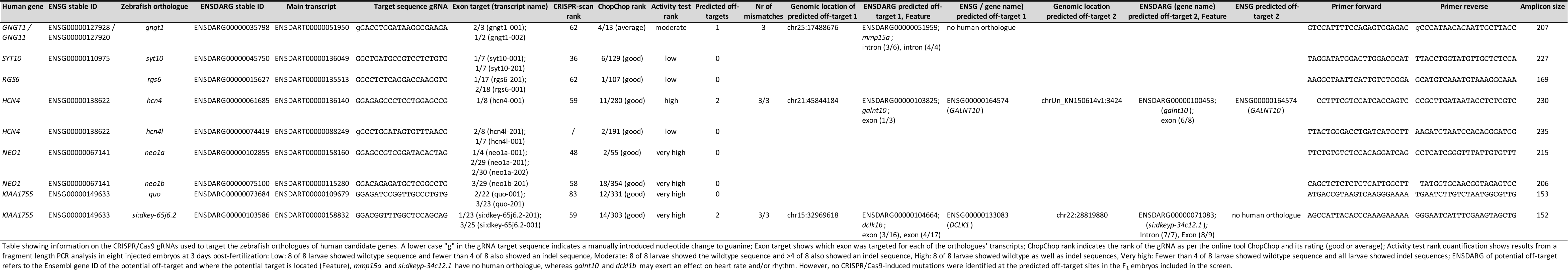
Overview of CRISPR/Cas9 guide RNAs

**Supplementary Table 3.**
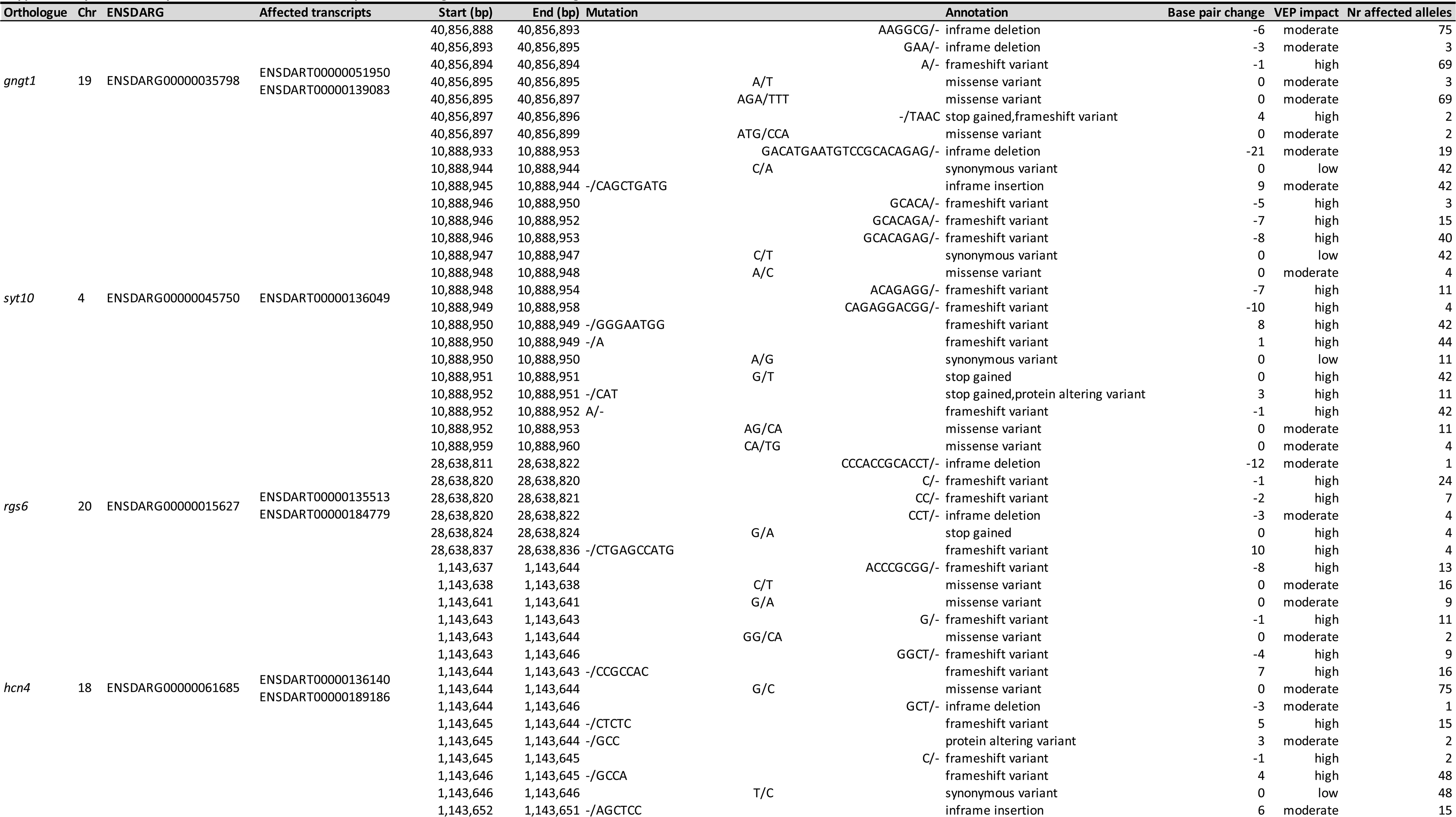
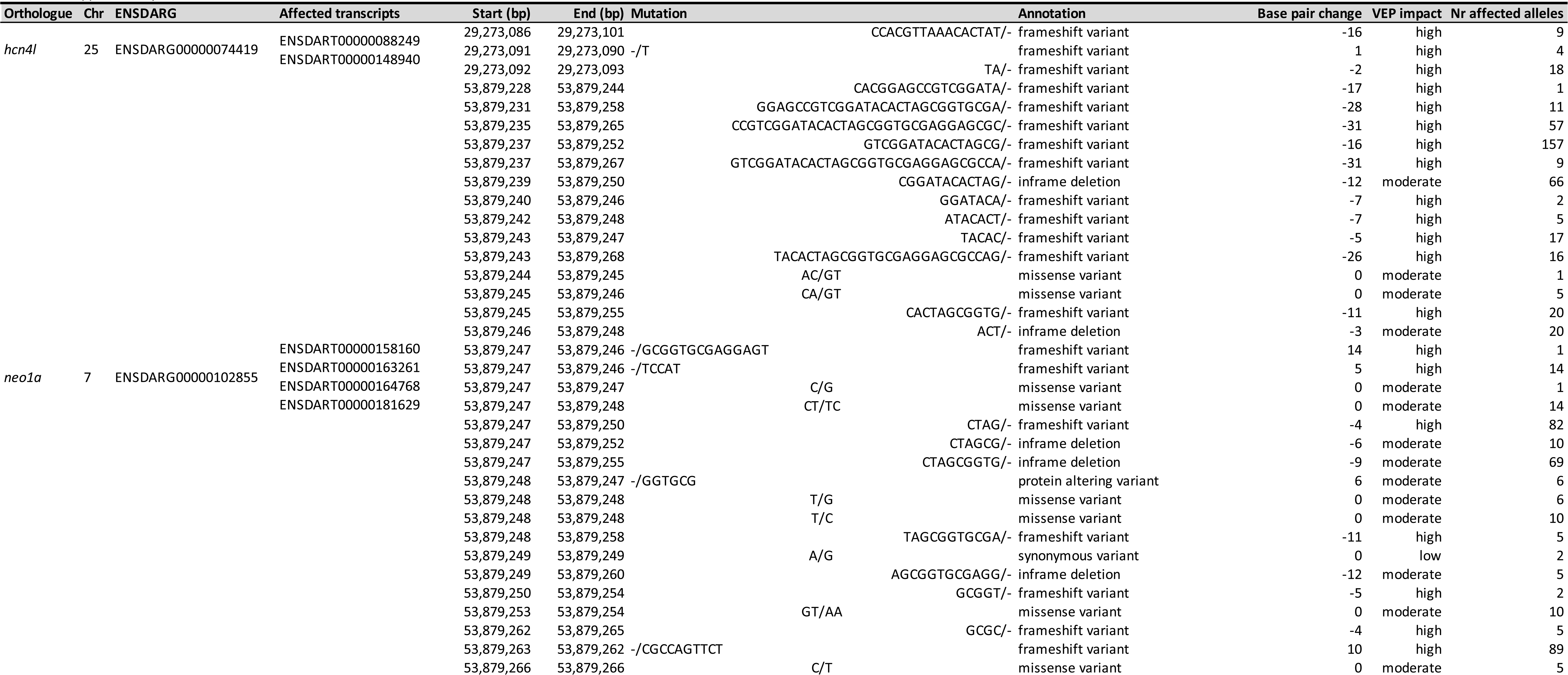
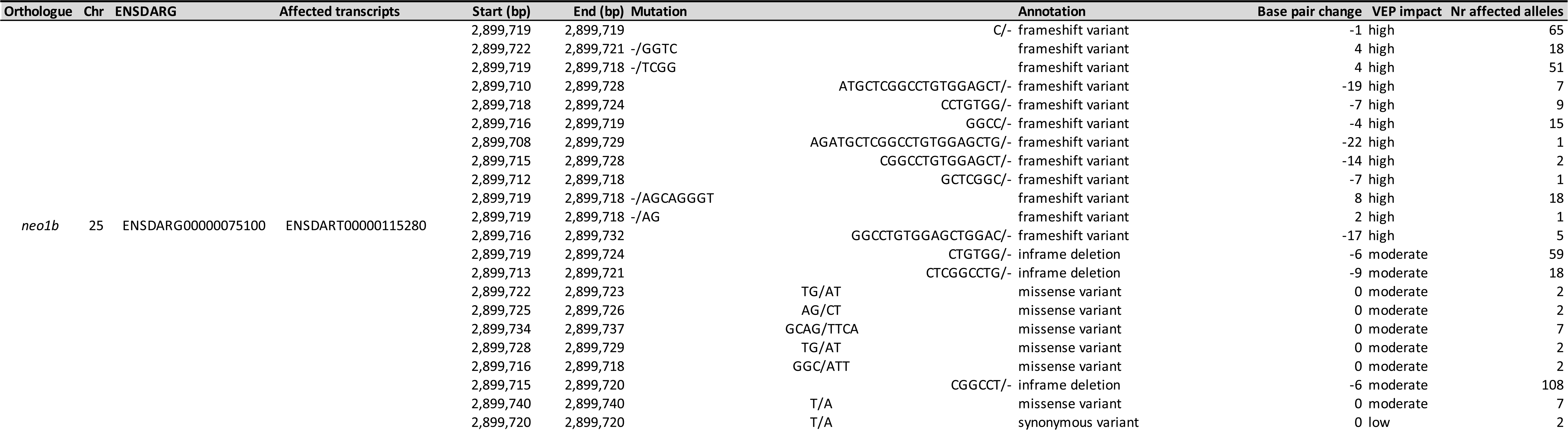
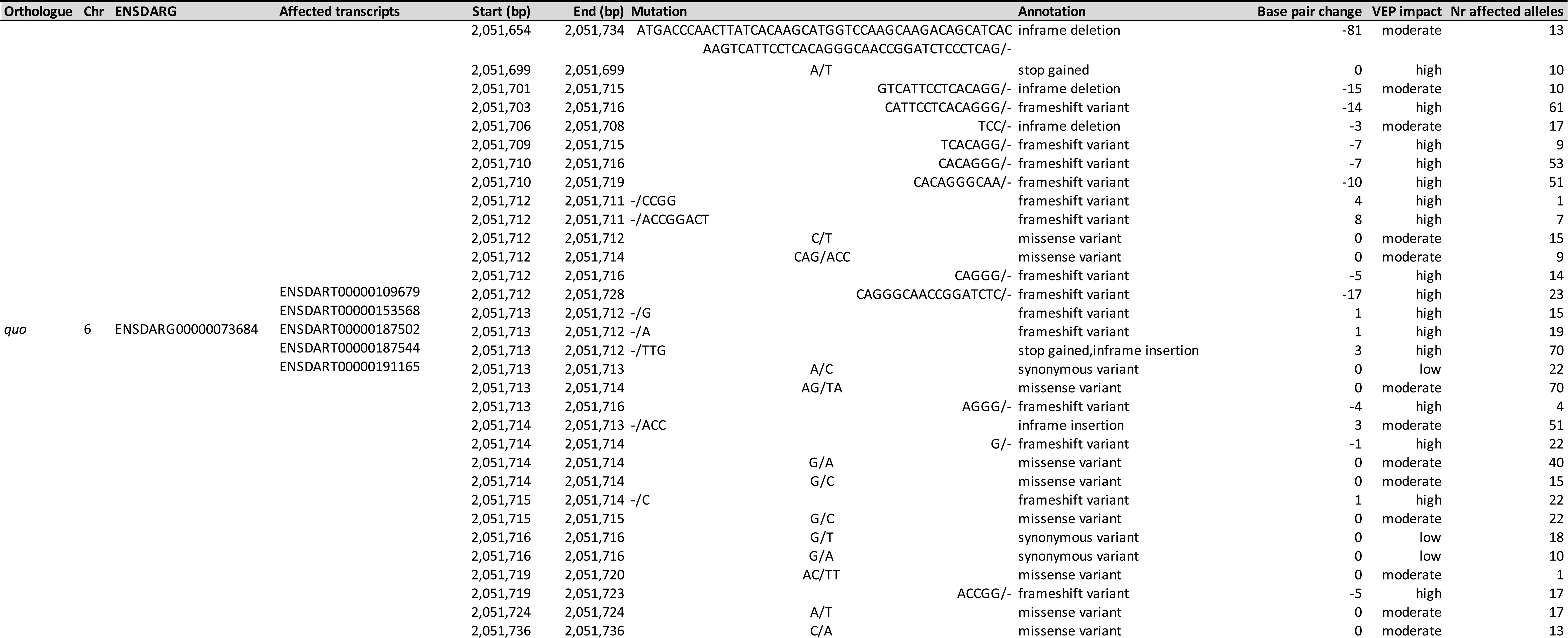
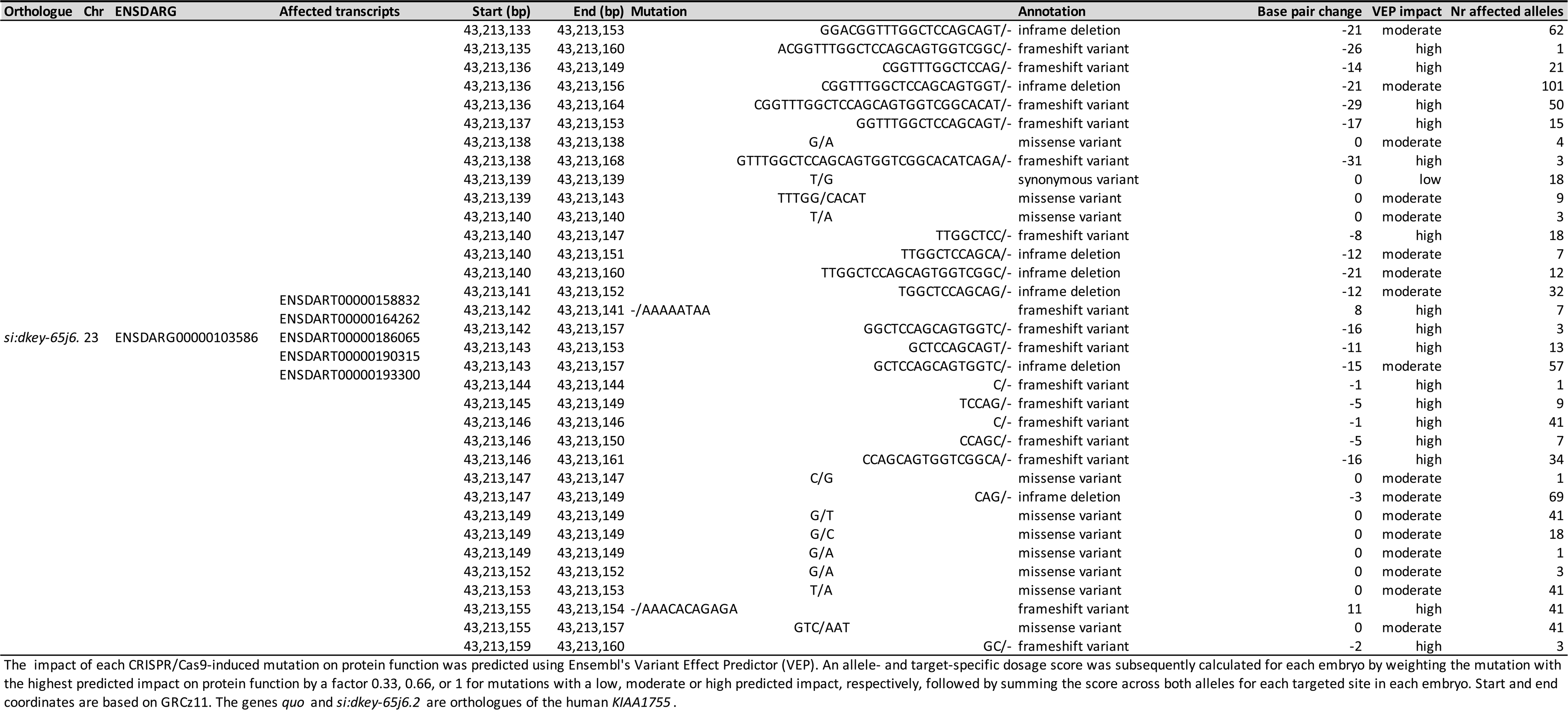
Unique variants and affected transcripts for each targeted zebrafish orthologue

**Supplementary Table 4.** Mutant allele frequencies for CRISPR/Cas9-induced mutations by zebrafish orthologue

| Gene | Nr mutated alleles |  |  | 2* | Missing | Sample call rate | Mutant allele freq | Expected nr mutated alleles |  |  | P<sub>HWE</sub> |
| --- | --- | --- | --- | --- | --- | --- | --- | --- | --- | --- | --- |
|  | 0 | 1 | 2 |  |  |  |  | 0 | 1 | 2 |  |
| *gngt1* | 234 | 130 | 12 | 2 | 5 | 0.987 | 0.205 | 238 | 122 | 16 | 2.33E-01 |
| *syt10* | 192 | 130 | 54 | 46 | 5 | 0.987 | 0.316 | 176 | 163 | 38 | 9.82E-05 |
| *rgs6* | 344 | 34 | 1 | 1 | 2 | 0.995 | 0.047 | 344 | 34 | 1 | 8.69E-01 |
| *hcn4* | 266 | 100 | 9 | 9 | 6 | 0.984 | 0.157 | 266 | 99 | 9 | 9.12E-01 |
| *hcn4l* | 349 | 29 | 1 | 1 | 2 | 0.995 | 0.041 | 349 | 30 | 1 | 6.32E-01 |
| *neo1a* | 19 | 113 | 240 | 109 | 9 | 0.976 | 0.797 | 15 | 120 | 236 | 2.39E-01 |
| *neo1b* | 14 | 12 | 277 | 116 | 78 | 0.795 | 0.934 | 1 | 37 | 264 | 3.24E-32 |
| *quo* | 55 | 42 | 281 | 221 | 3 | 0.992 | 0.799 | 15 | 121 | 241 | 4.69E-37 |
| *si:dkey-65j6.2* | 22 | 60 | 295 | 66 | 4 | 0.990 | 0.862 | 7 | 90 | 280 | 1.34E-10 |

Number of embryos with 0, 1 and 2 mutated alleles; the number of embryos with nonsense mutations in both alleles (2\*); and the number of embryos with a missing call for that gene. P<sub>HWE</sub>: P-value for a Hardy-Weinberg equilibrium (HWE) exact test, considering a $\pm 30$ bp window around the CRISPR/Cas9-targeted site as a single locus ( $P < 2.9 \times 10^{-3}$ is significant after Bonferroni correction). In cases of deviation from HWE, the number of embryos with 2 mutated alleles is not lower than expected. Embryos with missing sequencing calls at more than two targeted orthologues were excluded from the statistical analysis (i.e. two embryos). For all genes, missed calls were normally distributed throughout the HRV and heart rate distributions, so for embryos with missed calls in at most two genes, we imputed the mean call for that gene, including for *neo1b*, where exclusion of either the gene or the embryos with a missed call was

**Supplementary Table 5.** Additive effects of CRISPR/Cas9-induced mutations on sinoatrial pauses and arrests

| Age | Outcome | n controls | n cases | Gene | OR | LCI | UCI | P |
| --- | --- | --- | --- | --- | --- | --- | --- | --- |
| 2dpf | SA pause | 258 | 39 | *gngt1* | 1.163 | 0.494 | 2.740 | 7.29E-01 |
|  |  |  |  | *syt10* | 1.467 | 0.810 | 2.654 | 2.06E-01 |
|  |  |  |  | *rgs6* | 0.788 | 0.175 | 3.553 | 7.57E-01 |
|  |  |  |  | *hcn4* | 2.749 | 1.461 | 5.171 | 1.71E-03 |
|  |  |  |  | *hcn4l* | omitted |  |  |  |
|  |  |  |  | *neo1a* | 1.176 | 0.507 | 2.731 | 7.06E-01 |
|  |  |  |  | *neo1b* | 2.445 | 0.658 | 9.080 | 1.82E-01 |
|  |  |  |  | *quo* | 0.746 | 0.401 | 1.389 | 3.56E-01 |
|  |  |  |  | *si:dkey-65j6.2* | 0.755 | 0.339 | 1.685 | 4.93E-01 |
|  |  |  |  | time of day | 1.147 | 0.856 | 1.539 | 3.59E-01 |
|  |  |  |  | intercept | 0.004 | 0.000 | 0.185 | 4.77E-03 |
|  |  |  |  | Batch 1 | 7.380 | 0.755 | 72.106 | 8.57E-02 |
|  |  |  |  | Batch 2 | 14.723 | 1.714 | 126.443 | 1.42E-02 |
|  |  |  |  | Batch 3 | 5.507 | 0.633 | 47.917 | 1.22E-01 |
|  |  |  |  | Batch 4 | 4.948 | 0.457 | 53.573 | 1.88E-01 |
|  |  |  |  | Batch 5 | 0.890 | 0.042 | 18.778 | 9.40E-01 |
| 2dpf | SA arrest | 212 | 36 | *gngt1* | 0.951 | 0.387 | 2.336 | 9.12E-01 |
|  |  |  |  | *syt10* | 1.447 | 0.792 | 2.642 | 2.30E-01 |
|  |  |  |  | *rgs6* | 0.855 | 0.193 | 3.799 | 8.37E-01 |
|  |  |  |  | *hcn4* | 2.644 | 1.402 | 4.986 | 2.66E-03 |
|  |  |  |  | *hcn4l* | omitted |  |  |  |
|  |  |  |  | *neo1a* | 1.165 | 0.499 | 2.717 | 7.24E-01 |
|  |  |  |  | *neo1b* | 1.694 | 0.434 | 6.604 | 4.48E-01 |
|  |  |  |  | *quo* | 0.764 | 0.404 | 1.442 | 4.06E-01 |
|  |  |  |  | *si:dkey-65j6.2* | 0.750 | 0.327 | 1.720 | 4.97E-01 |
|  |  |  |  | time of day | 1.117 | 0.829 | 1.506 | 4.66E-01 |
|  |  |  |  | intercept | 0.008 | 0.000 | 0.414 | 1.64E-02 |
|  |  |  |  | Batch 1 | 7.540 | 0.770 | 73.835 | 8.27E-02 |
|  |  |  |  | Batch 2 | 11.656 | 1.342 | 101.224 | 2.60E-02 |
|  |  |  |  | Batch 3 | 5.798 | 0.665 | 50.571 | 1.12E-01 |
|  |  |  |  | Batch 4 | 5.656 | 0.523 | 61.178 | 1.54E-01 |
|  |  |  |  | Batch 5 | omitted |  |  |  |
| 5dpf | SA pause | 312 | 9 | *gngt1* | 0.309 | 0.037 | 2.572 | 2.77E-01 |
|  |  |  |  | *syt10* | 0.236 | 0.050 | 1.128 | 7.05E-02 |
|  |  |  |  | *rgs6* | 1.049 | 0.068 | 16.223 | 9.73E-01 |
|  |  |  |  | *hcn4* | 2.460 | 0.680 | 8.899 | 1.70E-01 |
|  |  |  |  | *hcn4l* | 0.937 | 0.086 | 10.181 | 9.58E-01 |
|  |  |  |  | *neo1a* | 0.897 | 0.190 | 4.232 | 8.91E-01 |
|  |  |  |  | *neo1b* | 1.561 | 0.162 | 15.023 | 7.00E-01 |
|  |  |  |  | *quo* | 0.672 | 0.221 | 2.043 | 4.84E-01 |
|  |  |  |  | *si:dkey-65j6.2* | 3.533 | 0.421 | 29.662 | 2.45E-01 |
|  |  |  |  | time of day | 1.066 | 0.514 | 2.209 | 8.64E-01 |
|  |  |  |  | intercept | 0.016 | 0.000 | 7.788 | 1.90E-01 |
|  |  |  |  | Batch 1 | 0.240 | 0.011 | 5.447 | 3.70E-01 |
|  |  |  |  | Batch 2 | 0.703 | 0.073 | 6.750 | 7.60E-01 |
|  |  |  |  | Batch 3 | 0.214 | 0.015 | 3.049 | 2.55E-01 |
|  |  |  |  | Batch 5 | 0.213 | 0.005 | 8.956 | 4.18E-01 |

Associations of dichotomous cardiac outcomes with the number of mutated alleles across each of the nine orthologues, weighted by their predicted effect on protein function. At 2 and 5 days post fertilization (dpf), associations were analyzed using logistic regression for outcomes with at least 10 cases. Associations were adjusted for time of day and for the weighted number of mutated alleles in the other genes as fixed factors. The genes *quo* and *si:dkey-65j6.2* are orthologues of the human *KIAA1755*.

**Supplementary Table 6.**
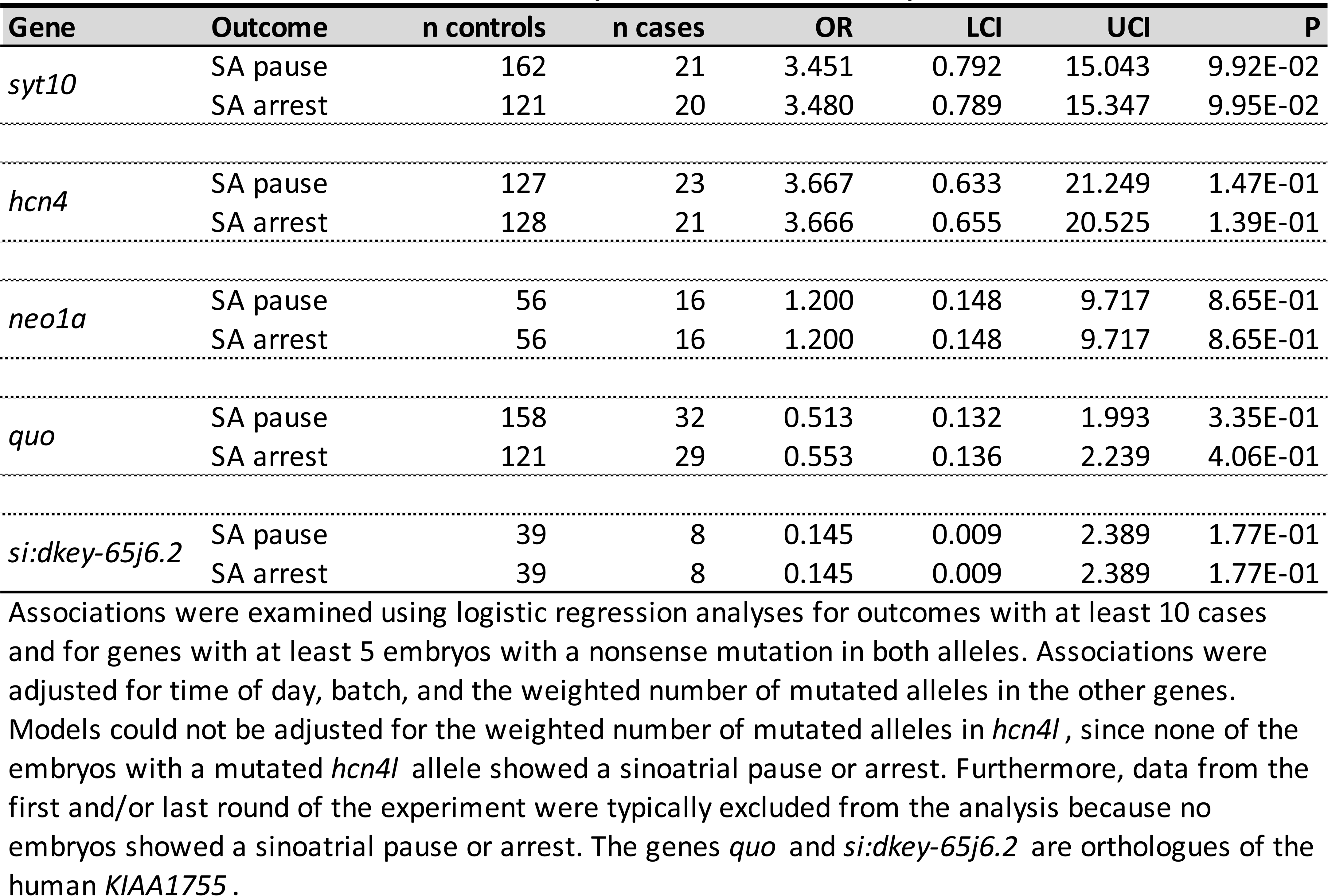
Effects of CRISPR/Cas9-induced nonsense mutations in both alleles vs. no CRISPR/Cas9-induced mutations on sinoatrial pauses and arrests at 2dpf

**Supplementary Table 7.**
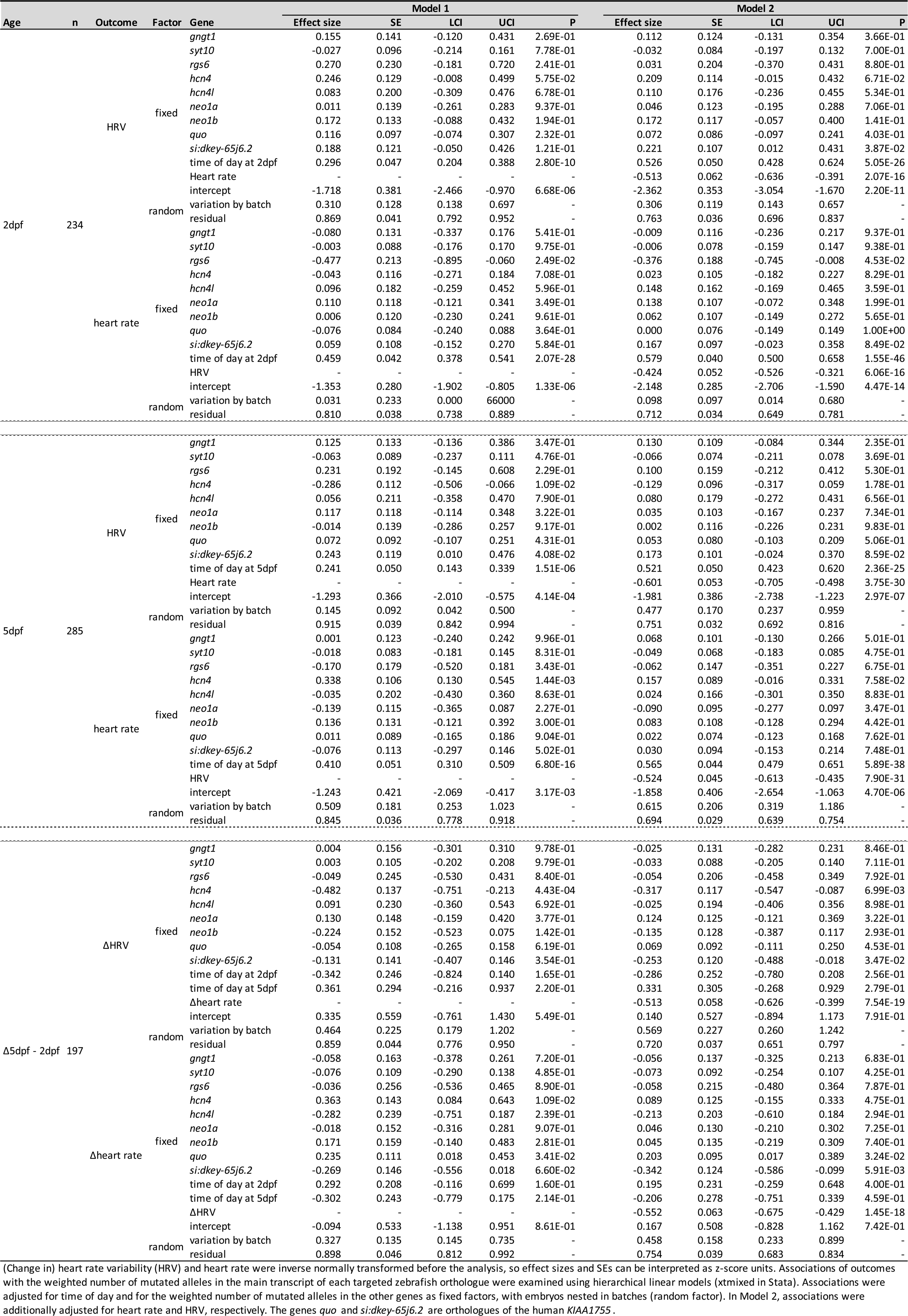
Additive effects of CRISPR/Cas9-induced mutations on (change in) heart rate variability and heart rate

**Supplementary Table 8.** Effects of CRISPR/Cas9-induced nonsense mutations in both alleles vs. no CRISPR/Cas9-induced mutations on (change in) heart rate variability and heart rate

| Gene | Age | n | Outcome | Model 1 |  |  |  |  | Model 2 |  |  |  |  |
| --- | --- | --- | --- | --- | --- | --- | --- | --- | --- | --- | --- | --- | --- |
|  |  |  |  | Effect size | SE | LCI | UCI | P | Effect size | SE | LCI | UCI | P |
| *syt10* | 2dpf | 146 | HRV | -0.073 | 0.205 | -0.475 | 0.329 | 7.22E-01 | -0.128 | 0.182 | -0.485 | 0.229 | 4.82E-01 |
|  |  |  | Heart rate | -0.115 | 0.199 | -0.506 | 0.276 | 5.64E-01 | -0.103 | 0.176 | -0.449 | 0.243 | 5.59E-01 |
|  | 5dpf | 180 | HRV | -0.066 | 0.191 | -0.440 | 0.308 | 7.30E-01 | -0.133 | 0.157 | -0.441 | 0.174 | 3.96E-01 |
|  |  |  | Heart rate | -0.184 | 0.189 | -0.555 | 0.187 | 3.31E-01 | -0.198 | 0.152 | -0.496 | 0.101 | 1.95E-01 |
| | Δ5dpf - 2dpf | 123 | ΔHRV | 0.014 | 0.198 | -0.375 | 0.403 | 9.43E-01 | -0.082 | 0.172 | -0.420 | 0.255 | 6.33E-01 |
| | | | Δheart rate | -0.228 | 0.225 | -0.668 | 0.213 | 3.12E-01 | -0.232 | 0.197 | -0.618 | 0.153 | 2.37E-01 |
| *hcn4* | 2dpf | 183 | HRV | 0.148 | 0.399 | -0.634 | 0.930 | 7.11E-01 | 0.520 | 0.344 | -0.153 | 1.194 | 1.30E-01 |
|  |  |  | Heart rate | 0.695 | 0.358 | -0.006 | 1.396 | 5.20E-02 | 0.674 | 0.309 | 0.068 | 1.280 | 2.93E-02 |
|  | 5dpf | 212 | HRV | -1.349 | 0.338 | -2.010 | -0.687 | 6.48E-05 | -0.759 | 0.294 | -1.335 | -0.182 | 9.91E-03 |
|  |  |  | Heart rate | 1.208 | 0.321 | 0.579 | 1.836 | 1.66E-04 | 0.444 | 0.278 | -0.100 | 0.988 | 1.10E-01 |
| | Δ5dpf - 2dpf | 152 | ΔHRV | -0.677 | 0.400 | -1.461 | 0.107 | 9.04E-02 | -0.350 | 0.344 | -1.024 | 0.324 | 3.09E-01 |
| | | | Δheart rate | 0.785 | 0.438 | -0.074 | 1.644 | 7.34E-02 | 0.373 | 0.378 | -0.368 | 1.113 | 3.24E-01 |
| *neo1a* | 2dpf | 70 | HRV | 0.029 | 0.343 | -0.643 | 0.702 | 9.32E-01 | 0.094 | 0.317 | -0.527 | 0.714 | 7.67E-01 |
|  |  |  | Heart rate | 0.102 | 0.294 | -0.473 | 0.677 | 7.28E-01 | 0.216 | 0.264 | -0.302 | 0.734 | 4.15E-01 |
|  | 5dpf | 93 | HRV | 0.089 | 0.286 | -0.473 | 0.650 | 7.57E-01 | 0.140 | 0.249 | -0.348 | 0.628 | 5.75E-01 |
|  |  |  | Heart rate | 0.166 | 0.274 | -0.372 | 0.704 | 5.45E-01 | 0.113 | 0.247 | -0.371 | 0.597 | 6.47E-01 |
| | Δ5dpf - 2dpf | 60 | ΔHRV | 0.192 | 0.392 | -0.575 | 0.960 | 6.23E-01 | 0.635 | 0.323 | 0.002 | 1.269 | 4.93E-02 |
| | | | Δheart rate | 0.702 | 0.342 | 0.032 | 1.372 | 4.02E-02 | 0.478 | 0.296 | -0.102 | 1.057 | 1.07E-01 |
| *neo1b* | 2dpf | 80 | HRV | 0.292 | 0.308 | -0.312 | 0.896 | 3.43E-01 | 0.332 | 0.290 | -0.236 | 0.901 | 2.52E-01 |
|  |  |  | Heart rate | 0.153 | 0.262 | -0.360 | 0.667 | 5.58E-01 | 0.225 | 0.243 | -0.251 | 0.701 | 3.55E-01 |
|  | 5dpf | 89 | HRV | 0.053 | 0.333 | -0.600 | 0.706 | 8.74E-01 | -0.092 | 0.289 | -0.658 | 0.474 | 7.50E-01 |
|  |  |  | Heart rate | 0.025 | 0.325 | -0.612 | 0.661 | 9.40E-01 | -0.073 | 0.276 | -0.614 | 0.469 | 7.92E-01 |
| | Δ5dpf - 2dpf | 63 | ΔHRV | -0.515 | 0.364 | -1.227 | 0.198 | 1.57E-01 | -0.555 | 0.308 | -1.159 | 0.049 | 7.17E-02 |
| | | | Δheart rate | 0.014 | 0.374 | -0.720 | 0.747 | 9.71E-01 | -0.277 | 0.331 | -0.925 | 0.372 | 4.03E-01 |
| *quo* | 2dpf | 170 | HRV | 0.227 | 0.206 | -0.176 | 0.631 | 2.70E-01 | 0.150 | 0.193 | -0.228 | 0.528 | 4.36E-01 |
|  |  |  | Heart rate | -0.104 | 0.178 | -0.454 | 0.245 | 5.58E-01 | 0.006 | 0.167 | -0.321 | 0.332 | 9.73E-01 |
|  | 5dpf | 214 | HRV | 0.240 | 0.193 | -0.139 | 0.618 | 2.14E-01 | 0.203 | 0.177 | -0.144 | 0.550 | 2.52E-01 |
|  |  |  | Heart rate | -0.065 | 0.192 | -0.441 | 0.310 | 7.34E-01 | 0.012 | 0.163 | -0.308 | 0.332 | 9.40E-01 |
| | Δ5dpf - 2dpf | 146 | ΔHRV | 0.045 | 0.250 | -0.445 | 0.535 | 8.57E-01 | 0.283 | 0.214 | -0.137 | 0.703 | 1.86E-01 |
| | | | Δheart rate | 0.424 | 0.236 | -0.039 | 0.887 | 7.27E-02 | 0.413 | 0.206 | 0.009 | 0.816 | 4.49E-02 |
| *si:dkey-65j6.2* | 2dpf | 60 | HRV | 0.291 | 0.298 | -0.294 | 0.876 | 3.29E-01 | 0.224 | 0.281 | -0.327 | 0.775 | 4.25E-01 |
|  |  |  | Heart rate | -0.110 | 0.226 | -0.552 | 0.332 | 6.27E-01 | 0.027 | 0.216 | -0.397 | 0.452 | 8.99E-01 |
|  | 5dpf | 67 | HRV | 0.364 | 0.350 | -0.322 | 1.051 | 2.98E-01 | 0.395 | 0.328 | -0.248 | 1.038 | 2.29E-01 |
|  |  |  | Heart rate | 0.111 | 0.298 | -0.473 | 0.695 | 7.08E-01 | 0.160 | 0.275 | -0.378 | 0.698 | 5.60E-01 |
| | Δ5dpf - 2dpf | 50 | ΔHRV | -0.648 | 0.425 | -1.480 | 0.184 | 1.27E-01 | -0.677 | 0.419 | -1.497 | 0.144 | 1.06E-01 |
| | | | Δheart rate | -0.160 | 0.412 | -0.967 | 0.647 | 6.97E-01 | -0.270 | 0.415 | -1.083 | 0.543 | 5.15E-01 |

(Change in) heart rate variability (HRV) and heart rate were inverse normally transformed before the analysis, so effect sizes and SEs can be interpreted as z-score units. Associations with outcomes are for embryos carrying frameshift and/or premature stop coding introducing mutations in both alleles vs. embryos free from CRISPR/Cas9-induced mutations. Associations were examined using hierarchical linear models (xtmixed in Stata) and were adjusted for time of day and for the weighted number of mutated alleles in the other genes as fixed factors, with larvae nested in batches (random factor). In Model 2, associations were additionally adjusted for (change in) heart rate and HRV, respectively. The genes *quo* and *si:dkey-65j6.2* are orthologues of the human *KIAA1755*.

**Supplementary Table 9.**
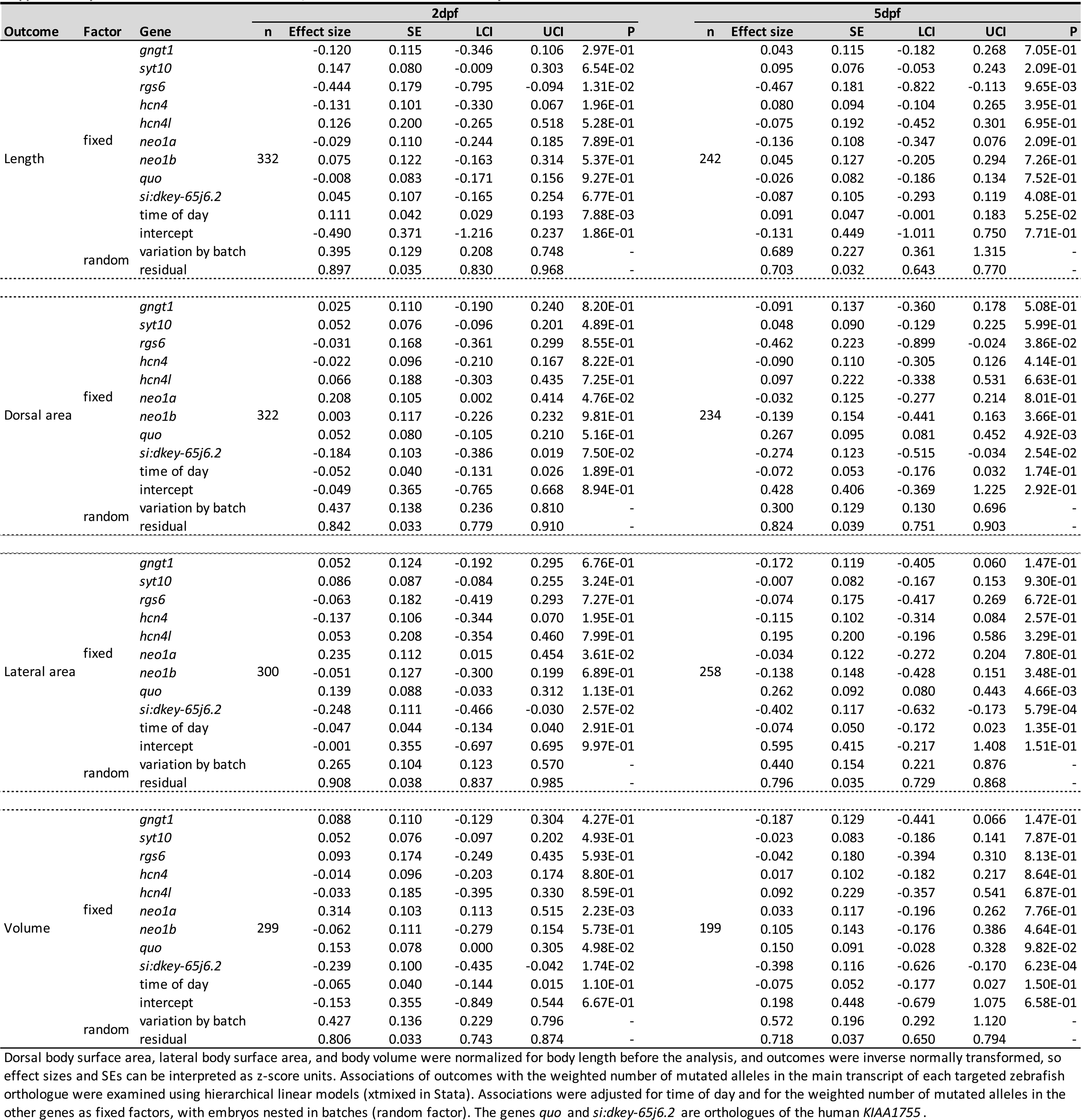
Additive effect of CRISPR/Cas9-induced mutations on body size

**Supplementary Table 10.**
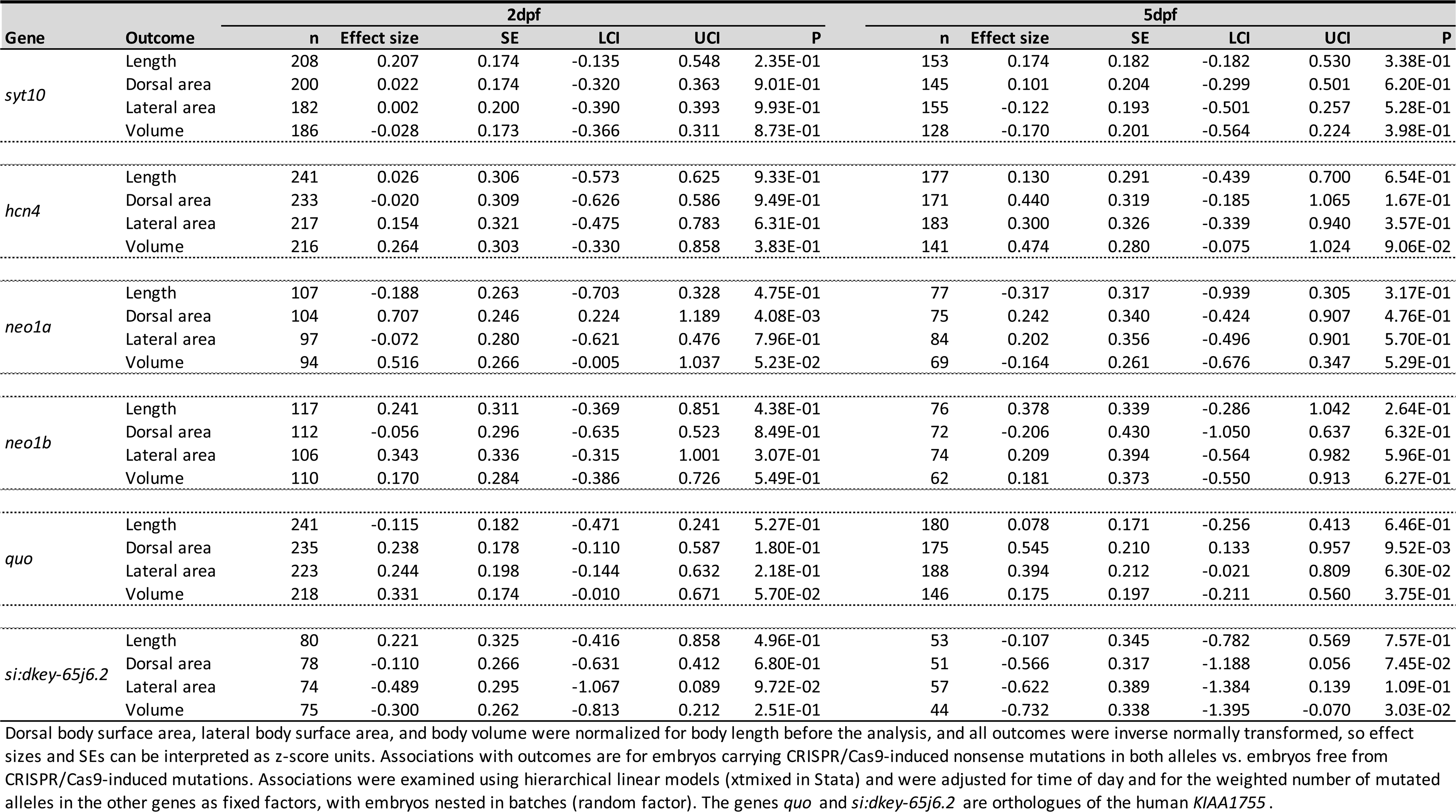
Effect of CRISPR/Cas9-induced nonsense mutations in both alleles vs. no CRISPR/Cas9-induced mutations on body size

| Gene | Outcome | 2dpf |  |  |  |  |  | 5dpf |  |  |  |  |  |
| --- | --- | --- | --- | --- | --- | --- | --- | --- | --- | --- | --- | --- | --- |
|  |  | n | Effect size | SE | LCI | UCI | P | n | Effect size | SE | LCI | UCI | P |
| *syt10* | Length | 208 | 0.207 | 0.174 | -0.135 | 0.548 | 2.35E-01 | 153 | 0.174 | 0.182 | -0.182 | 0.530 | 3.38E-01 |
|  | Dorsal area | 200 | 0.022 | 0.174 | -0.320 | 0.363 | 9.01E-01 | 145 | 0.101 | 0.204 | -0.299 | 0.501 | 6.20E-01 |
|  | Lateral area | 182 | 0.002 | 0.200 | -0.390 | 0.393 | 9.93E-01 | 155 | -0.122 | 0.193 | -0.501 | 0.257 | 5.28E-01 |
|  | Volume | 186 | -0.028 | 0.173 | -0.366 | 0.311 | 8.73E-01 | 128 | -0.170 | 0.201 | -0.564 | 0.224 | 3.98E-01 |
| *hcn4* | Length | 241 | 0.026 | 0.306 | -0.573 | 0.625 | 9.33E-01 | 177 | 0.130 | 0.291 | -0.439 | 0.700 | 6.54E-01 |
|  | Dorsal area | 233 | -0.020 | 0.309 | -0.626 | 0.586 | 9.49E-01 | 171 | 0.440 | 0.319 | -0.185 | 1.065 | 1.67E-01 |
|  | Lateral area | 217 | 0.154 | 0.321 | -0.475 | 0.783 | 6.31E-01 | 183 | 0.300 | 0.326 | -0.339 | 0.940 | 3.57E-01 |
|  | Volume | 216 | 0.264 | 0.303 | -0.330 | 0.858 | 3.83E-01 | 141 | 0.474 | 0.280 | -0.075 | 1.024 | 9.06E-02 |
| *neo1a* | Length | 107 | -0.188 | 0.263 | -0.703 | 0.328 | 4.75E-01 | 77 | -0.317 | 0.317 | -0.939 | 0.305 | 3.17E-01 |
|  | Dorsal area | 104 | 0.707 | 0.246 | 0.224 | 1.189 | 4.08E-03 | 75 | 0.242 | 0.340 | -0.424 | 0.907 | 4.76E-01 |
|  | Lateral area | 97 | -0.072 | 0.280 | -0.621 | 0.476 | 7.96E-01 | 84 | 0.202 | 0.356 | -0.496 | 0.901 | 5.70E-01 |
|  | Volume | 94 | 0.516 | 0.266 | -0.005 | 1.037 | 5.23E-02 | 69 | -0.164 | 0.261 | -0.676 | 0.347 | 5.29E-01 |
| *neo1b* | Length | 117 | 0.241 | 0.311 | -0.369 | 0.851 | 4.38E-01 | 76 | 0.378 | 0.339 | -0.286 | 1.042 | 2.64E-01 |
|  | Dorsal area | 112 | -0.056 | 0.296 | -0.635 | 0.523 | 8.49E-01 | 72 | -0.206 | 0.430 | -1.050 | 0.637 | 6.32E-01 |
|  | Lateral area | 106 | 0.343 | 0.336 | -0.315 | 1.001 | 3.07E-01 | 74 | 0.209 | 0.394 | -0.564 | 0.982 | 5.96E-01 |
|  | Volume | 110 | 0.170 | 0.284 | -0.386 | 0.726 | 5.49E-01 | 62 | 0.181 | 0.373 | -0.550 | 0.913 | 6.27E-01 |
| *quo* | Length | 241 | -0.115 | 0.182 | -0.471 | 0.241 | 5.27E-01 | 180 | 0.078 | 0.171 | -0.256 | 0.413 | 6.46E-01 |
|  | Dorsal area | 235 | 0.238 | 0.178 | -0.110 | 0.587 | 1.80E-01 | 175 | 0.545 | 0.210 | 0.133 | 0.957 | 9.52E-03 |
|  | Lateral area | 223 | 0.244 | 0.198 | -0.144 | 0.632 | 2.18E-01 | 188 | 0.394 | 0.212 | -0.021 | 0.809 | 6.30E-02 |
|  | Volume | 218 | 0.331 | 0.174 | -0.010 | 0.671 | 5.70E-02 | 146 | 0.175 | 0.197 | -0.211 | 0.560 | 3.75E-01 |
| *si:dkey-65j6.2* | Length | 80 | 0.221 | 0.325 | -0.416 | 0.858 | 4.96E-01 | 53 | -0.107 | 0.345 | -0.782 | 0.569 | 7.57E-01 |
|  | Dorsal area | 78 | -0.110 | 0.266 | -0.631 | 0.412 | 6.80E-01 | 51 | -0.566 | 0.317 | -1.188 | 0.056 | 7.45E-02 |
|  | Lateral area | 74 | -0.489 | 0.295 | -1.067 | 0.089 | 9.72E-02 | 57 | -0.622 | 0.389 | -1.384 | 0.139 | 1.09E-01 |
|  | Volume | 75 | -0.300 | 0.262 | -0.813 | 0.212 | 2.51E-01 | 44 | -0.732 | 0.338 | -1.395 | -0.070 | 3.03E-02 |

Dorsal body surface area, lateral body surface area, and body volume were normalized for body length before the analysis, and all outcomes were inverse normally transformed, so effect sizes and SEs can be interpreted as z-score units. Associations with outcomes are for embryos carrying CRISPR/Cas9-induced nonsense mutations in both alleles vs. embryos free from CRISPR/Cas9-induced mutations. Associations were examined using hierarchical linear models (xtmixed in Stata) and were adjusted for time of day and for the weighted number of mutated alleles in the other genes as fixed factors, with embryos nested in batches (random factor). The genes *quo* and *si:dkey-65j6.2* are orthologues of the human *KIAA1755*.

**Supplementary Table 11.** Transcripts with at least 75% sequence similarity to the zebrafish hcn4 cDNA sequence

| Gene | ENSDARG stable ID | BLAST Score | %ID | Human orthologue | ENSG stable ID | Target %id | Query %id |
| --- | --- | --- | --- | --- | --- | --- | --- |
| *hcn4l* | ENSDARG00000074419 | 479 | 85.20 | *HCN4* | ENSG00000138622 | 59.96 | 50.04 |
| *CABZ01086574.1* | ENSDARG00000116404 | 277 | 80.09 | *HCN2* | ENSG00000099822 | 60.30 | 22.72 |
| *hcn2b* | ENSDARG00000061665 | 249 | 85.63 | *HCN2* | ENSG00000099822 | - | - |
| *hcn3* | ENSDARG00000027192 | 200 | 82.35 | *HCN3* | ENSG00000143630 | - | - |
| *hcn1* | ENSDARG00000104480 | 160 | 78.42 | *HCN1* | ENSG00000164588 | 72.25 | 73.71 |

Zebrafish genes with transcripts showing at least 75% sequence similarity to the main zebrafish *hcn4* transcript were selected for a qRT-PCR experiment to explore possible compensatory responses to CRISPR/Cas9 mutagenesis of *hcn4*. BLAST score represents the highest score for a transcript of that zebrafish gene against the *hcn4* cDNA sequence. 'Human orthologue' shows for which human gene that zebrafish gene was flagged as an orthologue in Ensembl. Target %id is the percentage of the orthologous sequence that matches the human sequence (Ensembl); Query %id is the percentage of the human sequence matching the sequence of the orthologue.

**Supplementary Table 12.**
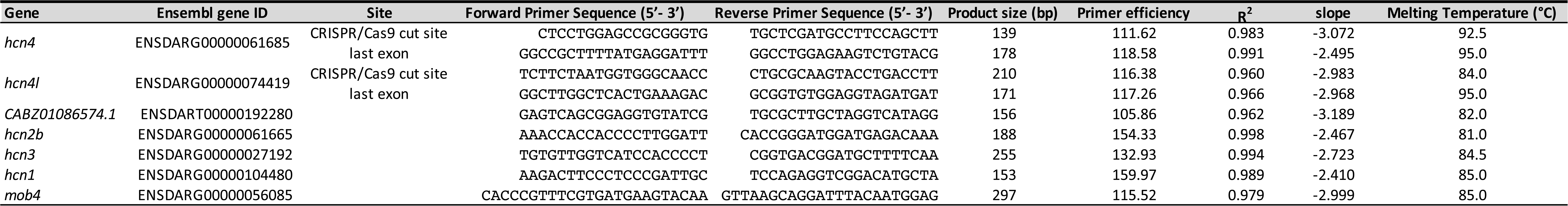
qRT-PCR target sites, primers and experimental conditions

**Supplementary Table 13.**
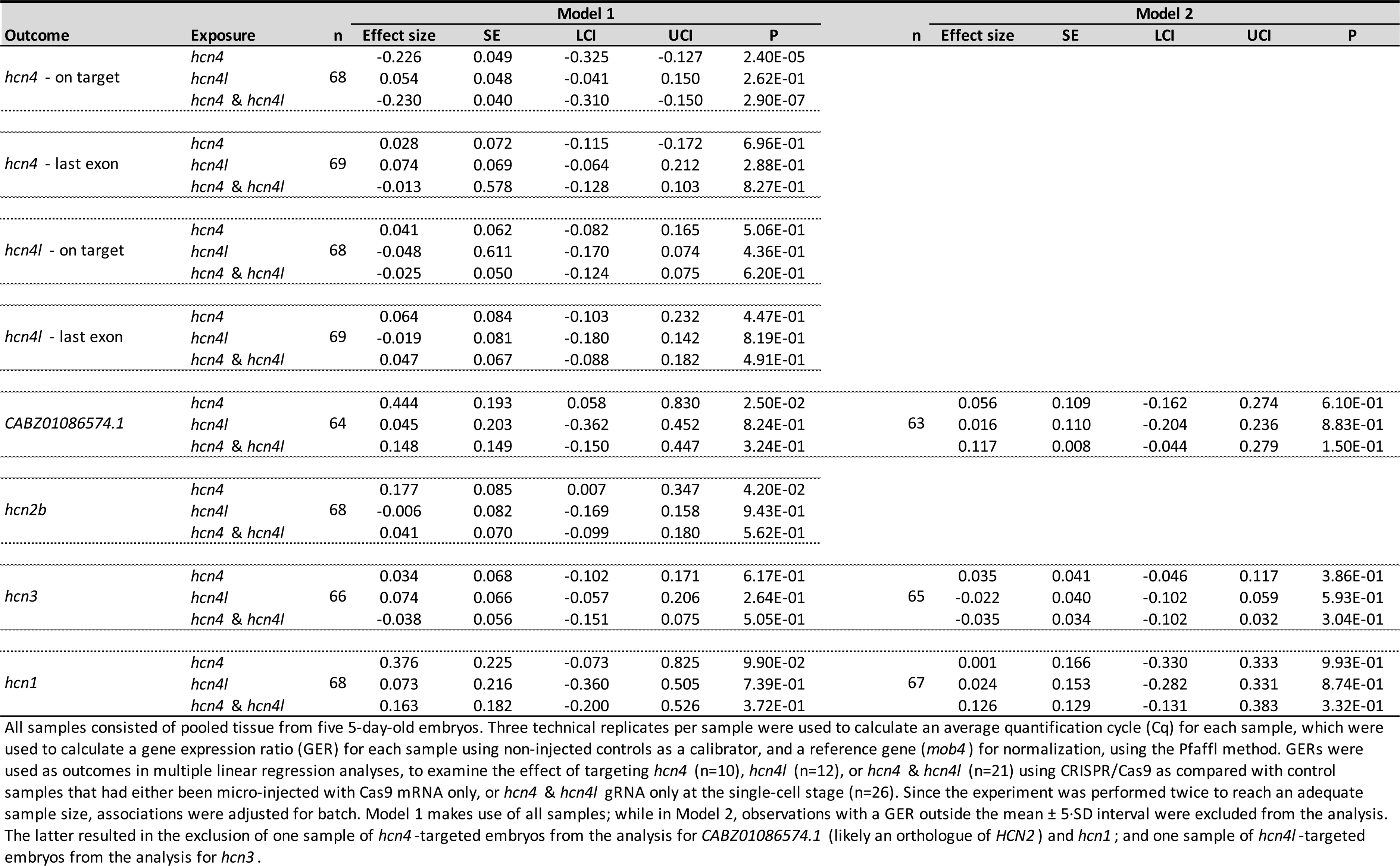
Effects of targeting hcn4, hcn4l, or hcn4 and hcn4l using CRISPR/Cas9 on the expression of genes with >75% sequence similarity to the main zebrafish hcn4 transcript

**Supplementary Table 14.**
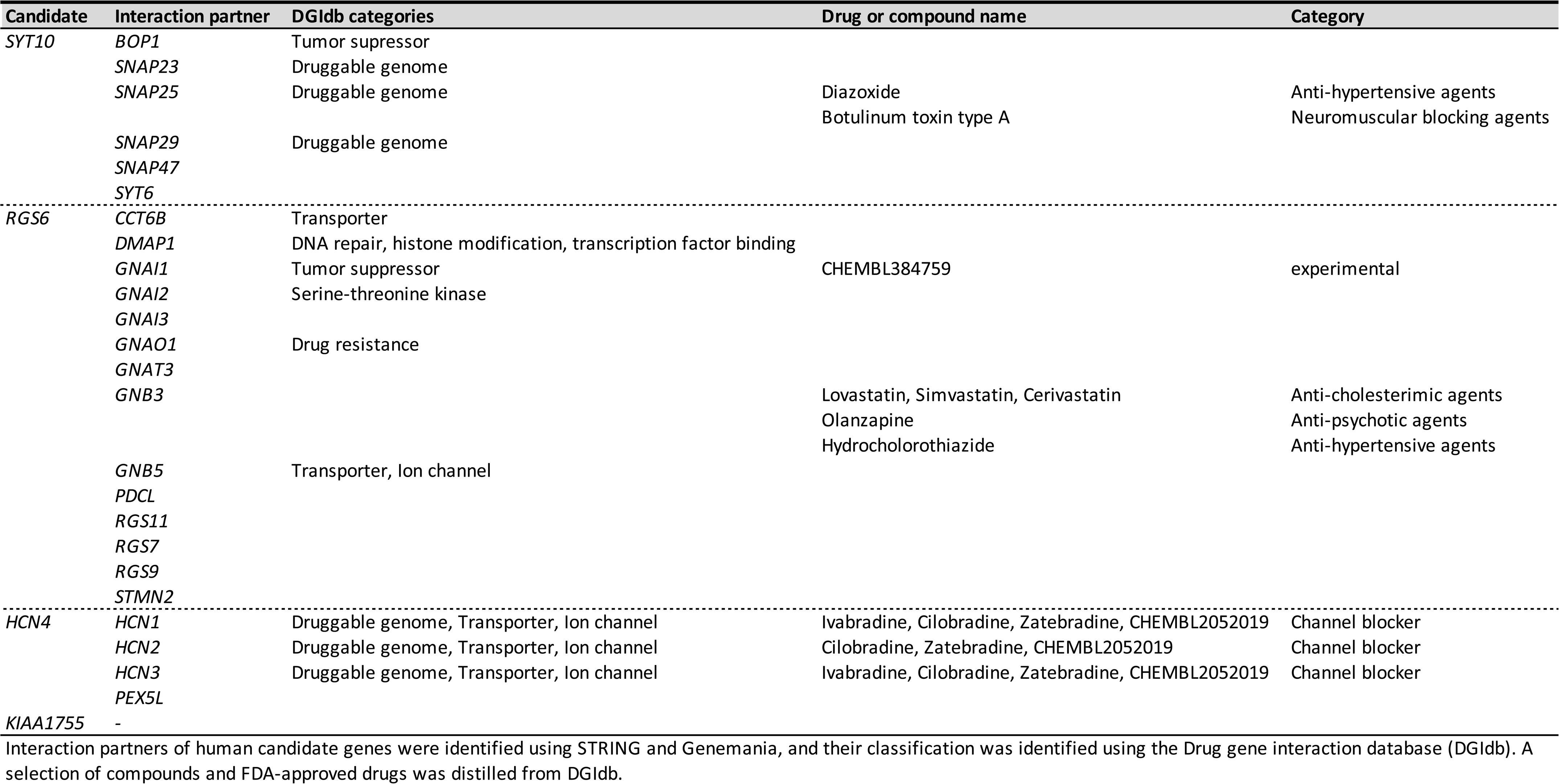
Druggability of interacting partners of putative causal genes

**Supplementary Table 15.** Effect of 24h of ivabradine treatment on heart rate variability and heart rate in 5dpf zebrafish embryos

| Outcome | Factor | Exposure | n | Model 1 |  |  |  |  | n | Model 2 |  |  |  |  |
| --- | --- | --- | --- | --- | --- | --- | --- | --- | --- | --- | --- | --- | --- | --- |
|  |  |  |  | Effect size | SE | LCI | UCI | P |  | Effect size | SE | LCI | UCI | P |
| HRV | fixed | ivabradine (10 μM) | 109 | 0.328 | 0.179 | -0.022 | 0.678 | 6.65E-02 | 109 | 0.051 | 0.165 | -0.273 | 0.374 | 7.58E-01 |
| | | ivabradine (25 μM) | | 0.905 | 0.172 | 0.568 | 1.242 | 1.45E-07 | | 0.258 | 0.190 | -0.115 | 0.631 | 1.76E-01 |
|  |  | heart rate (in SD) |  | - | - | - | - | - |  | -0.523 | 0.093 | -0.706 | -0.340 | 2.02E-08 |
|  |  | time of day |  | -0.057 | 0.071 | -0.197 | 0.083 | 4.24E-01 |  | 0.105 | 0.069 | -0.031 | 0.240 | 1.31E-01 |
|  |  | intercept |  | -0.541 | 0.283 | -1.096 | 0.015 | 5.64E-02 |  | -0.783 | 0.253 | -1.279 | -0.286 | 2.00E-03 |
|  | random | variation by batch |  | - | 0.000 | 0.000 | 0.001 | - |  | - | 0.000 | 0.000 | 1.009 | - |
|  |  | residual |  | - | 0.051 | 0.660 | 0.860 | - |  | - | 0.045 | 0.581 | 0.758 | - |
| Heart rate | fixed | ivabradine (10 μM) | 118 | -0.524 | 0.158 | -0.835 | -0.214 | 9.35E-04 | 109 | -0.401 | 0.142 | -0.678 | -0.123 | 4.68E-03 |
| | | ivabradine (25 μM) | | -1.373 | 0.154 | -1.675 | -1.071 | 5.29E-19 | | -0.872 | 0.151 | -1.169 | -0.576 | 8.07E-09 |
|  |  | HRV (in SD) |  | - | - | - | - | - |  | -0.415 | 0.075 | -0.563 | -0.267 | 3.63E-08 |
|  |  | time of day |  | 0.251 | 0.072 | 0.110 | 0.392 | 4.69E-04 |  | 0.274 | 0.061 | 0.154 | 0.395 | 8.28E-06 |
|  |  | intercept |  | -0.251 | 0.301 | -0.841 | 0.338 | 4.04E-01 |  | -0.655 | 0.246 | -1.137 | -0.173 | 7.76E-03 |
|  | random | variation by batch |  | - | 0.113 | 0.114 | 0.611 | - |  | - | 0.089 | 0.038 | 0.488 | - |
|  |  | residual |  | - | 0.046 | 0.611 | 0.794 | - |  | - | 0.041 | 0.512 | 0.672 | - |

Heart rate variability (HRV) and heart rate were inverse normally transformed before the analysis, so effect sizes and SEs can be interpreted as z-score units. Associations of outcomes with exposure to 10 or 25 μM ivabradine in DMSO vs. DMSO only were examined using hierarchical linear models (xtmixed in Stata). Associations were adjusted for time of day as fixed factors, with embryos nested in batches (random factor). In Model 2, associations were additionally adjusted for heart rate and HRV, respectively.

## Notes

### Competing Interest Statement

The authors have declared no competing interest.

### Summary of Updates

We performed a range of additional experiments and analyses, i.e.: - we examined the effect of 1 day of treatment with ivabradine on heart rate variability and heart rate in 5 day-old zebrafish embryos (lower heart rate; higher heart rate variability, as in humans). - we performed additional analyses to make better use of our longitudinal data (observed genetic effects on change over time). - we added results from a screen in 406 embryos with 0, 1, or 2 nonsense mutations in hcn4 (sa11188) on heart rate (consistent with results CRISPR screen). - we performed a qRT-PCR experiment to examine if upregulation of expression of transcripts with >75% sequence similarity to hcn4 may have influenced our results (affirmative). - we performed a genome-wide off target screen to examine if off-target activity of the CRISPR/Cas9 gRNAs for hcn4, hcn4l, neo1a and/or neo1b may have influenced our results (negative)

